# TAD hierarchy restricts poised LTR activation and loss of TAD hierarchy promotes LTR co-option in cancer

**DOI:** 10.1101/2024.05.31.596845

**Authors:** Elissa W. P. Wong, Merve Sahin, Rui Yang, UkJin Lee, Yingqian A. Zhan, Rohan Misra, Fanny Tomas, Nawaf Alomran, Alexander Polyzos, Cindy J. Lee, Tuan Trieu, Alexander M. Fundichely, Thomas Wiesner, Andrew Rosowicz, Shuyuan Cheng, Christina Liu, Morgan Lallo, Taha Merghoub, Pierre-Jacques Hamard, Richard Koche, Ekta Khurana, Effie Apostolou, Deyou Zheng, Yu Chen, Christina S. Leslie, Ping Chi

**Affiliations:** Human Oncology and Pathogenesis Program, Memorial Sloan Kettering Cancer Center, New York, NY, USA; Computational and Systems Biology Program, Memorial Sloan Kettering Cancer Center, New York, NY, USA; Tri-Institutional Training Program in Computational Biology and Medicine, New York, NY, USA; Weill Cornell Graduate School of Medical Science, Weill Cornell Medicine, New York, NY, USA; Center for Epigenetics Research, Memorial Sloan Kettering Cancer Center, New York, NY, USA; Department of Genetics, Albert Einstein College of Medicine, Bronx, NY, USA; Sandra and Edward Meyer Cancer Center, Weill Cornell Medicine, New York, NY, USA; Institute for Computational Biomedicine, Weill Cornell Medicine, New York, NY, USA; Department of Physiology and Biophysics, Weill Cornell Medicine, New York, NY, USA; Englander Institute for Precision Medicine, Weill Cornell Medicine, New York, NY, USA; Department of Dermatology, Medical University of Vienna, Vienna, Austria. Current address; Department of Pathology, Medical University of Vienna, Vienna, Austria. Current address; Icahn School of Medicine at Mount Sinai, New York, NY, USA. Current address; Molecular Pharmacology Program, Sloan Kettering Institute, Memorial Sloan Kettering Cancer Center, New York, NY, USA; Yale University School of Medicine, New Haven, CT, USA; Parker Institute for Cancer Immunotherapy, Memorial Sloan Kettering Cancer Center, New York, NY, USA; Department of Pharmacology and Edward Meyer Cancer Center and Ludwig Collaborative and Swim Across America Laboratory, Weill Cornell Medicine, New York, NY, USA. Current address; Sanford I. Weill Department of Medicine, Weill Cornell Medical College, New York, NY, USA. Current address; Sanford I. Weill Department of Medicine, Weill Cornell Medical College, New York, NY, USA; The Saul R. Korey Department of Neurology, Albert Einstein College of Medicine, Bronx, NY, USA; Dominick P. Purpura Department of Neuroscience, Albert Einstein College of Medicine, Bronx, NY, USA; Louis V. Gerstner, Jr. Graduate School of Biomedical Sciences, Memorial Sloan Kettering Cancer Center, New York, NY, USA; Department of Medicine, Memorial Sloan Kettering Cancer Center, New York, NY, USA

## Abstract

Transposable elements (TEs) are abundant in the human genome, and they provide the sources for genetic and functional diversity. The regulation of TEs expression and their functional consequences in physiological conditions and cancer development remain to be fully elucidated. Previous studies suggested TEs are repressed by DNA methylation and chromatin modifications. The effect of 3D chromatin topology on TE regulation remains elusive. Here, by integrating transcriptome and 3D genome architecture studies, we showed that haploinsufficient loss of *NIPBL* selectively activates alternative promoters at the long terminal repeats (LTRs) of the TE subclasses. This activation occurs through the reorganization of topologically associating domain (TAD) hierarchical structures and recruitment of proximal enhancers. These observations indicate that TAD hierarchy restricts transcriptional activation of LTRs that already possess open chromatin features. In cancer, perturbation of the hierarchical chromatin topology can lead to co-option of LTRs as functional alternative promoters in a context-dependent manner and drive aberrant transcriptional activation of novel oncogenes and other divergent transcripts. These data uncovered a new layer of regulatory mechanism of TE expression beyond DNA and chromatin modification in human genome. They also posit the TAD hierarchy dysregulation as a novel mechanism for alternative promoter-mediated oncogene activation and transcriptional diversity in cancer, which may be exploited therapeutically.

## Introduction

Approximately 50% of the mammalian genome consists of transposable elements (TEs), mainly retrotransposons, that once possessed the ability to move their locations in the genome^1,2^. TEs have actively shaped genomic functions through multitude of mechanisms, including genome structural rearrangements, introduction of mutations and transcriptional regulation^1,3^. In turn, the mammalian genome has evolved to take advantage of this abundant source of genetic material to regulate essential cellular functions during early stem cell and embryonic development^4^, as well as diversification of neuronal cells^5,6^. The intricate balance between the often-detrimental consequences of unchecked TE expression and occasional beneficial exaptation of TE is essential to both the survival of the TEs and host genome^4,7^. In mammals, several mechanisms have been described to tightly control the expression of TEs spatially and temporally including non-coding RNAs, DNA methylation and histone modification^4,8–10^. The interplay of DNA methylation and histone modification to safeguard TE repression is best exemplified in embryonic stem cells and primordial germ cells where global DNA demethylation overlaps with decoration of repressive histone marks to prevent aberrant transcription from TEs^11,12^. De-repression or insertion of TEs has been linked to cancer where TEs were either employed as alternative promoters for oncogene activation or as disruptors of tumor suppressors^7,8,13^. Recent studies suggest that TE-driven oncogene expression, a process known as onco-exaptation, is frequent across multiple cancer types^14^. Among TEs, retrotransposons and in particularly long terminal repeats (LTRs) that contain intact RNA polymerase II binding sites, transcription factor binding sites (TFBS) and in some cases, conserved splice donor sites, render LTRs with a higher propensity to be co-opted as alternative promoters than other classes of TEs such as LINE and SINE^7,15^. However, the regulatory mechanisms of alternative promoter usage through TE co-option in cancer remain poorly understood.

It is well established that repetitive DNA elements including TEs can be transcriptionally repressed by constitutive heterochromatin characterized by H3K9me3 modification and DNA methylation^16^. The H3K27me3 repressive chromatin mark also suppresses TE expression in a cell-type-specific manner^9,11,12,16^.Chromatin is further organized into 3D structures, consisting of self-interacting topologically associating domains (TADs). TADs are established and maintained through CTCF/cohesin-mediated looping and regulate long-range interactions between enhancers and promoters and consequent transcriptional activation^17,18^. TADs can be further organized into a hierarchical structure of nested subTADs, i.e., hierarchical TADs^19^. Compared to non-hierarchical TADs, hierarchical TADs are associated with functionally more active epigenetic states and a higher level of active gene transcription^19^.

Although TE has recently been described to modulate CTCF binding and influence TAD boundary^20,21^, whether and how TADs regulate TE expression remains largely unknown. Genomic regions within TADs are thought to interact more frequently within TAD boundaries defined by convergently oriented CTCF and ring-structured protein complex cohesin through loop extrusion. Unlike CTCF, the cohesin complex does not recognize specific DNA sequences and its residence on chromatin relies on the continuous loading action by the cohesin complex loading factors, Nipped-B-like protein (NIPBL) and MAU2^17^. NIPBL is essential in the establishment and maintenance of TADs^22^. Complete loss of NIPBL is lethal in dividing cells^22^. Pathologically, heterozygous loss-of-function (LoF) *NIPBL* mutations underline the molecular basis of the majority of Cornelia de Lange syndrome (CdLS). CdLS is a multisystem developmental disorder caused by genetic aberrations of the core structural or regulatory components (e.g., *NIPBL, SMC1A, SMC3*) of the cohesin complex. *NIPBL* mutations are typically linked to the most severe clinical presentation of the disease^23^. Further, monoallelic LoF mutations of *NIPBL* are frequently observed in multiple cancers, and *NIPBL* is recognized as a *bona fide* tumor suppressor in pan-cancer analyses^24^.

Here, we used a doxycycline-inducible system to mimic the monoallelic loss of *NIPBL* in disease and evaluated its impact on chromatin topology and transcriptional regulation in cancer. We found that haploinsufficient loss of *NIPBL* led to a reduction of hierarchical TADs and global transcriptional activation of alternative promoters. We observed that these alternative promoters arose primarily from LTR elements that were centrally located in the nested hierarchical TADs and already possessed open chromatin features. We further demonstrated that the reorganization of hierarchical TADs and subsequent recruitment and retargeting of proximal enhancers contributed to the activation of alternative promoters from LTRs. These data indicate that chromatin topological hierarchy that is maintained by NIPBL restricts transcriptional activation from LTRs, and perturbation of the hierarchical chromatin topology in cancer can lead to the co-option of LTRs and aberrant transcriptional activation of novel oncogenes and divergent transcripts.

## Results

### Haploinsufficient loss of *NIPBL* induces global alternative promoter activation

To mimic the monoallelic loss of *NIPBL* in cancer and CdLS, we used a doxycycline-inducible RNA interference (RNAi) system, shNIPBL_#1 and shNIPBL_#2 that downregulated the *NIPBL* RNA transcript levels by approximately 34% and 56%, respectively, in melanoma cells (**Extended Data Fig. 1a**). *NIPBL* downregulation in 501mel melanoma cells did not significantly alter the total protein level of SMC1A, a core component of the cohesin complex^22^ (**Extended Data Fig. 1b**); genome-wide binding of SMC1A protein on chromatin by chromatin immunoprecipitation coupled with next generation sequencing (ChIP-seq) showed a modest reduction, indicating a partial loss of cohesin loading as a result of the *NIPBL* downregulation (**Extended Data Fig. 1c**). To investigate the effects of haploinsufficient *NIPBL* loss on transcriptome and transcriptional start site (TSS) usage, we performed poly-A RNA-sequencing (RNA-seq) to capture most mature RNA transcripts and cap-analysis gene expression sequencing (CAGE-seq) to accurately assess TSS, including alternative promoters^25^, in 501mel melanoma cells. Downregulation of *NIPBL* resulted in significant transcriptome changes in a subset of genes (*FDR*<0.05 and log2FoldChange >1). Greater downregulation of *NIPBL* was associated with more transcriptome changes, e.g., shNIPBL_#1 (34% reduction in *NIPBL* transcript) resulted in 311/178 significantly upregulated/downregulated genes and shNIPBL_#2 (56% reduction in *NIPBL* transcript) resulted in 1329/1358 significantly upregulated/downregulated genes (**Fig. 1a and Supplementary Table 1**). CAGE-seq revealed a comparable number of CAGE-TSS (CTSS) changes as poly-A RNA-seq with *NIPBL* perturbation; there were more upregulated than downregulated CTSSs by both *NIPBL-*specific shRNAs (**Fig. 1b** and **Supplementary Table 2**). The CAGE-seq CTSSs correlated well with RNA-seq-detected TSSs for annotated transcripts (**Extended Data Fig. 2a-c**). Transcriptome changes detected by RNA-seq were concordant with the CTSS changes detected by CAGE-seq in both shNIPBL hairpins when compared to shLuc control (**Fig. 1c**). At baseline, approximately 56% of all CTSSs were located at annotated promoters; about 16%, 10%, 9%, and 6% were localized to annotated exons, introns, distal intergenic regions and 3’ UTRs, respectively (**Fig. 1d** and **Supplementary Table 2**). Notably, the shNIPBL-upregulated CTSSs were disproportionally located in introns (10.4% [baseline] vs 33.7% [shNIPBL_#1] and 22.2% [shNIPBL_#2]), distal intergenic regions (9.0% [baseline] vs. 35.0% [shNIPBL_#1] and 19.2% [shNIPBL_#2]), and less frequently at promoters (56.2% [baseline] vs 22.0% [shNIPBL_#1] and 50.7% [shNIPBL_#2]) (**Fig. 1d, e**), indicating alternative promoter usage. The downregulated CTSSs by shNIPBL_#1 were preferentially at promoters (56.2% [baseline] vs 92.1% [shNIPBL_#1] and 57.5% [shNIPBL_#2]) (**Fig. 1d, e**). Manual examination of the RNA-seq and CAGE-seq profiles for the top 30% significantly upregulated CTSSs in introns (25 out of 83 with shNIPBL_#1, and 54 out of 178 for shNIPBL_#2) confirmed that all these regions represented functional alternative promoters and produced mature spliced transcripts. Similarly, examination of the top 30% significantly upregulated CTSSs in distal intergenic regions (26 out of 86 with shNIPBL_#1 and 47 out of 155 for shNIPBL_#2) found that 81% and 96% of these regions accounted for alternative promoters with mature spliced transcripts, respectively. For example, the *ALK*^ATI^ arose from an alternative promoter in intron 19 and subsequently spliced into exons 20-29 of *ALK* expressing the entire kinase domain, which has been previously characterized as an oncogenic variant of *ALK* prevalent in melanoma and sporadically in other cancer types^26^ (**Fig. 1a, b, f**). Similarly, the *ULK4* variant arose from intron 31 and *LINC01387* from intron 2, both of which spliced into existing subsequent exons to produce mature transcripts (**Fig. 1a, b, f**). Moreover, we observed that some RNA transcripts originated from distal intergenic regions, such as between *SYN3* and *LINC01640* in chr22:33,468,501-33,471,517 and between *TAS2R39* and *TAS2R40* in chr7:142,895,998-142,896,596, respectively (**Fig. 1f**). The distal intergenic alternative promoter in chr7:142,895,998-142,896,596 revealed a distinctive feature where transcription was initiated, and the transcript was spliced across a genomic distance of ∼240kb into the coding gene *KEL* (**Fig. 1f**).

**Figure 1.**
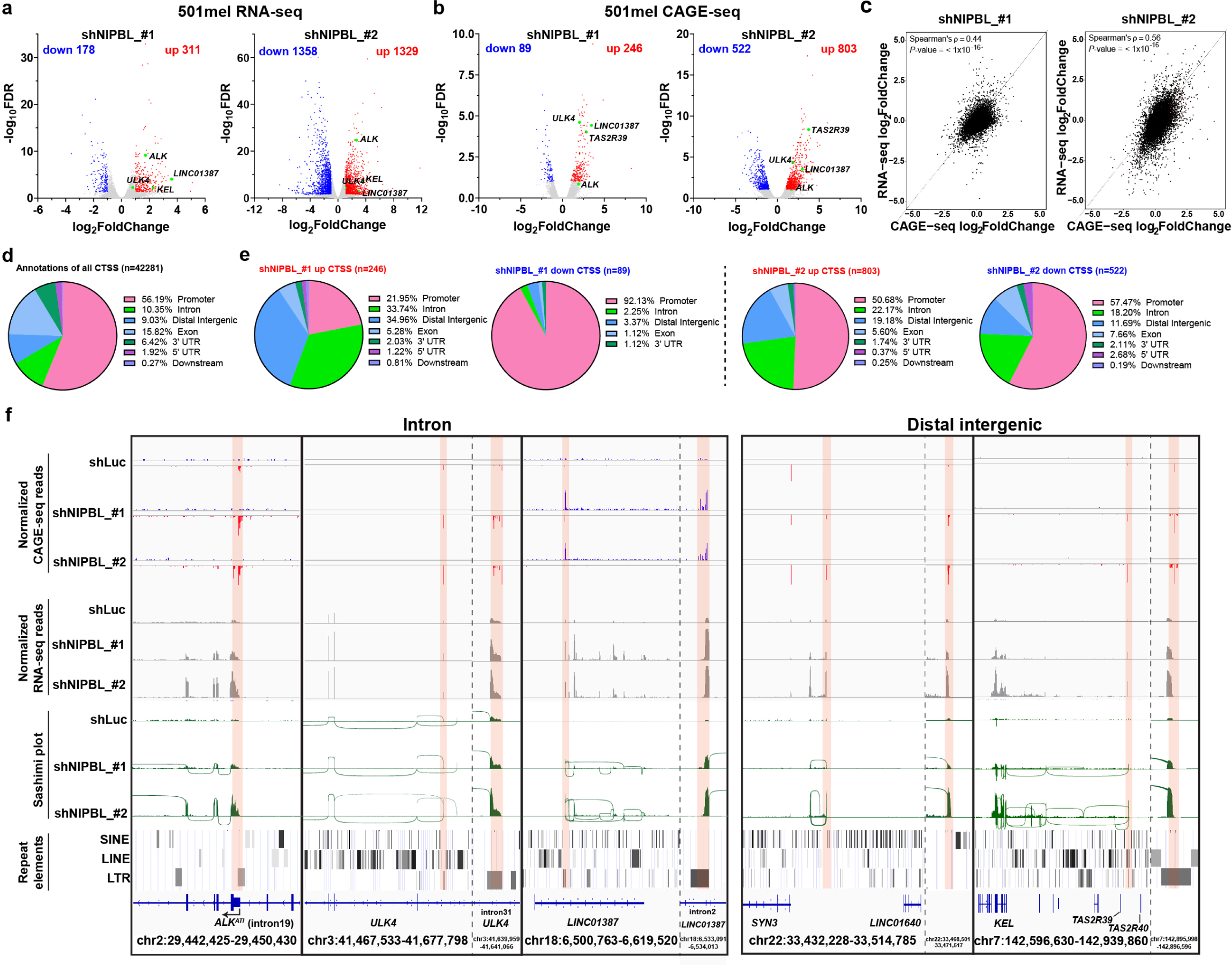
Haploinsufficient loss of *NIPBL* leads to alternative promoter activation in intron and distal intergenic regions. **a,** Volcano plots of differentially regulated genes (FDR<0.05) by whole transcriptome analysis of shRNA-mediated *NIPBL* perturbation in 501mel melanoma cells. Upregulated genes (red, log2FoldChange>1), downregulated genes (blue, log2FoldChange<-1) and genes of interest (green). n=2 biological replicates. **b,** Volcano plots of significantly altered transcriptional start sites (CTSSs, FDR<0.1, log2FoldChange>1) by CAGE-seq under *NIPBL* perturbation conditions as in **a.** n=2 biological replicates. **c,** Spearman correlation of transcriptome changes (log2FoldChange) of annotated transcripts between whole transcriptome analysis by poly-A RNA-seq and CTSS analysis by CAGE-seq under *NIPBL* perturbation conditions (*P*-value<1×10^−16^ for both shNIPBL conditions). **d,** Genomic annotation of all CTSSs by CAGE-seq under control condition. Percentage of each genomic feature is indicated. **e,** Distribution of genomic features of significantly (FDR<0.1) upregulated (log2FoldChange>1) and downregulated (log2FoldChange<-1) CTSSs with shRNA-mediated *NIPBL* downregulation. **f,** Representative examples of CAGE-seq and RNA-seq profiles of significantly upregulated CTSSs in intron and distal intergenic regions with *NIPBL* downregulation. Normalized CAGE-seq (blue, plus strand; red, minus strand) and RNA-seq profiles from two independent experiments are shown. Sashimi plots from one representative RNA-seq experiment depicted novel spliced transcripts initiated from respective CTSSs shaded in pink. The presence of repetitive elements is indicated by the “repeatmasker” track from the UCSC genome browser (hg19 [GRCh37] genomic version). Enlarged genomic regions are shown in dotted inset.

We employed the doxycycline-inducible RNAi system in two additional melanoma cell lines, COLO800 and A375, and analyzed their transcriptome changes by RNA-seq. We observed similar alternative promoter usages in *ALK* (intron 19), *LINC01387* (intron 2), *SYN3* and *KEL* (intergenic regions) (**Extended Data Fig. 1a, 3a-e**). In addition, we performed CRISPR interference (CRISPRi)^27^ for targeted silencing of *NIPBL* promoter in 501mel cells (**Extended Data Fig. 4a**). Consistent with the RNAi system, CRISPRi-mediated downregulation of *NIPBL* resulted in ∼50% decrease in *NIPBL* mRNA levels and significant increase in *ALK*^ATI^ RNA transcripts originating from the alternative promoter in *ALK* intron 19 (**Extended Data Fig. 4b**). These observations are reminiscent of a previous study where *Nipbl* knockout in non-dividing murine hepatocytes was associated with increased intergenic and antisense (exogenic) transcripton^22^, and demonstrated that the novel CTSSs detected by CAGE-seq were associated with production of mature transcripts.

To determine whether *NIPBL* loss leads to alternative promoter usage in cancer types other than melanoma, we performed CAGE-seq in nine representative cell lines from four major cancer types, including non-small cell lung cancer (NSCLC), colorectal cancer, hormone receptor-positive prostate and breast cancers, after downregulation of *NIPBL* by doxycycline-induced shNIPBL_#2 (**Extended Data Fig. 5a-d and Supplementary Table 3**). shNIPBL_#2 hairpin led to 42-73% reduction in *NIPBL* mRNA when compared to shLuc control in the cell lines used (**Extended Data Fig. 5a-d**). At baseline, we found similar distribution of all CTSSs that were located at annotated promoters (47-53%) in all nine cancer cell lines as in 501mel melanoma cells (56%) (**Fig. 1d, Extended Data Fig. 5e** and **Supplementary Table 3**); the remaining CTSSs were located at annotated exons (24-31%), followed by introns, distal intergenic regions and 3’ UTRs (each category,∼5-8%) (**Extended Data Fig. 5e** and **Supplementary Table 3**). With *NIPBL* downregulation, we observed consistent and disproportional enrichment (1.3-3.6 folds) of upregulated CTSSs at introns and distal intergenic regions, in all nine cancer cell lines (**Extended Data Fig. 5e** and **Supplementary Table 3**). These observations indicate that partial *NIPBL* loss can lead to global alternative promoter usage mainly from introns and distal intergenic regions in various cancer types.

### *NIPBL* perturbation leads to alternative promoter usage arising from LTR repetitive elements characterized by open chromatin features

To investigate the mechanisms of alternative promoter activation by *NIPBL* perturbation, we focused on the melanoma context. Previous studies have characterized the mammalian promoters by CAGE-seq as two classes, sharp and broad promoters^28,29^. Consistently, we observed similar bimodal distribution of sharp and broad promoters of all CTSSs at baseline with a 16 bp cut-off that separated sharp vs. broad promoters^28^ (**Fig. 2a**). Curiously, *NIPBL* downregulation preferentially affected CTSSs with broad promoter features (**Fig. 2a** and **Extended Data Fig. 6a**). Further, we noted the presence of repetitive elements, particularly LTRs in the *ALK*^ATI^*, ULK4, LINC01387* and *KEL* upregulated alternative promoters by *NIPBL* downregulation (**Fig. 1f**). Thus, we evaluated the genome-wide distribution of upregulated CTSSs at repetitive elements, particularly TEs, using multi-mapped CAGE-seq reads that were randomly assigned to one of their mappable genomic loci. At baseline, less than 3% of all CTSSs localized to annotated promoters contained repetitive elements, whereas approximately 45-48% of all CTSSs localized in the intronic or distal intergenic regions contained repetitive elements, particularly retrotransposons with approximately 10-17% of each of the LTR, LINE, and SINE subclasses of retrotransposons (**Fig. 2b**). With *NIPBL* downregulation, there was a marked increase in retrotransposon containing upregulated CTSSs, particularly LTRs, in intron (12.8% [shLuc] vs 38.8% [shNIPBL_#1] and 36.4% [shNIPBL_#2]), distal intergenic (16.7% [shLuc] vs 54.7% [shNIPBL_#1] and 59.1% [shNIPBL_#2]) and promoter (0.7% [shLuc] vs 19.6% [shNIPBL_#1] and 4.2% [shNIPBL_#2]) regions. Interestingly, there was a substantial decrease in SINE-containing upregulated CTSSs in intron (16.5% [baseline] vs 3.5% [shNIPBL_#1] and 3.8% [shNIPBL_#2]) and distal intergenic (10.4% [baseline] vs 2.1% [shNIPBL_#1] and 4.7% [shNIPBL_#2]) regions (**Fig. 2b**). Furthermore, we noted an enrichment of ERVL/ERVL-MaLR LTR families in upregulated LTRs in intron and distal intergenic regions after *NIPBL* knockdown (7.14%/58.3% [baseline] vs 14.1%/67.1% [shNIPBL_#1] and 8.84%/75.3% [shNIPBL_#2] for ERVL/ERVL-MaLR respectively) and a concomitant depletion of the ERV1 LTR family (25.1% [baseline] vs 10.6% [shNIPBL_#1] and 11.6% [shNIPBL_#2]) (**Extended Data Fig. 6b**). Accordingly, subtypes from ERVL-MaLR (e.g., THE1A and THE1D) and ERVL (e.g., LTR66) LTR families were selectively activated by shNIPBL (**Extended Data Fig. 6c**). These data indicated that LTR retrotransposons were preferentially activated by *NIPBL* downregulation.

**Figure 2.**
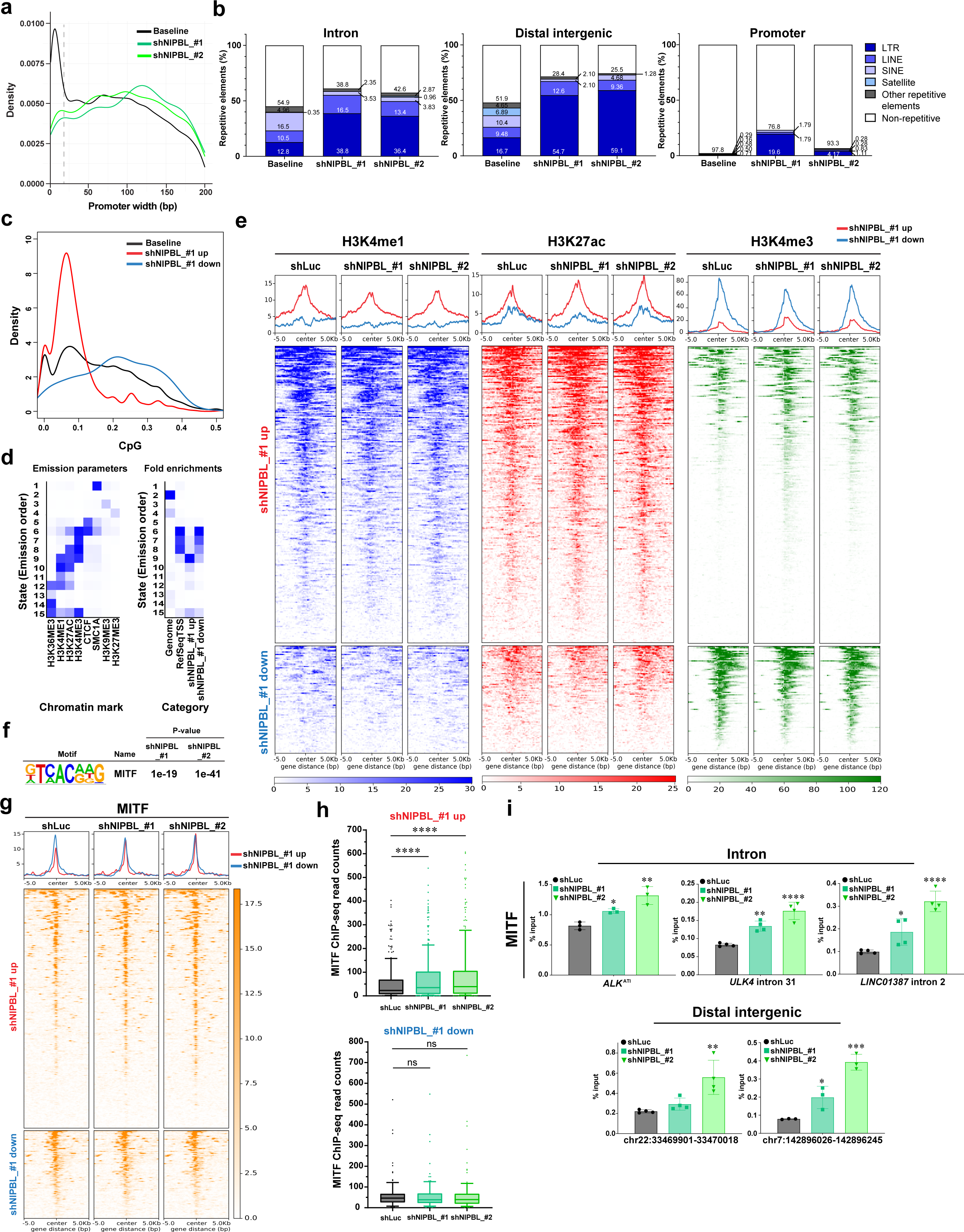
Loss of *NIPBL* induced alternative promoter usage arising from LTR repetitive elements characterized by open chromatin characteristics. **a,** Bimodal distribution of promoter width showing sharp and broad promoters at baseline (black); and the promoter width of differentially regulated CTSSs by shNIPBL perturbation in melanoma cells, demonstrating a preference for broad promoters (green). Grey dotted line depicted the cutoff at 16 bp that separated sharp and broad promoters based on the distance between CAGE tags for each clustered CTSS with TPM>0.5. **b,** Distribution of significantly upregulated CTSSs by shNIPBL at repetitive vs. non-repetitive elements in intron, distal intergenic and promoter regions. **c,** CpG compositions of significantly upregulated, downregulated and all CTSSs by shNIPBL_#1. **d,** Characterization of 15 chromatin states by ChromHMM (left) and enrichment of different chromatin states over genome distribution for RefSeq TSS and differentially regulated CTSS (right) under *NIPBL* perturbation conditions. **e,** Density plots of genome-wide distribution of indicated histone modification marks by CUT&RUN, centered on differentially upregulated and downregulated CTSSs by shNIPBL_#1. Data from one representative biological replicate (n=2) are shown. **f,** HOMER *de novo* motif analysis of significantly changed CTSSs at non-promoters demonstrated significant enrichment of the MITF motif by shNIPBL in melanoma cells. e, exponents of 10. **g,** Density plots of genome-wide distribution of MITF by ChIP-seq centered on differentially upregulated and downregulated CTSSs by shNIPBL, demonstrating an increase in MITF binding at upregulated CTSSs. Data from one representative biological replicate (n=2) are shown. **h,** Box and whiskers plots showing 10-90 percentile of MITF binding by ChIP-seq read counts at differentially upregulated (top panel) and downregulated (bottom panel) CTSSs by shNIPBL in melanoma cells. Each dot represented the quantification of MITF binding at one differentially regulated CTSS locus. ns, not significant; \*\*\*\**P*<0.0001, matched one-way ANOVA Dunnett’s multiple comparisons test. **i,** Quantification of MITF binding by ChIP-qPCR at representative alternative promoters in intron and distal intergenic regions under *NIPBL* perturbation conditions in melanoma cells. Data indicated the mean ± S.D. (n=3-4 biological replicates). \**P*<0.05, \*\**P*<0.01, \*\*\**P*<0.001, \*\*\*\**P*<0.0001, ordinary one-way ANOVA Dunnett’s multiple comparisons test.

Since LTR retrotransposons are generally silenced epigenetically by DNA methylation and repressive chromatin modifications in the genome^1,4,8,9^, we examined the local chromatin characteristics of the shNIPBL-upregulated CTSSs, including those containing LTRs. We first examined the CpG and C/G contents of the shNIPBL CTSSs. We observed substantially lower CpG and C/G representation in the shNIPBL_#1-upregulated compared to shNIPBL_#1-downregulated and baseline CTSSs (**Fig. 2c** and **Extended Data Fig. 6d**), and modestly lower CpG representation in the shNIPBL-#2-upregulated CTSSs than the -downregulated and baseline CTSSs (**Extended Data Fig. 6e**). To experimentally delineate the DNA methylation status at CTSSs with *NIPBL* perturbation, we performed bisulfite-free enzymatic methyl sequencing (EM-seq). We observed uniform sequencing coverage among replicates of shLuc, shNIPBL_#1 and shNIPBL_#2 (**Extended Data Fig. 7a**). *NIPBL* knockdown did not result in global changes in DNA methylation levels (**Extended Data Fig. 7b**). The baseline DNA methylation ratio at CTSSs exhibited a bimodal distribution, in line with prior studies^30,31^ without any obvious differences in shLuc, shNIPBL_#1 and shNIPBL_#2, consistent with the lack of global changes in DNA methylation after *NIPBL* downregulation (**Extended Data Fig. 7b-k**). We further evaluated DNA methylation levels at shNIPBL-differentially regulated CTSSs. We observed that the shNIPBL_#1-downregulated CTSSs and the shNIPBL_#2 up- and downregulated CTSSs exhibited enrichment of low DNA methylation ratios, while the shNIPBL_#1-upregulated CTSSs exhibited relatively uniform distribution across various DNA methylation levels (**Extended Data Fig. 7c-d**). However, the areas have relatively few CpG sites (**Extended Data Fig. 7e-f**), consistent with the low CpG content (**Extended Data Figure 6d, e**). We then specifically evaluated the representative CTSSs upregulated by shNIPBL_#1 and shNIPBL_#2, e.g., the *ALK* (intron 19), *ULK4* (intron 31), *LINC01387* (intron 2), *SYN3* (intergenic regions) (between *SYN3* and *LINC01640*) and *KEL* (intergenic regions) (between *TAS2R39* and *TAS2R40*) alternative promoter regions. These regions generally contained few CpG sites and had no or low levels of CpG methylation; the CpG methylation levels at these CTSSs were not altered by *NIPBL* perturbations (**Extended Data Fig. 7g-k**). Collectively, these results supported the notion that differentially activated CTSSs by *NIPBL* knockdown were not silenced by DNA methylation.

We next examined the chromatin characteristics of altered CTSSs by *NIPBL* loss, using cleavage under targets and release using nuclease (CUT&RUN)^32^ for histone modifications preferentially enriched at active enhancers (H3K4me1, H3K27ac), promoters (H3K27ac, H3K4me3), and silenced chromatin regions (H3K9me3, H3K27me3) (**Fig. 2d, e**). We further examined the genome-wide distribution of additional chromatin marks, e.g., H3K36me3, CTCF and SMC1A, to generate the 15 comprehensive chromatin states by ChromHMM analysis^33^ at baseline (**Fig. 2d**). As expected, the annotated TSS is mostly enriched for ChromHMM states 6 (H3K4me3/CTCF high)^34^, 7 (H3K4me3 high) and 8 (H3K27ac/H3K4me3 high), all are characterized by the enrichment of H3K4me3, most consistent with promoters (**Fig. 2d**). Accordingly, the significantly downregulated CTSSs by shNIPBL_#1, which were preferentially at promoters (**Fig. 1e**), were characterized by characteristic active promoter chromatin marks, e.g., H3K4me3 and H3K27ac, at baseline (**Fig. 2d, e**). Curiously, the significantly upregulated CTSSs by shNIPBL_#1 were most enriched for ChromHMM state 9 characterized by enrichment for active and poised chromatin marks (e.g., H3K27ac and H3K4me1) and devoid of repressive chromatin marks (e.g., H3K9me3 and H3K27me3) at baseline, reminiscent of active enhancers. Notably, a subset of these upregulated CTSSs were also enriched with H3K4me3 chromatin mark, at baseline, reminiscent of active or poised promoters (**Fig. 2d, e**). These active chromatin marks were further enriched at the significantly upregulated CTSS heatmaps with *NIPBL* downregulation and are concordant with CTSS transcriptional changes (**Fig. 2e** and **Extended Data Fig. 8a-c**). Moreover, we observed similar patterns of chromatin modifications (e.g., H3K4me3 and H3K27ac enrichment) for LTR-derived upregulated CTSSs with *NIPBL* downregulation (**Extended Data Fig. 8e-g**).

*De novo* and known motif analyses by Homer revealed that the significantly changed CTSSs by shNIPBL at non-promoter regions were most enriched for the motifs of MITF, a lineage-specific master regulator in melanocyte and melanoma^35^ (**Fig. 2f**, **Extended Data Fig. 9a, b** and **Supplementary Table 4).** To validate these findings, we performed ChIP-seq of MITF and found that the majority of differentially regulated CTSSs were bound by MITF in shLuc cells (**Fig. 2g, h** and **Extended Data Fig. 8d, h**). Consistent with the Homer motif analysis, differentially upregulated CTSSs enriched at intron and distal intergenic regions showed significantly increased binding of MITF with shNIPBL perturbation; and downregulated CTSSs enriched at promoter regions mostly showed no appreciable changes in MITF binding (**Fig. 2g, h** and **Extended Data Fig. 8d, h**). We confirmed the significant increase in MITF binding to selective shNIPBL-upregulated CTSSs localized in the intronic and intergenic regions by quantitative ChIP-qPCR of MITF (**Fig. 1f, 2g-i** and **Extended Data Fig. 8d, h, 10a**). We further observed enrichment of H3K4me3 at these selective upregulated CTSSs by CUT&RUN-seq and ChIP-qPCR, corroborating the corresponding transcriptional activation (**Fig. 1f** and **Extended Data Fig. 10b**). Moreover, the increase in MITF-binding to differentially upregulated CTSSs by *NIPBL* loss at transposable elements were preferentially at LTRs (**Extended Data 9b).** Collectively, these data demonstrated that loss of *NIPBL* led to alternative promoter usage arising from LTR repetitive elements that were already characterized by open chromatin features and localized in the intronic and intergenic regions of the genome. They also indicate that the transcriptional activation from these LTR retrotransposons is restricted despite open chromatin characteristics and may require additional chromatin structure changes that allow further recruitment of context-dependent master regulator transcription factors for activation, e.g., MITF, in melanoma cells.

### *NIPBL* partial loss leads to decrease in hierarchical TAD structures and activation of alternative promoters from LTRs

Previous studies of complete loss of the core cohesin family members on chromatin, either by biallelic genetic deletion of *Nipbl* in mouse hepatocytes^22^ or through auxin-inducible degron-mediated rapid protein degradation of RAD21^36^ have demonstrated global disappearance of all TADs. To evaluate the effects of haploinsufficiency loss of *NIPBL* on spatial organization of chromatin, we performed Hi-C in biological duplicates to generate contact maps at 10 kb resolution and analyzed the genome-wide chromatin interaction frequencies in melanoma cells with the doxycycline-inducible knockdown of *NIPBL* (shNIPBL*_*#2) and control (shLuc). The 3D chromatin organization is not uniform, but can be organized into hierarchies of TAD structures that correlated with more transcriptionally active chromatin regions when compared to single non-hierarchical structures^19,37,17,38^. We analyzed the Hi-C dataset with OnTAD, a TAD caller optimized for calling hierarchies of TAD with nested subTAD structures^19^. Hi-C analysis of the shLuc control cells revealed that most TADs contained a hierarchy of nested subTADs (**Fig. 3a**, green arrow) within a larger outer TAD structure (**Fig. 3a**, light blue arrow), and a minority of TADs existed as single structures (singleton TAD) and lacked hierarchy (**Fig. 3a**, dark blue arrow). Haploinsufficient loss of *NIPBL* resulted in significant weakening of TADs and preferential loss of the larger outer TAD structure with relative preservation of the inner nested subTADs (**Fig. 3a**). Strikingly, the dissolution of outer TAD structures was manifested by the loss/weakening of corner dots, representing the disappearance of the persistent cohesin-mediated interaction between the most distal boundaries of the hierarchical TADs^39^. We defined the hierarchy of TADs based on the number and layers of nested subTADs, where higher levels of hierarchical TADs correlated with more successive layers of nested subTADs. We then quantified the number of singleton and hierarchical TADs (**Fig. 3b**). A total of 3791 TADs were identified in shLuc control cells, of which 967 (25.5%) were singleton TADs and 2824 (74.5%) were hierarchical TADs. Compared to the shLuc control cells, with *NIPBL* loss (shNIPBL_#2), there was a substantial increase in the total number of TADs to 4541; particularly, there was an increase in the proportion of singleton TADs (1723/4541, 37.9%) with a simultaneous decrease in hierarchical TADs (2818/4541, 62.1%) (**Fig. 3b**). The decrease in the percentage of hierarchical TADs was more pronounced in level 3 and above hierarchical TADs, 38.2% (1448/3791, shLuc) vs. 27.3% (1239/4541, shNIPBL_#2) (**Fig. 3b**), which is likely due to the dissolution of higher-level hierarchical TADs ≥ level 3 into lower level hierarchical or singleton TADs (**Fig. 3a, b**). Consistently, with partial *NIPBL* loss, we observed an overall shift of TAD size to significantly smaller TADs, with a median TAD size of 530 kb in shLuc control and 340 kb in shNIPBL_#2 cells (**Fig. 3c**). Similarly, Hi-C read-pair revealed shNIPBL_#2 cells contained more short-range interactions of ∼10-100 kb (log10 4-5) while shLuc control cells contained more mid-to-long-range interactions ∼10 kb-3 Mbp (log10 5-6.5) (**Extended Data Fig. 11a**). Furthermore, in agreement with previous study^22^, partial loss of *NIPBL* resulted in finer segregation of compartments (**Extended Data Fig. 11b**). Interestingly, we observed more mega-loop long-range interactions (>3 Mbp, log10 >6.5) in melanoma cells with shNIPBL_#2 perturbation than shLuc control (**Extended Data Fig. 11a**). We also evaluated the TAD boundary strength by insulation score^40^ and observed a significant reduction in shNIPBL_#2 compared to shLuc control cells (**Fig. 3d**). This data is consistent with prior studies of cohesin perturbations that led to TAD boundary defects^22,36,41^, and indicate that partial *NIPBL* loss can also lead to weakened TAD boundaries.

**Figure 3.**
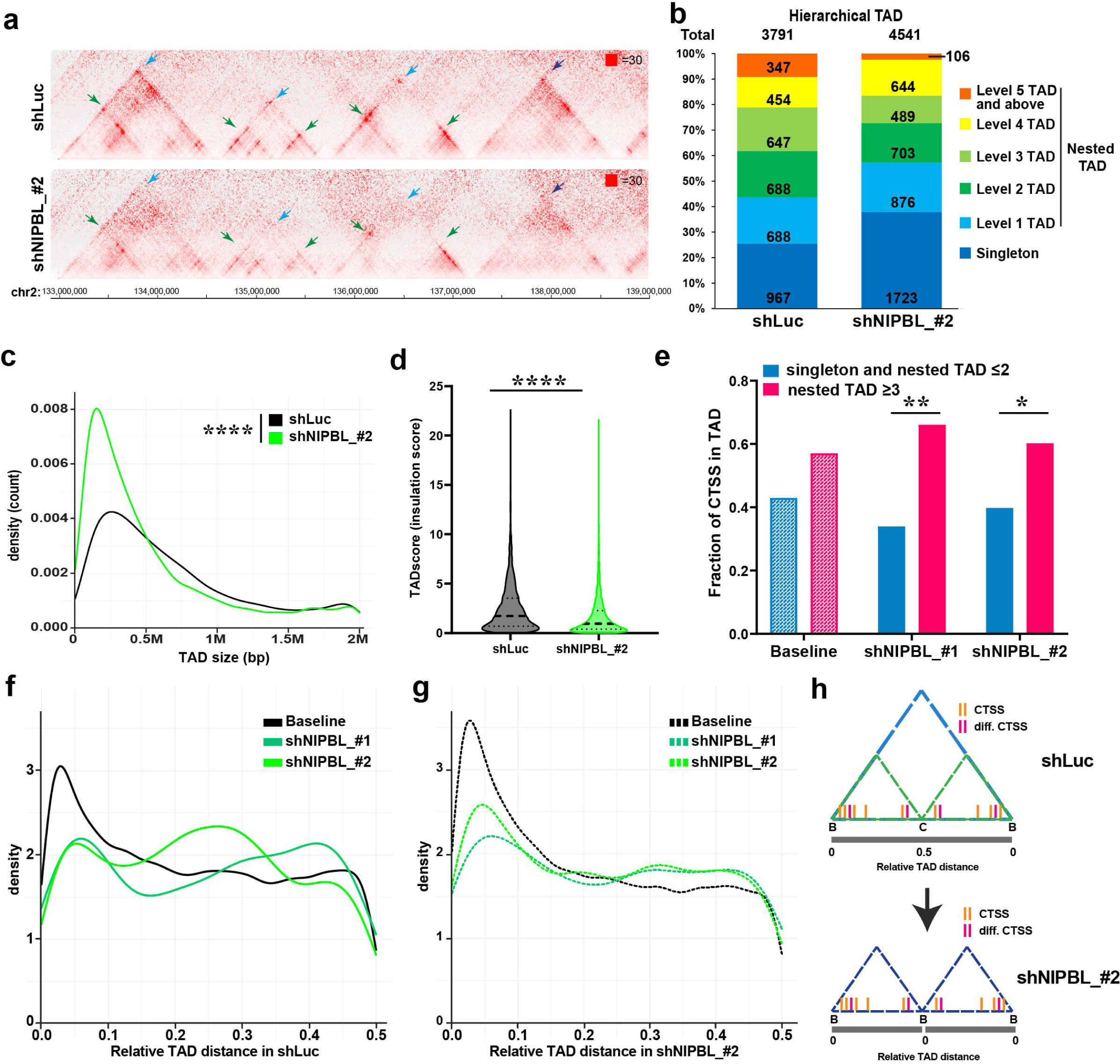
*NIPBL* partial loss leads to a decrease in hierarchical TAD structures and preferentially affected CTSSs resided in high-level hierarchical TADs. **a**, Representative hierarchical TAD structures by Hi-C contact maps (10 kb resolution) under control and *NIPBL* perturbation conditions in melanoma cells. TADs are organized as singletons (navy blue arrows) or hierarchical structures characterized by nested subTADs (green arrows) and nested inside meta-TADs (light blue arrows). *NIPBL* downregulation weakened TAD structures and preferential loss of outermost TADs (light blue arrows) of hierarchical TADs. n=2 biological replicates. **b-d,** The effects on hierarchical TAD structures by partial loss of *NIPBL* in melanoma cells, including on the number of various levels of hierarchical TAD structures by OnTAD (**b**), size distribution (**c**), and TAD boundary/insulation scores (**d**). \*\*\*\**P*<0.0001, Mann-Whitney two-tailed unpaired non-parametric t-test. **e,** Comparison of fraction of baseline CTSS versus differentially regulated CTSSs by shNIPBL_#1 and shNIPBL_#2 in hierarchical and non-hierarchical TADs defined under control condition (≥ level 3 vs. ≤ level 2 and singleton). \**P*<0.05, \*\**P*<0.01, two-tailed chi square test. Number of CTSS in each category is in Extended Data Figure 12. **f-g,** Relative CTSS distance to nearest TAD boundaries defined in shLuc conditions (**f**) and in shNIPBL conditions (**g**). The relative CTSS distance to TAD boundaries was calculated as the distance of CTSS to the nearest TAD boundary divided by the size of the respective TAD defined in the shLuc and shNIPBL conditions. **h,** Schematics illustrating the change in location of CTSSs relative to TAD structure changes with *NIPBL* perturbation.

We next evaluated the effect of changes in hierarchical TAD structures following *NIPBL* knockdown on differential CTSSs expression. We observed imbalanced distribution of differential CTSSs localized to hierarchical TADs. There was a significant enrichment of differential CTSSs localized to ≥ level 3 hierarchical TADs compared to lower levels of hierarchical and singleton TADs that were defined in the control condition, with both 34% (sh*NIPBL*_#1) and 56% (sh*NIPBL*_#2) downregulation of *NIPBL* compared to shLuc controls in melanoma cells (**Fig. 3e** and **Extended Data Fig. 12**).

It has been reported that actively transcribed genes frequently clustered at TAD boundaries that contained nested subTADs^38,42,43^. We delineated the relative positions of CTSSs in individually normalized TADs defined under control and *NIPBL* perturbation conditions. The relative CTSS position was calculated as the relative CTSS distance within TAD to the nearest TAD boundary normalized by the respective TAD size, with the TAD boundary defined as 0 and TAD center as 0.5. At baseline (shLuc), the majority of the CTSSs were localized closer to TAD boundary than TAD center (**Fig. 3f, h**). In contrast, the differentially regulated CTSSs by shNIPBL_#1 and shNIPBL_#2 were distributed closer to the TAD center when the TADs were defined in the shLuc control condition (**Fig. 3f, h**). Interestingly, with the shNIPBL_#2-associated decrease in TAD hierarchy and TAD structure changes, the differentially regulated CTSSs by shNIPBL_#1 and shNIPBL_#2 were now localized at the boundary of the redefined TADs in the shNIPBL_#2 condition (**Fig. 3g, h**). These data indicate that the TAD hierarchical structure changes can directly impact transcriptional activation from alternative promoters.

### Promiscuous gene activation through reorganization of hierarchical TADs and retargeting of enhancers to alternative promoters in proximity

To investigate the molecular mechanisms by which shNIPBL-mediated TAD hierarchy perturbation affect alternative promoter activation, we selectively investigated the 3D chromatin topology around activated alternative promoters, e.g., *ALK* intron 19 (**Fig. 4a, b**) and *ULK4* intron 31 (**Fig. 4c, d**). Both alternative promoters (highlighted in blue) were situated inside complex hierarchical TADs (≥level 5) that contained multiple layers of nested subTADs in control shLuc cells (**Fig. 4a-d**). With partial reduction of *NIPBL*, the Hi-C contact matrix revealed loss of the outer TAD structures (dark blue circles) with relative preservation of the inner nested subTADs (green circles) and splitting of hierarchical TADs into multiple smaller TADs (grey horizontal bars) with lower levels of hierarchy (**Fig. 4a, c**). Consistently, OnTAD hierarchical domain analysis of the *ALK* intron 19 (**Fig. 4b**) and *ULK4* intron 31 (**Fig. 4d**) alternative promoter regions revealed decreased hierarchical TAD structures, from ≥ level 5 hierarchical TADs to ≤ level 4 hierarchical TADs. Additionally, both alternative promoter regions that were initially embedded in the center of a complex network of hierarchically nested subTAD structures in the shLuc control condition are now repositioned to the boundary of split and “simplified” TAD structures with reduced hierarchy following *NIPBL* loss (**Fig. 4b, d**).

**Figure 4.**
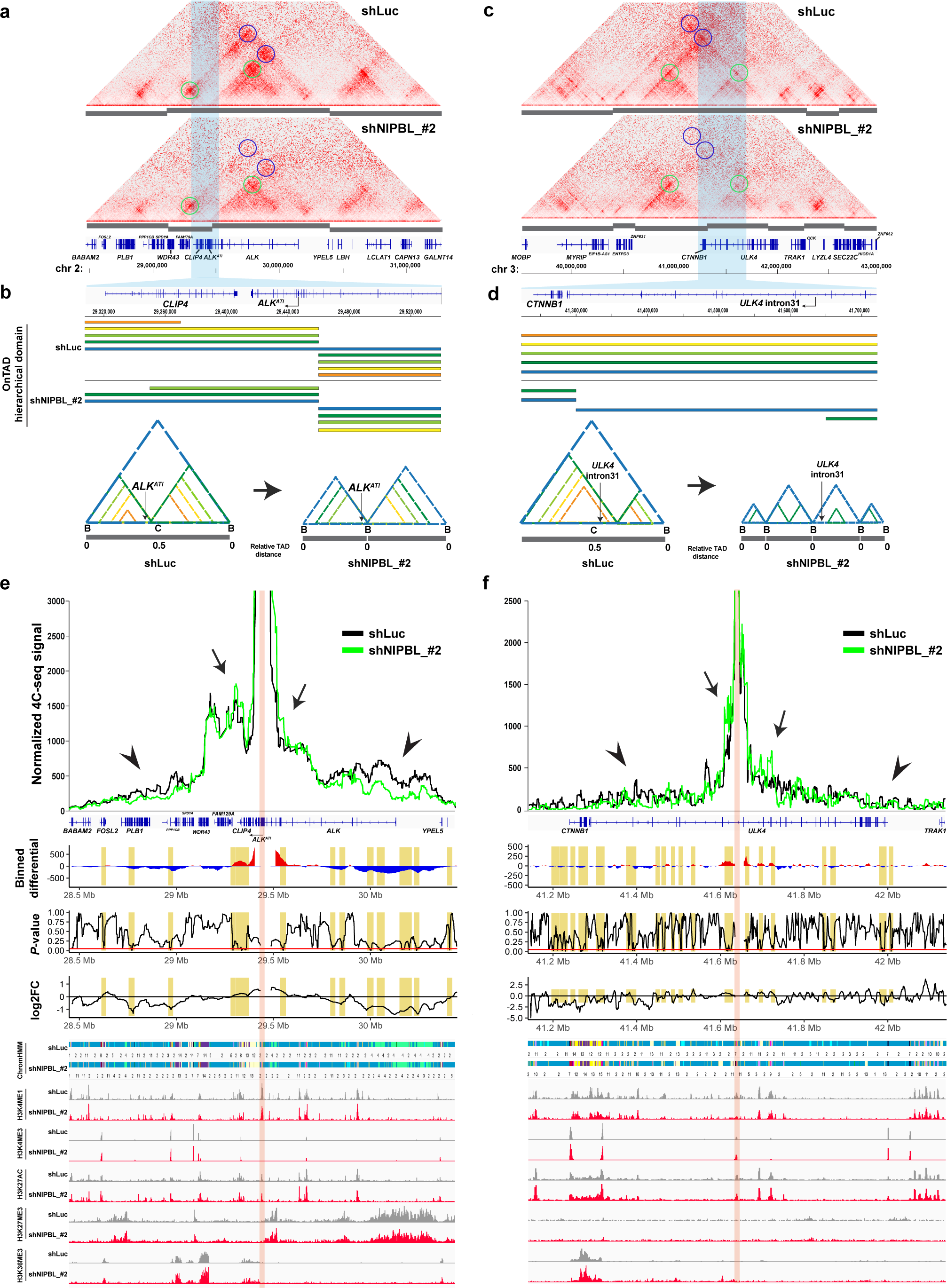
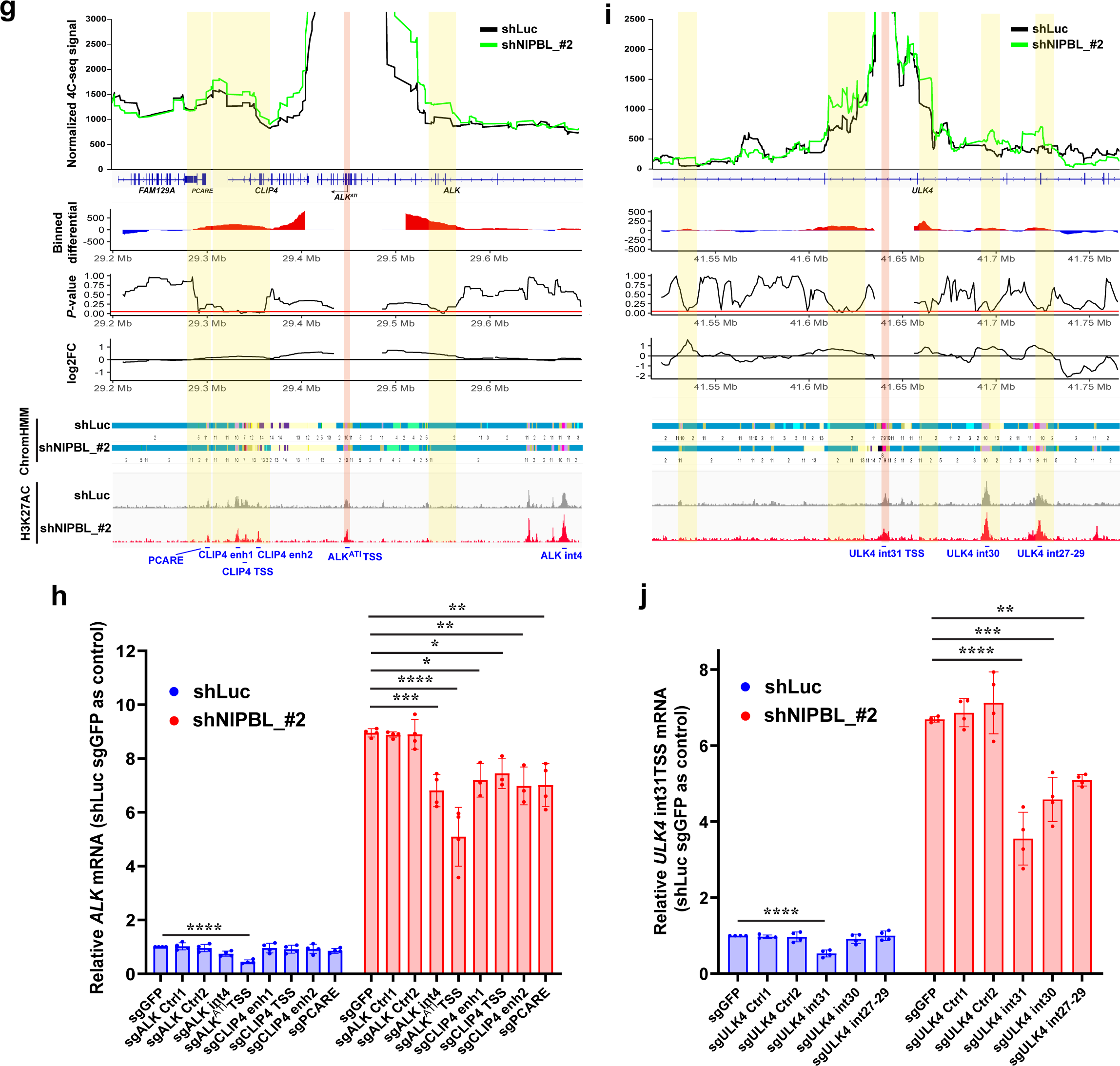
Examples of promiscuous gene activation (*ALK*^ATI^, *ULK4* from intron 31) through reorganization of hierarchical TADs and retargeting of enhancers to alternative promoters in proximity. **a, c**, Hi-C map of hierarchical TADs of selective genomic regions, *ALK*^ATI^ (**a**) and *ULK4* (**c**) in control (top, shLuc) and *NIPBL* perturbation (bottom, shNIPBL_#2) conditions in melanoma cells. Grey bars below each Hi-C map denotes computationally annotated TADs. *ALK*^ATI^ and *ULK4* are located in a nested TAD comprising multiple subTADs. *NIPBL* loss leads to splitting (grey bars) and loss of hierarchy of outer TADs (blue circles), with relative preservation of subTADs (green circles). Genomic regions of *ALK*^ATI^ and *ULK4* intron 31 alternative promoters were shaded in blue. **b, d,** OnTAD hierarchical domains changes of selective genomic regions, *ALK*^ATI^ (**b**) and *ULK4* (**d**) in control (shLuc) and *NIPBL* perturbation (shNIPBL_#2) conditions in melanoma cells. The hierarchical domains are color-coded similarly as Figure 3b, demonstrating the split of the top outer level of the hierarchical TADs (blue bar). Bottom illustration: B, TAD boundary. C, TAD center. **e-f**, Representative normalized 4C-seq profiles under control and *NIPBL* perturbation conditions, using *ALK*^ATI^ (**e**) and intron 31 of *ULK4* (**f**) as viewpoints (shaded in pink), demonstrating enhanced interaction profile close to the viewpoint (arrows) and diminished long-range interaction (arrow heads) with *NIPBL* downregulation. Normalized average 4C-seq signal was binned with either a 25 kb window (*ALK*^ATI^ locus) or 10 kb window (*ULK4* intron 31 locus) with a 1 kb shift to calculate differential and log2FoldChange in interaction (shNIPBL_#2 signal – shLuc signal) and statistical significance by two-tailed paired t-test. Respective chromatin states by ChromHMM defined in Fig. 2d at the corresponding genomic regions were also shown. **g-j**, Zoomed-in regions from normalized 4C-seq profile under control and *NIPBL* perturbation conditions, demonstrating proximal contacts gain surrounding *ALK*^ATI^ (**g**) and *ULK4* intron 31 (**i**) alternative promoters with *NIPBL* perturbation. CRISPRi of the indicated genomic regions (blue, **g, i**) confirmed engagement of neighboring proximal H3K27ac-enriched peaks as retargeted enhancers for the transcriptional activation of *ALK*^ATI^ (**h**) and *ULK4* from intron 31 (**j**). Data indicated the mean ± S.D. (n=3-4 biological replicates). \**P*<0.05, \*\**P*<0.01, \*\*\**P*<0.001, \*\*\*\**P*<0.0001, ordinary one-way ANOVA Dunnett’s multiple comparisons test.

TADs are thought to facilitate and restrict long-range enhancer-promoter (E-P) interactions within and outside of TADs, respectively^17^. We speculate that the novel enhancer-alternative promoter (E-altP) interactions could be established because of hierarchical TAD structure changes. To investigate if changes in local chromatin architecture directly affect transcription from *ALK* intron 19 and *ULK4* intron 31, we selectively disrupted CTCF insulators located around the alternative promoter regions by either dCas9-KRAB-mediated blocking of CTCF binding or Cas9-mediated deletion of the CTCF binding sites at the TAD/sub-TAD boundaries (**Extended Data Fig 13a, 14a**). dCas9-KRAB that targeted cognate CTCF binding motifs resulted in significant reduction of CTCF binding by ChIP-qPCR (**Extended Data Fig 13b-f, 14b-d**). Specifically, we observed increased expression of alternative transcripts from *ALK* intron 19 and *ULK4* intron 31 when CTCF insulators at TAD/sub-TAD boundaries were disrupted by dCas9-KRAB (**Extended Data Fig 13g, 14e**). Consistently, we observed that that Cas9-mediated deletion of CTCF binding sites at TAD/sub-TAD boundaries similarly enhanced alternative transcripts expression at both loci (**Extended Data Fig 13h-i, 14f-g**), indicating that hierarchical TAD structure changes can impact alternative promoter usage, likely through rearrangement of E-altP interactions.

To more directly assess the genome-wide E-altP interactions that may contribute to the transcriptional activation of the *de novo* CTSSs, we performed 4C-seq^45^, using the *ALK* and *ULK4* alternative promoters in intron 19 and intron 31 as viewpoints, respectively (**Fig. 4e, f**, red shades). In the controls (shLuc), we observed a high frequency of interactions from genomic regions close to the viewpoints and a rapid decay of interaction frequencies with increased genomic distance from the alternative promoters by 4C-seq. With partial *NIPBL* loss (shNIPBL_#2), we observed a substantial decrease of interaction frequencies with the alternative promoters, preferentially from distal genomic regions (**Fig. 4e, f**, arrow heads); in contrast, there was a marked increase in interactions from the proximal genomic regions (**Fig. 4e, f**, arrows, and **Fig. 4g, i**). These data indicate that with the collapse of high-level hierarchical TAD structures, there was a shift of genomic interactions and possibly retargeting of enhancers from distal to proximal genomic regions to the alternative promoters.

Consistent with the genome-wide chromatin analysis (**Fig. 2d**), the alternative promoters of *ALK* and *ULK4* were devoid of heterochromatin marks, e.g., H3K27me3, and were enriched for open chromatin features, e.g., H3K4me1, in shLuc control cells (**Fig. 4e, f**). With *NIPBL* loss (shNIPBL_#2), there was no significant change in H3K4me1 enrichment at the alternative promoters (**Fig. 4e, f**). Consistent with transcriptional activation, we observed increased enrichment of H3K4me3 and H3K27ac at the alternative promoters, and of H3K36me3 at the gene bodies of the alternative transcripts of *ALK* and *ULK4* (**Fig. 4e, f**). Moreover, we observed multiple H3K27ac enriched peaks throughout the genomic regions flanking *ALK* intron 19 and *ULK4* intron 31 within the hierarchical TADs, suggesting that they may function as enhancers to the alternative promoters. To validate this, we performed CRISPRi to target the various potential enhancers, using guide RNAs specific for the *ALK* (intron 19) or *ULK4* (intron 31) alternative promoters as positive and non-targeting guide RNA (sgGFP) as negative controls (**Fig. 4g-j**). We also included two sets of sgRNAs (sgALK Ctrl1 & 2, sgULK4 Ctrl1& 2) targeting neighboring non-enhancer regions (without H3K27ac enrichment) of the *ALK* (intron 19) and *ULK4* (intron 31) alternative promoters as additional controls (**Fig. 4h, j**). At baseline (shLuc), there were marginally detectable levels of *ALK* and *ULK4* alternative transcripts. As expected, CRISPRi-sgRNAs targeting the *ALK* intron 19 (sgALK^ATI^ TSS) and *ULK4* intron 31 (sgULK4 int31 TSS) alternative promoters effectively inhibited the mRNA expression of *ALK* and *ULK4* by ∼58% and ∼54%, respectively (**Fig. 4g-j**). Interestingly, in shLuc control cells, although CRISPRi-sgRNAs targeting selective altP-distal enhancer region, e.g., TSS of *PCARE* (sgPCARE), modestly but significantly decreased *ALK* expression; CRISPRi-sgRNAs targeting more altP-proximal enhancer regions, e.g., *CLIP4* TSS (sgCLIP4 TSS), *CLIP4* enhancer 1 (sgCLIP4 enh1) and *CLIP4* enhancer 2 (sgCLIP4 enh2), did not significantly alter the expression of *ALK* (**Fig. 4g, h**). With *NIPBL* loss (shNIPBL_#2), there was a significant increase of *ALK*^ATI^ expression (∼9.5 fold); CRISPRi-sgRNAs targeting both the distal and proximal genomic regions (e.g., sgPCARE, sgALK int4, sgCLIP4 TSS and sgCLIP4 enh1/2) to *ALK* intron 19 all significantly reduced the expression of *ALK*^ATI^, whereas targeting the altP-neighboring non-enhancer regions (sgALK Ctrl1 & 2) did not, confirming the recruitment of the proximal enhancers to further activate the transcription of *ALK*^ATI^ from intron 19 of *ALK*^ATI^ (**Fig. 4h**). Similarly, for the *ULK4* intron 31 locus, we observed further recruitment of proximal enhancers (e.g., *ULK4* int27-29) with *NIPBL* downregulation (**Fig. 4i, j**).

These observations corroborated the 4C-seq data. They indicated that in the controls with relatively preserved hierarchical TADs, the distal enhancers provided basal interactions with alternative promoters, but the basal interactions did not provide sufficient strength to activate transcription from alternative promoters. With *NIPBL* loss and the collapse of the hierarchical TADs, there was a shift of E-altP interactions to recruit proximal enhancers to activate transcription from alternative promoters arising from LTR repetitive elements. Notably, there were no significant changes in the H3K27ac enrichment or local ChromHMM states at either the distal or the proximal genomic regions relative to the alternative promoters, suggesting that the candidate enhancers particularly the proximal E-altP interactions were restricted by hierarchical TAD structures and this restriction was released when the hierarchical TAD structures were altered as a result of *NIPBL* downregulation.

### *NIPBL* loss-mediated alternative promoter usage from LTR can give rise to oncogene expression

To further evaluate the functional consequences of *NIPBL* loss-mediated alternative promoter usage, we noted the robust induction of the *ALK*^ATI^ variant originating from the LTR16B2 region in intron 19 in melanoma cells (F**ig. 1a, b, f**). This *ALK*^ATI^ variant has been previously characterized as a novel oncogenic *ALK* isoform that is biallelically expressed and independent of genetic alterations at the *ALK* locus in ∼11% of melanomas and sporadically in other cancer types^26^. To evaluate if *NIPBL* perturbations contributes to the expression of *ALK*^ATI^ in melanoma, we selected the top 50 cases of TCGA-SKCM samples that had high level of *ALK*^ATI^ expression by RNA-seq, compared to bottom 50 cases that had no or very low expression of *ALK*^ATI^, and analyzed the *NIPBL* somatic mutations by FunSeq2^46^, a validated computational algorithm that can prioritize mutational variants with functional significance using a weighted score system (**Fig. 5a** and **Extended Data Fig. 15**). Combining the number of *NIPBL* somatic mutations in each tumor sample and the FunSeq2 scores, we generated a composite value that estimated the functional inactivation of *NIPBL* mutations for each case (**Fig. 5a** and **Extended Data Fig. 15**). The composite FunSeq2 score was significantly higher (*P<0.05*), indicating higher level of *NIPBL* functional inactivation, in melanoma tumor samples that expressed high levels than those that had no-low expression levels of *ALK*^ATI^ (**Fig. 5a**). To further establish the link between *NIPBL* perturbation and *ALK*^ATI^ expression, we screened available patient derived melanoma cell lines for *ALK*^ATI^ expression and identified SKMEL-23 and SKMEL-1128 with *ALK*^ATI^ expression by RNA-seq (**Fig. 5b**). Comparing to two well characterized melanoma cell lines, COLO800 and A375, that did not express *ALK*^ATI^, SKMEL23 and SKMEL-1128 express significantly lower levels of *NIPBL* mRNA transcripts (**Fig. 5c**). Consistently, H3K27ac ChIP-seq revealed H3K27ac enrichment at the *ALK*^ATI^ alternative promoter in all the melanoma cell lines examined^26^ (**Fig. 5d**), suggesting that melanoma might be primed to express *ALK*^ATI^. Surrounding the alternative promoter of *ALK*^ATI^ in intron 19, there were multiple H3K27ac enriched peaks at the proximal genomic regions that can potentially function as enhancers to interact with the alternative promoter of *ALK*^ATI^. We performed H3K27ac HiChIP to investigate the enhancer connectome and observed significant interactions between the proximal H3K27ac enriched enhancers with *ALK* intron 19. Moreover, we observed that genomic regions, e.g., the 5’uprestream, introns 1 and 4 of the *ALK* locus, and surrounding the *PCARE* locus, were the main E-altP interaction sites with the alternative promoter of *ALK* by virtual 4C and H3K27ac HiChIP hotspot analysis using *ALK* intron 19 as viewpoint (**Fig. 5d**, blue highlights). We performed CRISPRi to specifically target H3K27ac enriched enhancer regions identified by H3K27ac HiChIP that have the strongest interactions with the *ALK* intron 19 alternative promoter in *ALK*^ATI^-expressing melanoma cells. As anticipated, CRISPRi-sgRNA targeting the intron 19 alternative promoter of *ALK*^ATI^ most effectively reduced the transcription of *ALK*^ATI^ by ∼87% and 68% in SKMEL-23 and SKMEL-1128 cells, respectively (**Fig. 5e**). We observed more robust reduction of *ALK*^ATI^ expression by CRISPRi-sgRNA targeting the more proximal enhancers (e.g., sgALK int4, sgPCARE) compared to more distal enhancers (e.g., sgALK5’Upstream). Together, these observations suggest that *NIPBL* somatic mutations and reduced expression can induce context-dependent oncogene activation through proximal enhancer retargeting and alternative promoter usage.

**Figure 5.**
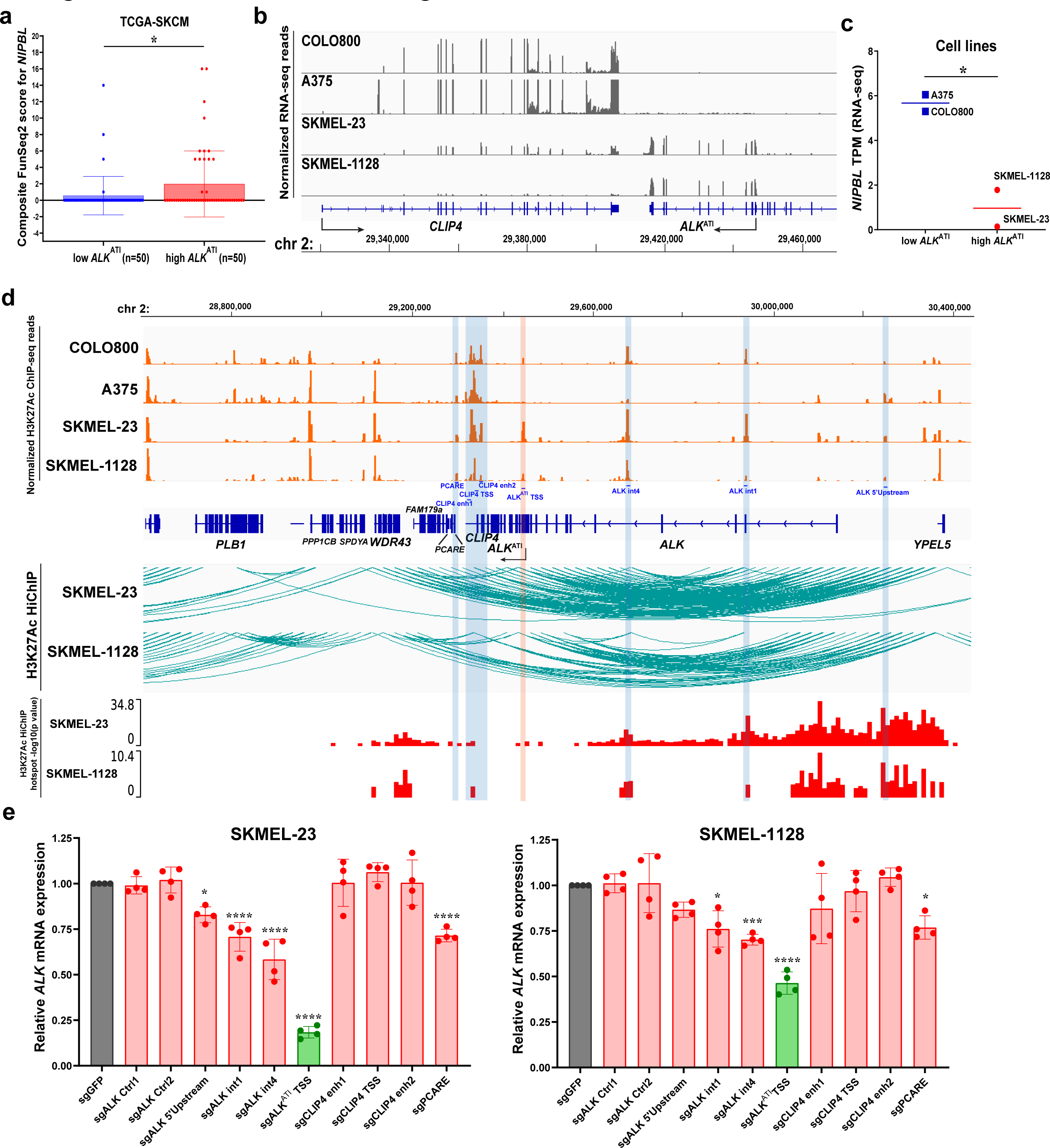
*NIPBL* perturbation contributes to transcriptional activation of *ALK*^ATI^ from the alternative promoter through engagement of proximal H3K27ac-enriched enhancers in melanoma. **a**, Evaluation of *NIPBL* functional perturbation by FunSeq2 of TCGA melanoma cases with high and low *ALK*^ATI^ expression. 50 cases from each of the high and low *ALK*^ATI^-expressing TCGA-SKCM cases were used to calculate the composite FunSeq2 score of *NIPBL*. \**P*<0.05, two-tailed unpaired t-test. **b,** Representative RNA-seq profiles of the *ALK* locus, demonstrating the expression of *ALK*^ATI^ in SKMEL-23 and SKMEL-1128, but not in COLO800 and A375 melanoma cell lines. **c,** Comparison of *NIPBL* transcript per million (TPM) in *ALK*^ATI^-expressing and non-expressing melanoma cell lines. n=2 biological replicates \**P*<0.05, two-tailed unpaired t-test. **d,** Normalized ChIP-seq profiles of H3K27ac (Top) demonstrating putative proximal enhancers (blue shade) surrounding the *ALK*^ATI^ alternative promoter (pink shade); H3K27ac HiChIP arcs with hotspot analysis (bottom) using the *ALK*^ATI^ alternative promoter in intron 19 (pink shade) as virtual viewpoint confirmed the proximal enhancer interactions (blue shades) with the *ALK*^ATI^ alternative promoter. **e,** CRISPRi of the indicated proximal H3K27ac-enriched putative enhancers regions indicated in **d** led to a reduction of *ALK*^ATI^ mRNA expression, confirming their contribution as enhancers. Data indicated the mean ± S.D. (n=4 biological replicates). \**P*<0.05, \*\**P*<0.01, \*\*\**P*<0.001, \*\*\*\**P*<0.0001, ordinary one-way ANOVA Dunnett’s multiple comparisons test.

Finally, we investigated whether MITF plays a role in transcriptional activation of *ALK*^ATI^ in melanoma cell lines, analogous to its recruitment to activate alternative promoter after *NIPBL* loss (**Fig. 2f-h** and **Extended Data Fig. 8d, h, 9a, b**). Compared to known MITF binding sites at *TYR* and *DCT* promoters, we observed comparable levels of MITF binding to *ALK*^ATI^ proximal H3K27ac-enriched enhancers by MITF ChIP-qPCR in both SKMEL-23 and SKMEL-1128 (**Extended Data Fig. 16a**). Downregulation of MITF expression by RNAi (siMITF_#1 and siMITF_#2) led to significant decrease of *ALK*^ATI^ expression (**Extended Data Fig. 16b, c**), indicating that MITF binding is critical in the active transcription of *ALK*^ATI^ from the *ALK* intron 19 alternative promoter.

## Discussion

Promoter usage is highly dynamic in cancer and the resulting isoform diversity could potentially increase the fitness of cancer cells^47^. Compared to normal tissue, cancer-specific alternative promoters are prevalent in various cancer types^47^. Further, cancer types such as ovarian and stomach cancers that pervasively use cancer-associated alternative promoters are also those that express more TE-encoded oncogenes^14,47^, suggesting that TE co-option might be one of the underlying mechanisms that drive alternative promoter usage in cancer. Among TEs, LTRs were found to be most enriched as onco-exapted TEs due to the presence of TFBS and splice donors^7,14,15^. Although TEs contribute significantly to the repertoire of TFBS in genome during transposition^1,48^, tight control of TE expression is required to maintain genome stability. Multiple layers of interconnected epigenetic mechanisms have been described, including DNA methylations and repressive chromatin modifications, to restrict TE expression^29^. Here, we report a novel mechanism of TE regulation by the 3D chromatin hierarchy to specifically restrict the activation of TEs that already possess active chromatin features, e.g., H3K4me1, H3K27ac, and the lack of CpG methylation, as active promoters.

Highly diverse ERVs/LTRs emerged as the key contributors in disseminating lineage-specific TFBS in the human genome, where selected subfamilies of LTRs are enriched for binding sites of specific transcription factors^49,50^. Consistently, our study observed that in melanoma cells, the melanocyte/melanoma-specific master regulator MITF^35^ was responsible for activating alternative promoters from LTRs that were already characterized by enhancer/promoter chromatin features, following the collapse of hierarchical TAD structures. We found that *NIPBL* downregulation can also lead to alternative promoter activation in multiple cancer types in addition to melanoma (**Extended Data Fig. 5** and **Supplement Table 3**), suggesting that 3D chromatin hierarchy likely also restricts alternative promoter activation in cancer types other than melanoma. It is possible that other lineage/cell context-specific transcription factors may regulate cell type and lineage-dependent alternative promoter usage from TEs under regulation of higher-order chromatin structures. This will be an area of interest that warrants future investigation. A previous study has hinted at the transactivation potential of TEs that are bound by transcription factors with the decoration of active histone modifications marks (e.g., H3K27ac, H3K4me1 and H3K4me3) and lower DNA methylation compared to their counterparts that are not associated with transcription factors^33^. We found in our study that most of the TE-derived alternative promoters have low levels of MITF binding at baseline, which is further enriched by the loss of *NIPBL*. The increase in MITF at transposable elements is preferentially at LTR (**Fig. 2f-i**, **Extended Data Fig. 8 and 9**). Consistently, these regions are also decorated with a high H3K27ac level and H3K4me1/H3K4me3 ratio, closely resembling conventional enhancer elements. Recent studies indicate that the transcriptional machinery and architecture are strikingly similar between promoter and enhancer cis-regulatory elements^51,52^.

Whether a cis-regulatory element results in productive mRNA depends on its transcriptional strength^52^. Consistently, our findings support that this class of enhancer-like TEs is typically restricted from transcriptional activation by hierarchical 3D chromatin organization. Upon the collapse of hierarchical TAD structures, long-range interactions between distal enhancers and promoters enabled by cohesin are lost; instead, H3K27ac-enriched regions close to TEs are recruited to enhance the transcriptional strength, resulting in productive mRNA transcription. Notably, the upregulated CTSSs have lower CG content or CpG methylation levels than CTSSs that are either downregulated (mostly canonical promoters) or unchanged. These observations have suggested that these types of CTSSs are unlikely to be repressed by DNA methylation; they also highlight one of the differentiating features between enhancers and promoters in that enhancers are usually devoid of DNA methylation and have low CpG content.

Enhancer is one of the key determinants for context-specific gene expression^43,44^. Transcription is an inherently stochastic process that is episodic and pulsative in nature^45,46^. The expression level of a gene depends on the transcriptional burst frequency which in turn is controlled by enhancer activity and contact frequency^47–49^. Transcriptome-wide studies revealed a strong linear relationship between H3K27ac density in enhancer and burst frequencies^47^. On the other hand, transcription depends on the contact probability of enhancers with promoters as enhanced or restricted by TADs, although not in a linear manner^50^. Consistent with short-term degradation of cohesin, as well as CdLS mouse models and patient studies^22,36,53^, we found defects in *NIPBL* function caused minor perturbations in gene expression. Supporting these observations, high-resolution Micro-C studies revealed that fine-scale enhancer-promoter (E-P) interactions predominantly lacked CTCF and cohesin occupancy^38^, and were largely robust to acute loss of cohesin^54^. Nevertheless, disruption of CTCF/cohesin-associated TAD structures leads to rewiring of long-range E-P interactions and cause aberrant gene expression^55^. In addition, a recent study found that cohesin is exclusively required for the activation of distant genes by long-range enhancers^56^. Collectively, we speculate that a subset of genes that are under control by long-range enhancers is preferentially affected by loss of NIPBL/cohesin. Based on our study, this set of genes is enriched for CTCF (**Fig. 2d**).

It remains debatable whether alternative promoters arise from imprecise control of transcription, or if they are more purposefully generated and provide adaptive functions at the organism level by expanding the diversity of the transcriptome^57^. It is possible that, in some cases, alternative transcripts are functionally distinct and allow fine-tuning of genetic functions in a versatile manner by virtue of differential stability, translational efficiency, protein structure and binding partners^58^. Furthermore, various cancer types are found to extensively use cancer-specific alternative promoters that are otherwise not used in normal tissues. Differential isoforms expression can be used to stratify cancer molecular subtypes and predict patient survival, highlighting the distinctive functional roles of certain transcript isoforms^47,59^. More functional studies on individual gene isoforms will ascertain the benefits of such genetic diversity in both benign and malignant conditions. On the other hand, the generation of chimeric or novel transcripts from transcriptional activation at LTRs in cancer might generate neoantigens that can be exploited by immuno-oncology therapeutics^60,61^.

## Materials and Methods

### Cell culture

501mel cell line was a gift from Levi A. Garraway laboratory (Broad Institute/Dana-Farber Cancer Institute) and was cultured in Ham’s F-12 medium. COLO800 and A375 cell lines were gifts from Joan Massague laboratory (MSKCC) and were cultured in RPMI and high glucose DMEM media respectively. SKMEL-23 was a patient-derived melanoma cell line established and provided by Taha Merghoub laboratory (MSKCC) and cultured in RPMI medium. SKMEL-1128 was developed from a melanoma tumor biopsy core (MSKCC) and grafted subcutaneously in SCID mice to establish tumor. Tumor was harvested, dissociated to single cells, and cultured in MGM-4 melanocyte growth medium (Lonza) + 200 nM 12-O-tetradecanoylphorbol-13-acetate (TPA; Sigma-Aldrich) + 5% FBS. Established cell line was then maintained in RPMI without TPA. Digital PCR was performed to confirm over 98% of cells were of human origin. A549, H358 and H2228 were gifts from the laboratory of Charles M. Rudin (MSKCC) and cultured in RPMI medium. 22RV1 was obtained from ATCC and cultured in RPMI medium. VCAP was a gift from Charles L. Sawyers laboratory (MSKCC) and cultured in high glucose DMEM medium. SW620, C106 and HCT116 were gifts from Rona Yaeger laboratory (MSKCC) and were cultured in RPMI, EMEM and F12/DMEM media respectively. CAMA-1 cell line was a gift from Sarat Chandarlapaty laboratory (MSKCC) and cultured in F12/DMEM medium. All media were supplemented with 10% FBS, 2 mM L-glutamine, 100 U/ml penicillin and 100 µg/ml streptomycin. All cells were cultured at 37 °C in 5% CO2 and tested negative for mycoplasma contamination by MycoAlert mycoplasma detection kit (Lonza). Expression of doxycycline-inducible shRNA was induced by supplementing media with 0.1-1 µg/ml doxycycline for 6 days, depending on the cell line. Cells that were transduced with lentivirus carrying puromycin-resistant plasmids were treated with 0.5-2 µg/ml puromycin, depending on the cell line. Cells that were transduced with lentivirus carrying neomycin-resistant plasmids were treated with 1000 µg/ml G418.

### siRNA transfection

SKMEL-23 and SKMEL-1128 cells were transfected with 25 nM and 50 nM siRNA respectively using lipofectamine RNAiMAX (Thermo Fisher Scientific). Silencer select negative control siRNA#1 (4390843) and #2 (4390846); Silencer siMITF#1 (AM51331) and #2 (4390824) were used. Cells were harvested 48 hr after transfection for immunoblot or ChIP-qPCR.

### Production of lentivirus

Lentivirus was produced based on published protocol with modifications^62^. Lenti-X 293T cells (Clontech) were maintained in high glucose DMEM supplemented with 10% FBS, 2 mM L-glutamine, 100 U/ml penicillin and 100 µg/ml streptomycin at 37 °C in 5% CO2. Low passage (<12) Lenti-X 293T cells were used for calcium phosphate precipitation to transfect packaging plasmids gag-pol and pMD.G [generous gifts from Yinon Ben-Neriah and Naama Kanarek (Hebrew University of Jerusalem)] and lentiviral transfer vector. Cells were incubated with calcium phosphate-precipitated DNA mixture at 37 °C in 3% CO2 overnight. Thereafter, cells were replenished with high glucose DMEM supplemented with 10% FBS, 2 mM L-glutamine, 100 U/ml penicillin and 100 µg/ml streptomycin and incubated at 37 °C in 5% CO2. Supernatant containing lentivirus was collected 48 and 72 hr post-transfection and filtered through a 0.45-μm PVDF membrane filter (Millipore). To infect cells, filtered lentiviral supernatant was incubated with cells in 8 µg/ml polybrene (Sigma-Alrich) containing media and spinoculation was performed at 500 xg for 45 min to increase transduction efficiency.

### Reverse transcription and quantitative PCR analysis (qPCR)

RNA was harvested and purified with homogenizer column and E.Z.N.A total RNA kit (Omega bio-tek). Reverse transcription was performed using high-capacity cDNA reverse transcription kit (Thermo fisher scientific). Real-time qPCR was performed using PowerUp SYBR green master mix (Thermo Fisher Scientific) and respective primers (**Supplementary Table 5**). Taqman primers and probes detecting *ALK* [against 5’UTR of *ALK*^ATI^; custom probes or *ALK* kinase domain; Hs00608286_gH, FAM-MGB (Thermo Fisher Scientific)] and *RPL27* [Hs03044961_g1, VIC-MGB (Thermo Fisher Scientific)] were co-amplified to determine the mRNA level of *ALK*. *ALK* or *RPL27* was first amplified singly to determine dynamic range and PCR efficiency and then co-amplified to confirm no interference. Subsequent experiments were performed with Taqman gene expression assay with *ALK* detecting kinase domain and *RPL27* due to higher PCR efficiency and dynamic range. To confirm the expression of *ALK*^ATI^ instead of full-length *ALK*, immunoblot and RNA-seq were performed. All qPCR was run on QuantStudio 6 flex real-time PCR system (Applied Biosystems). Relative abundance was calculated with delta-delta-Ct method. All qPCR products were sanger sequenced to confirm identity.

### Immunoblot

Immunoblot was performed as previously published^26^ using LI-COR Odyssey CLx scanner. Anti-MITF (Cell Signaling Technology, 12590), anti-SMC1A (Bethyl Laboratories Inc, A700-018) and anti-beta-actin (Cell Signaling Technology, 3700) were used as primary antibodies.

### Cloning

Tet-pLKO-puro (Addgene #21915) was used to express doxycycline-inducible shRNA. shRNAs were designed using GPP web portal (Broad Institute; **Supplementary Table 5**). Oligos were annealed and both oligos and vector were digested with AgeI and EcoRI (NEB). Lenti-sgRNA-neo (Addgene #104992) and lentiGuide-Puro (Addgene #52963) digested with BsmBI were used to express 2 sets of sgRNAs (**Supplementary Table 5**) to CRISPR delete CTCF sites. Digested oligos and vector were gel-purified with Qiagen gel purification kit and ligated by T4 DNA ligase (NEB). Ligate was transformed into Stbl3 competent bacteria (Thermo Fisher Scientific). Individual colonies were sanger sequenced to confirm identity. Absence of vector recombination was confirmed by digestion with AflII (Thermo Fisher Scientific) that has unique cut sites at the LTR regions.

### CRISPR interference (CRISPRi)

Cells were transduced with lentivirus expressing pHR-SFFV-KRAB-dCas9-P2A-mCherry (Addgene #60954) and sorted for mCherry positive cells. To target TSSs and enhancers, sgRNAs were designed using CRISPOR^63^ (**Supplementary Table 5**). To perturb CTCF binding, sgRNAs were designed using CRISPOR^63^ that target cognate CTCF binding motif (**Supplementary Table 5**). sgRNAs were cloned into a modified U6-based expression vector pSLQ1651-sgTelomere (F+E) (Addgene #51024) that enhanced sgRNA stability and assembly with dCas9 protein^64^. To remove mCherry reporter gene from the pSLQ1651-sgTelomere (F+E) vector, PCR was done using exCherry_NheI and exCherry_puro_EcoRI primers (**Supplementary Table 5**), with pSLQ1651-sgTelomere (F+E) as template. The PCR fragment and pSLQ1651-sgTelomere (F+E) were digested with NheI and EcoRI (NEB), ligated and transformed into Stbl3 competent bacteria (Thermo Fisher Scientific). sgRNA sequence was amplified using pSLQ1651-sgTelomere (F+E) vector with FE_AS and region specific sgRNA primer. A pool of 2-6 sgRNAs was used to target each region. PCR product was gel purified, digested with BstXI and XhoI (NEB) and inserted into pSLQ1651-sgTelomere (F+E) without mCherry. Identity of all plasmids was confirmed by sanger sequencing. Lentivirus carrying sgRNA was then transduced into KRAB-dCas9-P2A-mCherry expressing cells, selected by 2 µg/ml puromycin, and harvested after 6D to test for mRNA level or CTCF binding by ChIP-qPCR.

### RNA sequencing (RNA-seq)

Cells were lyzed directly on culture dish with TRIzol RNA isolation reagent (Thermo Fisher Scientific). Total RNA was extracted by thorough mixing with chloroform and isopropanol precipitation of upper aqueous phase. Total RNA precipitate was washed once with 75% ethanol, air dried and resuspended in RNase-free water. Absence of RNA degradation was confirmed by Agilent BioAnalyzer. 500 ng of total RNA was used for poly-A selection of RNA and library construction using TruSeq stranded mRNA LT Kit (Illumina, RS-122-2102). Samples were barcoded and run on a HiSeq 4000 in PE50 to generate 40-50 M reads. Ribosomal reads represented 0.75-1.87% of the total reads generated and the percent of mRNA bases averaged 80%.

### Cap analysis of gene expression sequencing (CAGE-seq)

CAGE-seq was performed according to Takahashi et al.,^65^ protocol with modifications to detect 5’ends of capped RNAs. 5 µg total RNA was reverse transcripted using a primer that carried a 15-nt random region and EcoP15I recognition sequence and SuperScriptIII RNase H-reverse transcriptase (Thermo Fisher Scientific) in the presence of trehalose/sorbitol to increase efficiency of reverse transcription. Diol groups on RNA/cDNA duplex were oxidated and labelled with biotin hydrazide (Vector laboratories). RNase I (Promega) treatment was performed to cleave single-stranded RNA regions that were not hybridized with first-strand cDNA. tRNA-coated streptavidin beads (Thermo Fisher Scientific) were used to capture biotinylated RNA/cDNA duplex. Sample was washed three times with wash buffer A [4.5 M NaCl, 50 mM EDTA (pH 8.0) and 0.1 % Tween20], once with wash buffer B [10 mM Tris–HCl (pH 8.5), 1 mM EDTA (pH 8.0), 0.5 M NaOAc (pH 6.1) and 0.1 % Tween20] and once with wash buffer C [0.3 M NaCl, 1 mM EDTA (pH 8.0) and 0.1 % Tween20]. Single-stranded cDNAs were released from streptavidin beads by heating at 95 °C for 5 min. Released cDNA was treated with RNase H (Thermo Fisher Scientific) and RNase I (Promega) at 37 °C for 45 min and proceeded for library construction. Library was constructed with modified primers for compatible sequencing on Illumina Nextseq 500 sequencer. Ligation of bar-coded 5’ linker was performed. Second-strand cDNA was synthesized using Deep Vent (exo-) DNA polymerase (NEB) and biotinylated second-strand primer. cDNA was treated with shrimp alkaline phosphatase (Thermo Fisher Scientific) to remove phosphate group from 5’ linker. EcoP15I (NEB) was used to cleave 27 nt downstream of the recognition site at the 5’ linker. 3’ linker was ligated to the 27-nt-long tags, purified with tRNA-coated streptavidin beads and amplified with 2x Phusion high-fidelity PCR mix (NEB). CAGE libraries were treated with Exonuclease I (NEB) to degrade any single-stranded DNA before sequencing on NextSeq 500 using SR50 to obtain around 40 M reads for each library. All primers used for CAGE-seq were listed in **Supplementary Table 5**.

### Enzymatic methyl sequencing (EM-seq)

High-molecular-weight genomic DNA was extracted using Puregene cell core kit (Qiagen) and sent to MSKCC Epigenetics Research Innovation Lab for processing. EM-seq was performed with 100ng of purified genomic DNA using NEBNext® Enzymatic Methyl-seq Kit (NEB, E7120) and following manufacturer’s instructions. After quantification and size distribution assessment using a Qubit Flex fluorometer (Thermo Fisher Scientific) and a TapeStation (Agilent), respectively, the purified libraries were sequenced on an Illumina NovaSeq 6000 to generate around 200 M PE100 reads per library.

### Chromatin immunoprecipitation (ChIP) and ChIP-seq

ChIP was performed as previously described^26,66^. 20 x 10^6^ cells were harvested and fixed in 1% formaldehyde for 10 min with gentle shaking. Glycine was added to final concentration of 125 mM and incubated for 5 min. Cells were washed once with PBS, resuspended in LB1 [50 mM HEPES (pH 7.5), 140 mM NaCl, 1 mM EDTA, 10% glycerol, 0.5% NP-40, 0.25% Triton X-100 and 1× complete EDTA-free protease inhibitor] and incubated with rotation at 4 °C for 10 min. Samples were pelleted, resuspended in LB2 [10 mM Tris-HCl (pH 8.0), 200 mM NaCl, 1 mM EDTA, 0.5 mM EGTA and 1× complete protease inhibitor] and incubated rotating at 4 °C for 10 min. Nuclei were pelleted, resuspended in LB3 with 0.5% SDS [0.5% SDS, 20 mM Tris-HCl (pH 8.0), 150 mM NaCl, 2 mM EDTA, 1% Triton X-100 and 1× complete protease inhibitor] and sonicated using Covaris E220 focused ultrasonicator (peak power = 150 W, duty factor = 2%, cycle per burst = 200) for 12 min. Soluble chromatin extracts were cleared by centrifugation and 1% was kept as input DNA. Sheared chromatin was diluted to contain 0.16% SDS for antibody binding and incubated overnight at 4 °C with 2 ug anti-H3K27Ac (Abcam, ab4729), anti-H3K4ME3 (Cell Signaling Technology, 9751S), anti-SMC1A (Bethyl laboratories inc, A300-055A), anti-MITF (Active motif, 91201) or CTCF (Cell signaling technology, 3418S). Spike-in chromatin (Active motif, 53083) and antibody (Active motif, 61686) were included to normalize SMC1A ChIP-seq. 20 ul Pierce ChIP-grade magnetic beads were used for precipitation at 4 °C for 6 hr. Beads were washed 3x LiCl buffer [250 mM LiCl, 10 mM Tris-HCl (pH 8.0), 1% sodium deoxycholate, 1% NP-40 and 1 mM EDTA], 1x RIPA wash buffer [500 mM LiCl, 50 mM HEPES-KOH (pH 7.6), 0.7% sodium deoxycholate, 1% NP-40 and 1 mM EDTA] and 1x TE [10 mM Tris-HCl (pH 8.0) and 1 mM EDTA] + 50 mM NaCl. Chromatin was eluted in elution buffer [1% SDS, 50 mM Tris-HCl (pH 8.0) and 10 mM EDTA] and incubated at 65 °C for 30 min. Eluate was reverse cross-linked at 65 °C overnight, treated with RNaseA (Roche) at 37 °C for 1 hr and Proteinase K (Roche) at 55 °C for 2 hr before recovered using a miniElute PCR purification kit from Qiagen. ChIP-qPCR was performed using primers listed in **Supplementary Table 5** and PowerUp SYBR green master mix (Thermo Fisher Scientific). Data were presented and normalized by percent input. For ChIP-seq, library was prepared using KAPA HTP library kit (KAPA biosystems). ChIP libraries were sequenced to obtain 30-40 M reads for each sample on HiSeq 2500 or 4000 with either SR50 or PE50.

### CUT&RUN

CUT&RUN was performed using CUT&RUN assay kit (Cell Signaling Technology) based on published protocol by Skene P. et al.^32^. Briefly, 0.25 x 10^6^ live cells were immobilized on activated Concanavalin A beads and permeabilized with digitonin. Cell:bead suspension was washed with 1x wash buffer supplemented with spermidine and protease inhibitor cocktail and incubated with 1 µg of antibody at 4 °C overnight. The following antibodies were used: anti-H3K4me1 (Thermo Fisher Scientific/Invitrogen, 710795), anti-H3K4me3 (Thermo Fisher Scientific/Invitrogen, PA5-27029), anti-H3K27ac (Abcam, ab4729), anti-H3K27me3 (Cell Signaling Technology, 9733BF), anti-H3K9me3 (Abcam, ab176916), anti-H3K36me3 (Active motif, 61021) and anti-CTCF (Cell Signaling Technology, 3418F) antibodies. Antibody-targeted digestion of chromatin was performed by incubation of pAG-MNase at 4 °C for 1 hr and activation by calcium chloride at 4 °C for 30 min. MNase cleavage was stopped by addition of 1x stop buffer. Digested chromatin was allowed to diffuse out of cells at 37 °C for 20 min and DNA was purified by Nucleospin gel and PCR clean-up spin column (Takara bio). Sequencing libraries were prepared using the KAPA HTP Library Preparation Kit (KAPA Biosystems). Barcoded libraries were run on NovaSeq 6000 in a PE100 run. An average of 25 M reads were generated per sample.

### Hi-C

Arima Hi-C kit (Arima genomics) was used for all Hi-C experiments. Standard input protocol was followed with 3 x 10^6^ cells for each condition. Proximally ligated DNA was fragmented to around 400bp by Covaris E220 focused ultrasonicator (peak power = 140 W, duty factor = 10%, cycle per burst = 200) for 70 sec. Fragmented DNA was sized selected to have a distribution between ∼200-600bp using AMPure XP beads (Beckman Coulter). ∼ 2µg of size-selected DNA was used for library preparation using KAPA hyper prep kit (KAPA biosystems) and TruSeq DNA unique dual indexes (Illumina). The number of PCR cycle used for library amplification was determined by KAPA library quantification kit for Illumina platform (KAPA biosystems). Each Hi-C library was sequenced to obtain 500M reads PE100 with biological duplicates to achieve a total of 1 B reads for each condition.

### Circular chromosome conformation capture sequencing (4C-seq)

4C-seq was performed in reference to Krijger P. et al.^45^ paper with modifications. In brief, 10 x 10^6^ cells were fixed with 1% formaldehyde for 10 min. Cross-linking was quenched with 200 mM glycine and cells were lysed with NP-40 lysis buffer [10 mM Tris-HCl (pH 8.0), 10 mM NaCl, 0.2% NP-40 and 1x complete EDTA-free protease inhibitors (Roche)] for 20 min on ice. Nuclei were incubated with 0.5 % SDS at 65 °C for 10 min followed by 1.25 % Triton X-100 at 37 °C for 15 min. Nuclei were digested with 500 U EcoRI (*ALK*^ATI^ viewpoint) or NlaIII (*ULK4* intron 31 viewpoint) overnight. qPCR was performed using primers that span restriction enzyme cut site at the bait versus uncut control region to confirm the efficiency of restriction enzyme digestion to over 90%. Digested chromatin was ligated at 16 °C overnight, reverse cross-linked and purified. For efficient PCR amplification and next generation sequencing, a second round of restriction enzyme digestion was performed using 10 µg 3C product and 100 U DpnII (for both *ALK*^ATI^ and *ULK4* intron 31 viewpoints) at 37 °C for 5 hr. Sample was ligated overnight at 16 °C, ethanol precipitated at −80 °C and purified with QIAquick PCR purification column. Inverse PCR was performed using 4C_ALKATI_EcoRI or 4C_ULK4int31_NlaIII (reading) and 4C_ALKATI_DpnII or 4C_ULK4int31_DpnII (non-reading) primers with 2x Phusion high-fidelity PCR mix (NEB). Each 4C sample was amplified in 4 PCR reactions, each containing 200 ng of 4C DNA template for 18 cycles. A second round of PCR was performed to add P5 and P7 adapters as well as index sequences for multiplex sequencing. Double-sided size selection was performed using AMPure XP beads (Beckman coulter) with 0.5x and 1x of the PCR volume. Sequencing was performed on Illumina NovaSeq 6000 sequencer PE75. Each sample was sequenced to generate 10 M reads. All primers used for 4C-seq were listed in **Supplementary Table 5**.

### HiChIP

HiChIP was performed based on published protocol^67^. 15 x 10^6^ cells were cross-linked with 1% formaldehyde for 10 min. Fixation was quenched with 125 mM glycine. Cells were washed once with PBS and lysed for 30 min using in situ Hi-C lysis buffer [10mM Tris-HCl (pH 7.5), 10mM NaCl, 0.2% NP-40 and 1x complete EDTA-free protease inhibitors (Roche)]. Nuclei were treated with 0.5% SDS followed by 1.25% Triton X-100 before digesting with 400 U MboI (NEB) for 4 hr. Restriction enzyme digested overhangs were filled in with biotin-dATP (Thermo Fisher Scientific) and ligated at 16 °C overnight. Nuclei were sonicated in nuclear lysis buffer [50mM Tris-HCl (pH 7.5), 10mM EDTA, 0.5% SDS and 1x EDTA-free protease inhibitor (Roche)] using Covaris E220 with the following parameter: peak power = 140 W, duty factor = 5%, cycle per burst = 200 and time = 4 min. Pull-down with anti-H3K27ac antibody (Abcam, ab4729) was performed by diluting sheared chromatin to 0.16% SDS in buffer with Pierce ChIP-grade magnetic beads. ChIP reaction was washed 3x LiCl buffer [250 mM LiCl, 10 mM Tris-HCl (pH 8.0), 1% sodium deoxycholate, 1% NP-40 and 1 mM EDTA], 1x RIPA wash buffer [500 mM LiCl, 50 mM HEPES-KOH (pH 7.6), 0.7% sodium deoxycholate, 1% NP-40 and 1 mM EDTA] and 1x TE [10 mM Tris-HCl (pH 8.0) and 1 mM EDTA] + 50 mM NaCl. Chromatin was eluted in 50 mM sodium bicarbonate (pH 8.0) with 1% SDS and reverse cross-linked with proteinase K at 65’C overnight. Streptavidin C1 beads (Thermo Fisher Scientific) were used to capture biotinylated DNA. Beads were washed twice with tween wash buffer [5 mM Tris-HCl (pH 7.5), 0.5mM EDTA. 1M NaCl and 0.05% Tween 20]. To build library for Illumina NovaSeq 6000 sequencing, Tn5 (Illumina) was used for tagmentation and amplified with Nextera Ad1_noMX and Ad2.X primers (**Supplementary Table 5**) using 2x Phusion high-fidelity PCR mix (NEB). AMPure beads (Beckman coulter) were used for double-sided size selection (0.5x and 1x of the PCR volume) and purification of library DNA that sequenced with PE75 500M reads.

### RNA-seq analysis

RNA-seq reads were trimmed and filtered for quality using Trimmomatic (v0.38). Processed reads were then aligned to hg19 using STAR (v2.7.1a). For each GENCODE (v19) gene, reads were counted using featureCounts (v1.6.4). Gene counts were used to perform differential analysis using DESeq2 (v1.22.2). TPM normalized bigWig tracks were generated using deepTools (v3.1.10 bamCoverage with ‘--normalizeUsing BPM’) for each replicate. BedGraph files from each replicate, converted from bigWig using bigWigToBedGraph, were combined using bedtools (v2.27.1) unionbedg and scores were averaged for each condition. The averaged bedGraph files were converted to TPM averaged bigWig tracks using bedGraphToBigWig (v4). TPM counts were generated with StringTie (v2.1.1) using BAM files and GENCODE (v19) gtf.

### CAGE-seq analysis

CAGE-seq reads were analyzed based on the Nextflow CAGE-seq analysis pipeline^68^. Index sequences, EcoP15I recognition site and the first G following the EcoP15I site at the 5’-end from reads were removed using Cutadapt (v2.6). rRNA sequences were filtered out using SortMeRNA (v4.2.0) and filtered reads were aligned to the hg19 genome using STAR (v2.7.1a). At this step, two sets of BAM files were created for each sample: one with uniquely mapped reads (adding --outFilterMultimapNmax 1 to STAR command) and one with multi-mapped reads (adding --outFilterMultimapNmax 10 to STAR command). Unmapped reads from BAM files were removed. BAM files were converted to BED files with the summed up 1 bp unclustered CAGE tags using SAMtools (v1.9) and bamtobed (v.2.26.0). We pooled CAGE tags across samples, filtered them by TPM threshold of 0.5 and then clustered them with paraclu^69^ (minValue of 30) to generate a BED file with the clustered CAGE transcription start sites (CTSSs). Then, paraclu-cut with default settings (remove single-position clusters, clusters longer than 200 bp, clusters with maximum density/baseline density less than 2, and any cluster contained in a larger cluster) was used to filtered out clusters. Promoter width was calculated by the distance between CAGE tags for each clustered CTSS with a 200 bp cut-off using CAGEr (v1.33.1). CAGEr analyzed promoter width across all samples by considering both the position and the CAGE tag signal at TSSs along the tag cluster. The width of every tag cluster is calculated based on the cumulative distribution of the CAGE signal along the cluster CAGE tags were intersected with the clusters identified by paraclu and a raw count table with columns for each sample and rows for each tag cluster was generated. DESeq2 (v1.30.1) was used to perform differential CAGE expression using the raw count table. CTSS clusters were annotated using R/Bioconductor packages ChIPseeker (v1.26.2) and AnnotationDbi (v1.52) using GENCODE (v19) gene annotations. The annotatePeak function from ChIPseeker package was used to assign peaks to genomic annotation including promoter (±1kb around TSS), exon, 5’ UTR, 3’ UTR, intronic, intergenic, and downstream (downstream of gene end) and calculate the distance of a peak to the nearest gene.

### Correlation between RNA-seq and CAGE-seq

Spearman correlation was calculated between either log2TPM or log2FoldChange in RNA-seq for each gene and CAGE-seq for the corresponding promoter CTSS.

### Repetitive region analysis

Multi-mapped CTSSs were compared to all the repeatmasker annotated repetitive elements by the genomic coordinates downloaded from the UCSC browser (hg19). An overlap of at least 1 bp was used to transfer the names of repeat class, family, and subfamily to each CTSS.

### ChIP-seq analysis

ChIP-seq was analyzed according to the ENCODE 3 pipeline as previously published^70^. ChIP-seq reads were aligned to hg19 using BWA (v0.7.17-r1188). Unmapped and low MAPQ reads (< 30) were filtered out using SAMtools (v1.9) and PCR duplicates were removed using Picard tools (v2.18.16). Depth normalized bigWig tracks were generated using deepTools (v3.1.1) bamCoverage with ‘bamCoverage -b ${input_bam} -o ${output_bigwig} --normalizeUsing RPGC --ignoreForNormalization chrX chrM -bl hg19-blacklist.v2.bed’. Peak calling was performed using MACS2 (2.2.7.1) with SMC1A ChIP-seq. deepTools (v3.5.1) was used to examine read density at the SMC1A peaks or differentially expressed CTSSs and +/-5kb from the centers of the respective regions.

### CUT&RUN analysis

All CUT&RUN data were processed as described for ChIP-seq to generate bam files. Replicates were merged using the pysam python module. The merged bam files were then converted to bigWig files with deepTools (v3.1.1) bamCoverage (binSize as 10, RPGC normalization, and other parameters as default). Density plot was generated using deepTools as in ChIP-seq.

### ChromHMM

ChromHMM (v1.24) was used to partition the human genome into 15 states based on the CUT&RUN-seq data from H3K4me1, H3K4me3, H3K27ac, H3K27me3, H3K9me3, H3K36me3, and CTCF; and ChIP-seq from SMC1A. The two biological replicates were analyzed separately. The OverlapEnrichment function was used to determine enrichment of the upregulated and downregulated CTSSs in the 15 states.

### CG content analysis

All CTSSs were expanded to 300bp from the center and then used to count the percentages of C, G, and CG in the sequences with both strands considered. Due to the short length of CTSSs, an expansion of 300bp was made to get an accurate determination of the local CG composition. The analysis was also tested on 100-bp and 500-bp expansion and produced similar results.

### EM-seq and DNA methylation analysis

EM-sequencing reads were processed using Bismark pipeline^71^. Raw reads with low-quality (less than 20) and adapter sequences were removed by Trim Galore (v0.6.4). C(G) in the trimmed sequences was converted to T(A) and mapped to similarly converted reference human genome (hg19), lambda, and pUC19 separately using default Bowtie2 settings implemented in Bismark. Duplicated reads were discarded. The remaining alignments were then used for cytosine methylation calling by Bismark methylation extractor. Batch effects were adjusted using limma (v3.46.0). Any CpGs with coverage less than 10 were removed prior downstream analyses. After filtering, 969,316 CpG loci were recovered with an average coverage of 19x across all samples.

### HOMER motif analysis

HOMER motif analysis was performed to discover both *de novo* and known motifs using the ’findMotifsGenome.pl’ script (v 4.11) from the HOMER software suite. The analysis was configured with a size parameter of 200, using hg19 genome sequences. CTSS was used as the background.

### 4C-seq analysis

R1 was used for analysis. From the single-end reads, non-reading primers at the 3’ end were trimmed using Cutadapt (v3.7). A maximum error-rate of 0.15 was allowed and trimmed reads shorter than 40 bp were discarded. 5’ end of the trimmed reads was further trimmed until the primary restriction enzyme site (RE1; EcoRI or NlaIII) using R. Only reads containing RE1 next to the viewpoint were considered for downstream analysis. Bowtie2 (v2.4.5) was used for aligning the trimmed reads to the human genome (hg19) while SAMtools (v1.14) (parameter: -q 1 -bSu) was used to filter for high-quality uniquely mapped reads. Fragment maps of the hg19 genome were generated for each restriction enzyme pair. Blind fragments (which have RE1 on both ends) were excluded from the analysis. Aligned reads were assigned to overlapping fragment ends. Fragment read counts were depth normalized to per M reads on the cis chromosome where the viewpoint is located. The top 3 fragments with the highest read counts corresponding to the undigested and self-ligated fragments (usually the ones closest to the viewpoint) were excluded when calculating depth normalization. For smoothening, running mean across the fragment read counts was calculated using a sliding window sized 21 (averaging 10 fragments left and right to the fragment of interest). For statistical analysis, mean normalized read counts were calculated for 25 kb (*ALK* intron19) or 10 kb (*ULK4* intron 31) bins with a 1 kb shift. With an assumption that the measured read counts follow normal distribution for each bin, a two-tailed paired t-test was done for each replicate to call significantly differential bins between conditions using a *P*-value cutoff of 0.05. All 4C-seq data was visualized using Gviz (1.41.1)^72^ and ggplot2 (v3.4.4) package in R.

### FunSeq2 analysis

Normalized RNA-seq for TCGA-SKCM samples was downloaded from NCI GDC Data Portal. *ALK* expression was ranked, and the top *ALK*-expressing tumors were further examined (RSEM >210). Tumors with *ALK* amplification were excluded (n=3). Exon quantification data for *ALK* was analyzed and high *ALK*^ATI^-expressing tumors were defined as more than 10-fold differential expression in *ALK* exons 20-29 than to exons 1-19^26^. The top 50 cases of high *ALK*-expressing tumors that met these criteria were assigned as high *ALK*^ATI^-expressing tumors and used to compare to the bottom 50 cases with low/no *ALK*^ATI^ expression. FunSeq2^46^ analysis for *NIPBL* was run on default settings and each mutation was assigned a FunSeq2 score of 1 to 5 with a higher score that prioritized somatic alterations with functional significance. A composite FunSeq2 score for *NIPBL* was calculated by combining FunSeq2 score for each *NIPBL* mutation in the same tumor sample. Tumor sample that has no *NIPBL* mutation was assigned FunSeq2 score of 0.

### Hi-C and HiChIP analysis

#### Generation of valid paired-end reads

Hi-C sequencing files were processed with HiC-Pro pipeline (v3.0.0). In silico digestion of the hg19 genome by ‘Arima’ restriction enzymes was performed with ‘digest_genome.py’ tool from HiC-Pro, and bed files generated were used for assigning mapped reads to DNA fragments. All aligned filtered deduplicated read pairs from each replicate were used to calculate cis valid paired-end read distance after normalizing for sequencing depth in R.

#### Pre-processing of Hi-C and HiChIP data

Hi-C or HiChIP reads were aligned to the hg19 genome using the Juicer pipeline (v1.7.6) with default settings. MAPQ30 threshold .hic files were used for the subsequent analyses.

#### Calling significant and differential loops

HiC-DC+^73^ was run on both pooled and individual replicates of Hi-C and HiChIP data to find significant interactions at 5, 10 or 25 kb resolutions. We defined TSS interactions as the ones with one anchor overlapping a promoter. HiC-DC+ differential interaction calling (function hicdcdiff) was also performed at different resolutions for pairwise comparison. HiC-DC+ normalized scores (observed over expected counts) were used to show Hi-C interactions in plots. To generate H3K27ac HiChIP hotspots, virtual 4C plots were calculated as the maximum −log10(*P-value*) of the interactions of regions with the *ALK*^ATI^ alternative promoter in intron 19 as anchor.

#### TAD annotations

TADs were annotated using OnTAD (v1.3)^19^ on ICE normalized and MAPQ30 threshold Hi-C data at 10 kb resolution. A penalty score of 0.1 and maximum TAD size of 2 Mbp were set while running OnTAD. Singleton TADs have no inner sub-TADs. TADs with nested sub-TADs were assigned a hierarchical TAD level according to the highest level of sub-TADs contained within a nested TAD.

Each TAD was annotated with the overlapping CAGE-seq CTSSs. The relative CTSS distance to TAD boundaries was calculated by dividing the distance of the CTSS to the nearest TAD boundary by the length of the respective TAD.

The insulation score for each TAD was obtained from OnTAD outputs.

## Data availability

All next-generation sequencing data were deposited at the Gene Expression Omnibus (GEO) under accession code GSE232303. Secure token slsveygyrjudzsp can be used to access the data while they remain private before publication.

## Code availability

No custom code package or newly developed algorithm was generated in this study. All analysis was performed using standard settings unless otherwise described in Materials and Methods.

## Extended Data Figure Legends

**Extended Data Figure 1.**
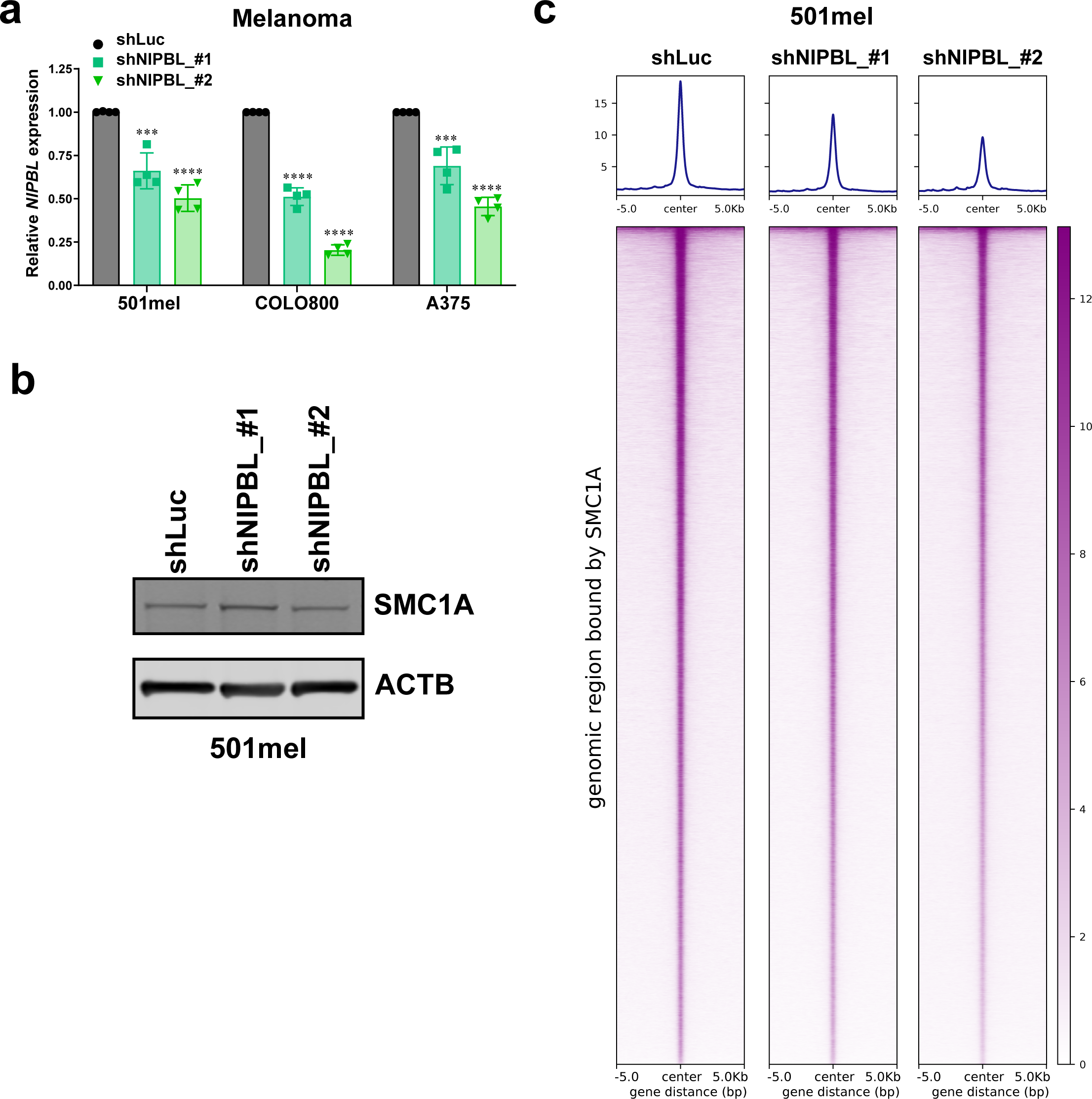
Efficiency of *NIPBL* downregulation by dox-inducible shRNAs and their effect on cohesin complex chromatin distribution. **a**, Relative *NIPBL* mRNA levels by dox-inducible shRNA-mediated downregulation (shNIPBL_#1 and shNIPBL_#2) in various melanoma cell lines, 501mel, COLO800 and A375. Error bars are mean ± S.D., n=4 biological replicates. \*\*\**P*<0.001, \*\*\*\**P*<0.0001, ordinary one-way ANOVA Dunnett’s multiple comparisons test. **b,** Immunoblots of SMC1A and control actin (ACTB) in whole cell lysate with shNIPBL knockdown. **c**, Representative density plots of normalized genome-wide ChIP-seq profile of SMC1A, demonstrating partial loss of NIPBL resulted in the reduction of the cohesin complex core subunit, SMC1A, on chromatin in 501mel cells. Data from one representative biological replicate (n=3) are shown.

**Extended Data Figure 2.**
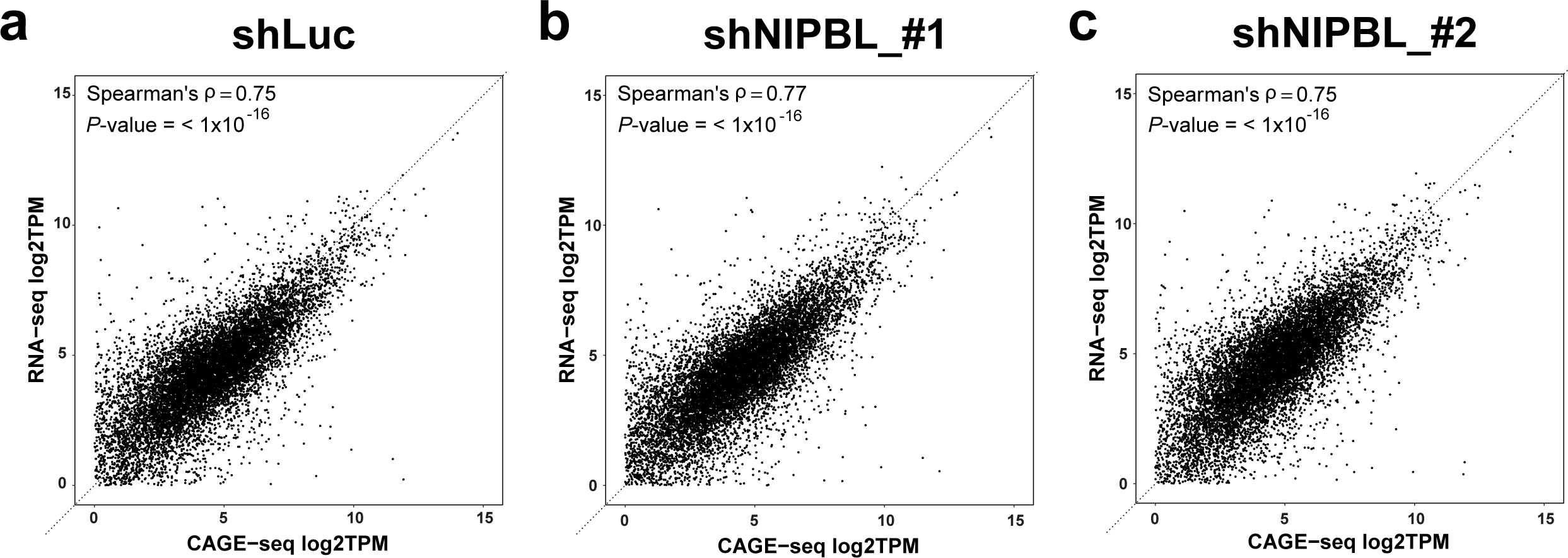
Spearman correlation of annotated transcripts’ expression between whole transcriptome analysis by poly-A RNA-seq and CTSS analysis by CAGE-seq under shLuc (control) (a), shNIPBL_#1 (b) and shNIPBL_#2 (c) *NIPBL* perturbation conditions.

**Extended Data Figure 3.**
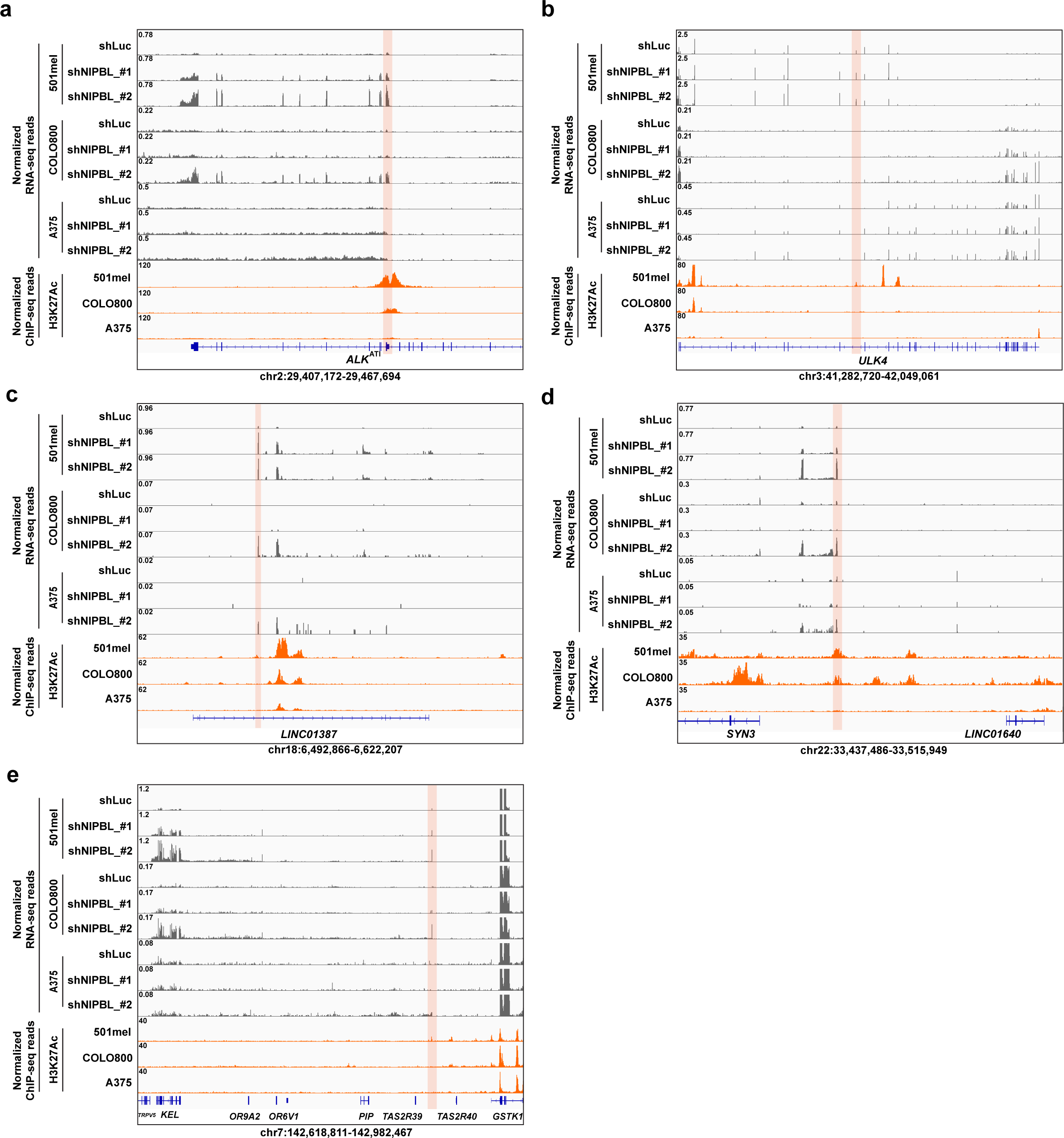
Examples of *NIPBL* loss-induced alternative promoter usage in intron and distal intergenic regions in melanoma cell lines. a-e, Normalized RNA-seq and H3K27ac ChIP-seq profiles of selective examples of activated alternative promoters (shaded in pink) with *NIPBL* perturbation (shNIPBL_#1 and shNIPBL_#2 compared to shLuc controls) in 501mel, COLO800 and A375. Examples included *ALK*^ATI^ intron 19 (**a**), *ULK4* intron 31 (**b**), *LINC01387* intron 2 (**c**); distal intergenic regions, such as between *SYN3* and *LINC*01640 in chr22:33,468,501-33,471,517 (**d**) and chr7:142,895,998-142,896,596 (**e**).

**Extended Data Figure 4.**
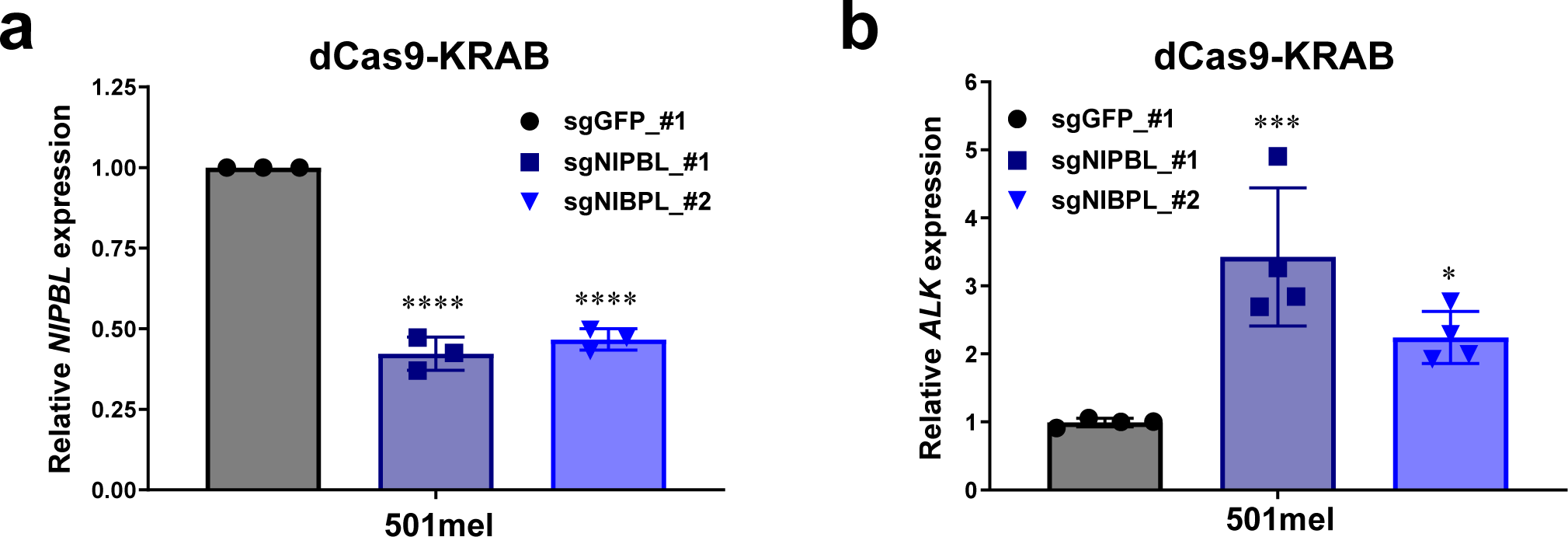
Efficiency of *NIPBL* downregulation by CRISPRi and their impact on *ALK*^ATI^ expression. **a,** Normalized *NIPBL* mRNA expression by *NIPBL*-specific CRISPRi perturbation. **b**, The effect of CRISPRi-mediated downregulation of *NIPBL* on *ALK*^ATI^ mRNA level in 501mel melanoma cells. Error bars are mean ± S.D., n= 3-4 biological replicates. \**P*<0.05, \*\*\**P*<0.001, \*\*\*\**P*<0.0001, ordinary one-way ANOVA Dunnett’s multiple comparisons test.

**Extended Data Figure 5.**
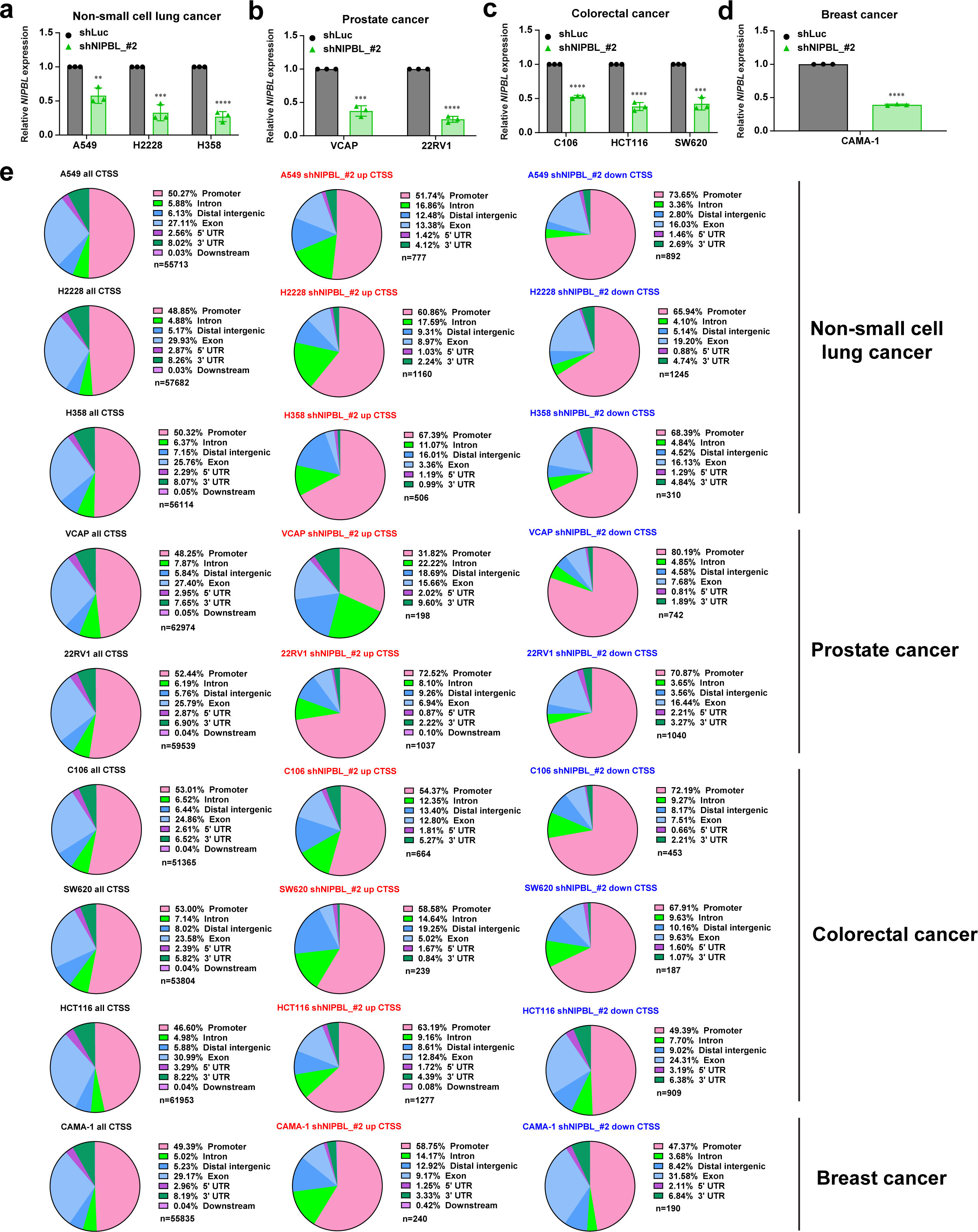
The impact of *NIPBL* downregulation by dox-inducible shRNA on CTSS by CAGE-seq in various cancer cell lines. **a-d**, Efficiency of *NIPBL* downregulation by doxycycline-inducible shRNA in non-small cell lung cancer (*KRAS*-mutant: A549 and H358; *ALK*-fusion-positive: H2228) (**a**), prostate cancer (AR-positive: VCAP and 22RV1) (**b**), colorectal cancer (C106, HCT116, and SW620) (**c**) and breast cancer (ER-positive, HER2-negative: CAMA-1) (**d**) cell lines. Error bars are mean ± S.D., n=3 biological replicates \*\**P*<0.01, \*\*\**P*<0.001, \*\*\*\**P*<0.0001, two-tailed unpaired t-test. **e**, Genomic annotation of all, significantly (FDR<0.1) upregulated (log2FoldChange>1) and downregulated (log2FoldChange<-1) CTSS by CAGE-seq with shNIPBL_#2-mediated *NIPBL* downregulation in indicated cell lines. Percentage of each genomic feature is indicated.

**Extended Data Figure 6.**
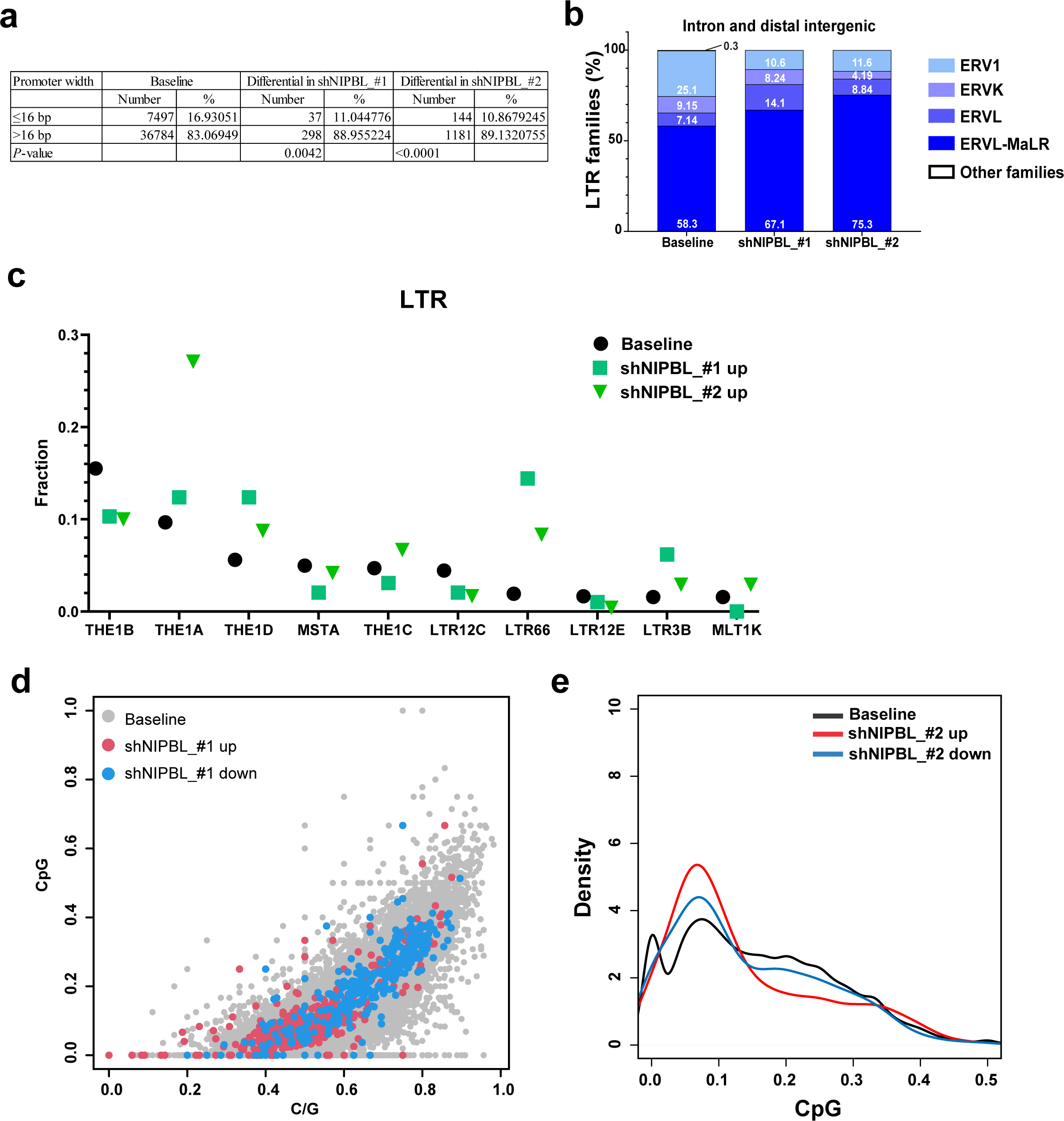
CTSS bimodal distribution significance and characteristics and distribution of transposable elements and subclasses in differentially regulated CTSSs by *NIPBL* downregulation. **a**, The number and corresponding percentage of CTSSs with promoter width lower or equal to 16 bp (sharp promoter) or higher than 16 bp (broad promoter) in baseline and shNIPBL perturbation. *P*-value by two-tailed chi square test. **b-c,** Distribution of LTR families (**b**) and subtypes (**c**) for baseline and upregulated CTSSs by shNIPBL. **d,** CpG and CG composition in baseline CTSSs and differentially regulated CTSSs by shNIPBL_#1. **e,** CpG compositions of significantly upregulated, downregulated and all CTSSs by shNIPBL_#2.

**Extended Data Figure 7.**
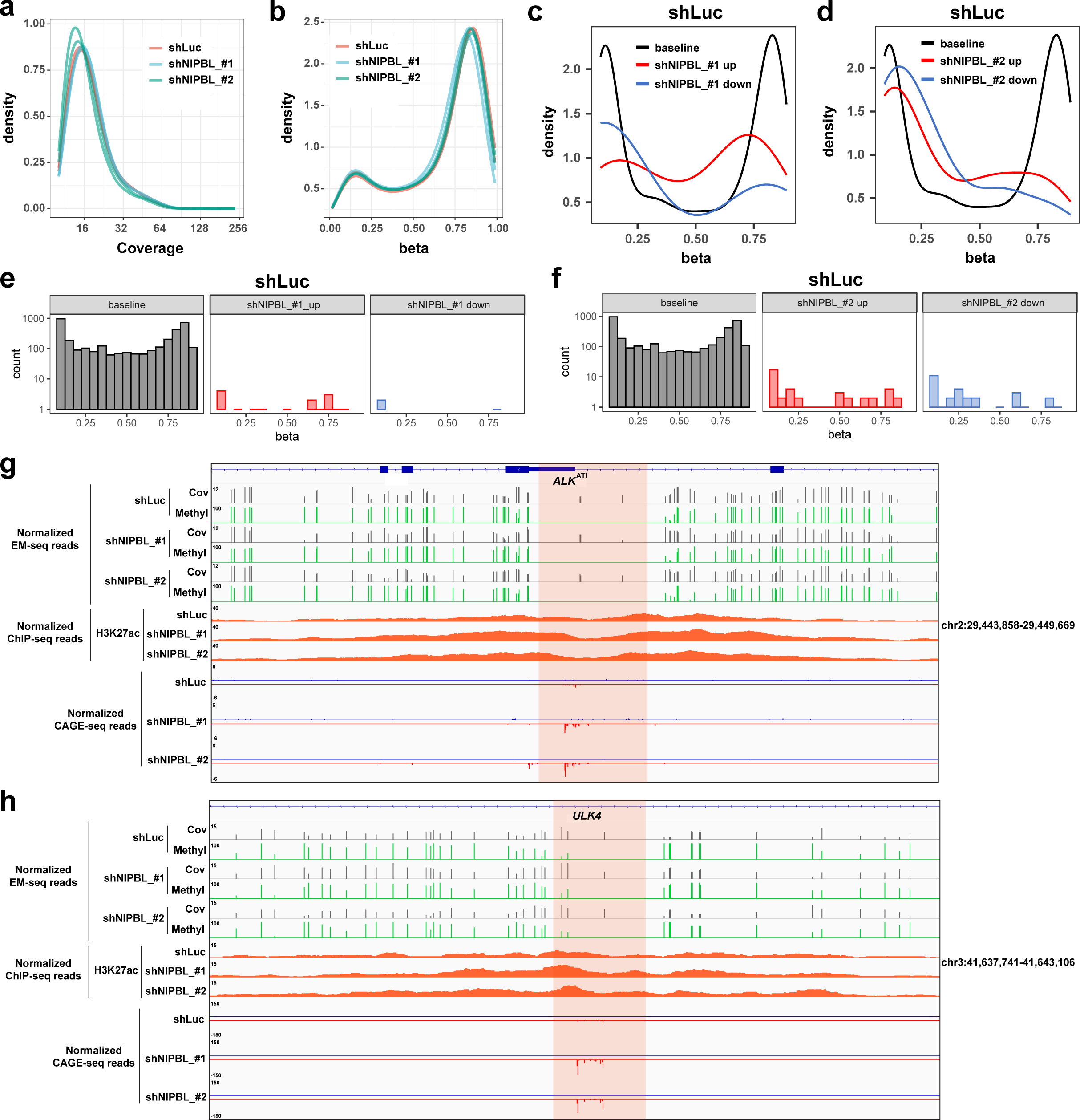
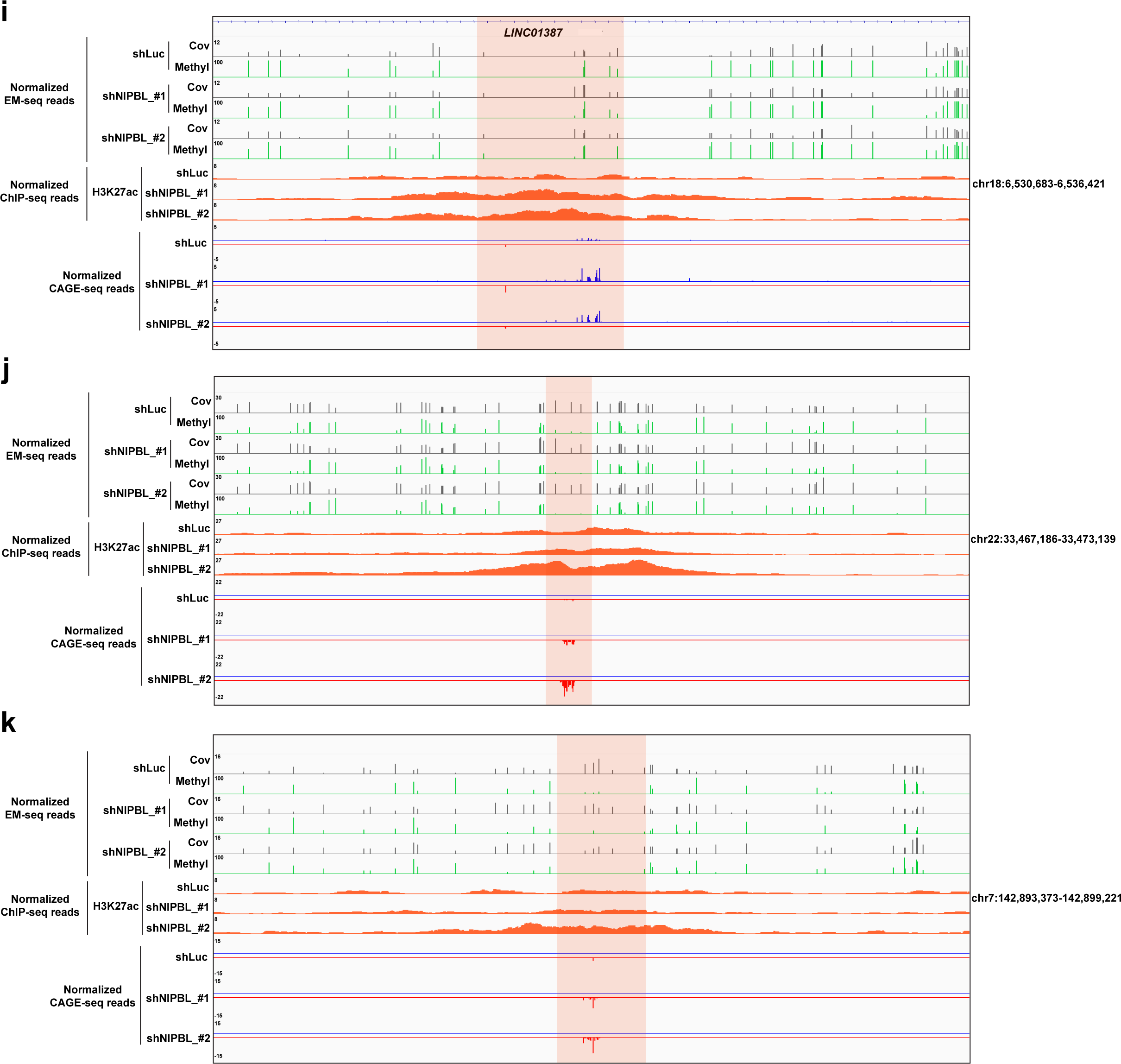
Global DNA methylation analysis by enzymatic methyl sequencing (EM-seq) **a-b**, Representative distribution plots of DNA methylation sequencing coverage (**a**) and global DNA methylation levels (**b**) with *NIPBL* perturbation (shNIPBL_#1 shNIPBL_#2) and control (shLuc). n=2 biological replicates. **c-d**, Representative DNA methylation density plots under shLuc condition for baseline CTSSs and differentially regulated CTSSs by shNIPBL_#1 (**c**) and shNIPBL_#2 (**d**) perturbations. **e-f**, Histogram of CpG methylated regions under shLuc condition for baseline CTSSs and differentially regulated CTSSs by shNIPBL_#1 (**e**) and shNIPBL_#2 (**f**). **g-k**, Representative DNA methylation profiles by EM-seq of significantly upregulated CTSSs in *ALK* intron 19 alternative promoter region (**g**), *ULK4* (intron 31) (**h**), *LINC01387* (intron 2) (**i**), *SYN3* (intergenic regions) (between *SYN3* and *LINC01640*) (**j**) and *KEL* (intergenic regions) (between *TAS2R39* and *TAS2R40)* (**k**). Coverage (Cov) and methylation conversion (Methyl) tracks were shown.

**Extended Data Figure 8.**
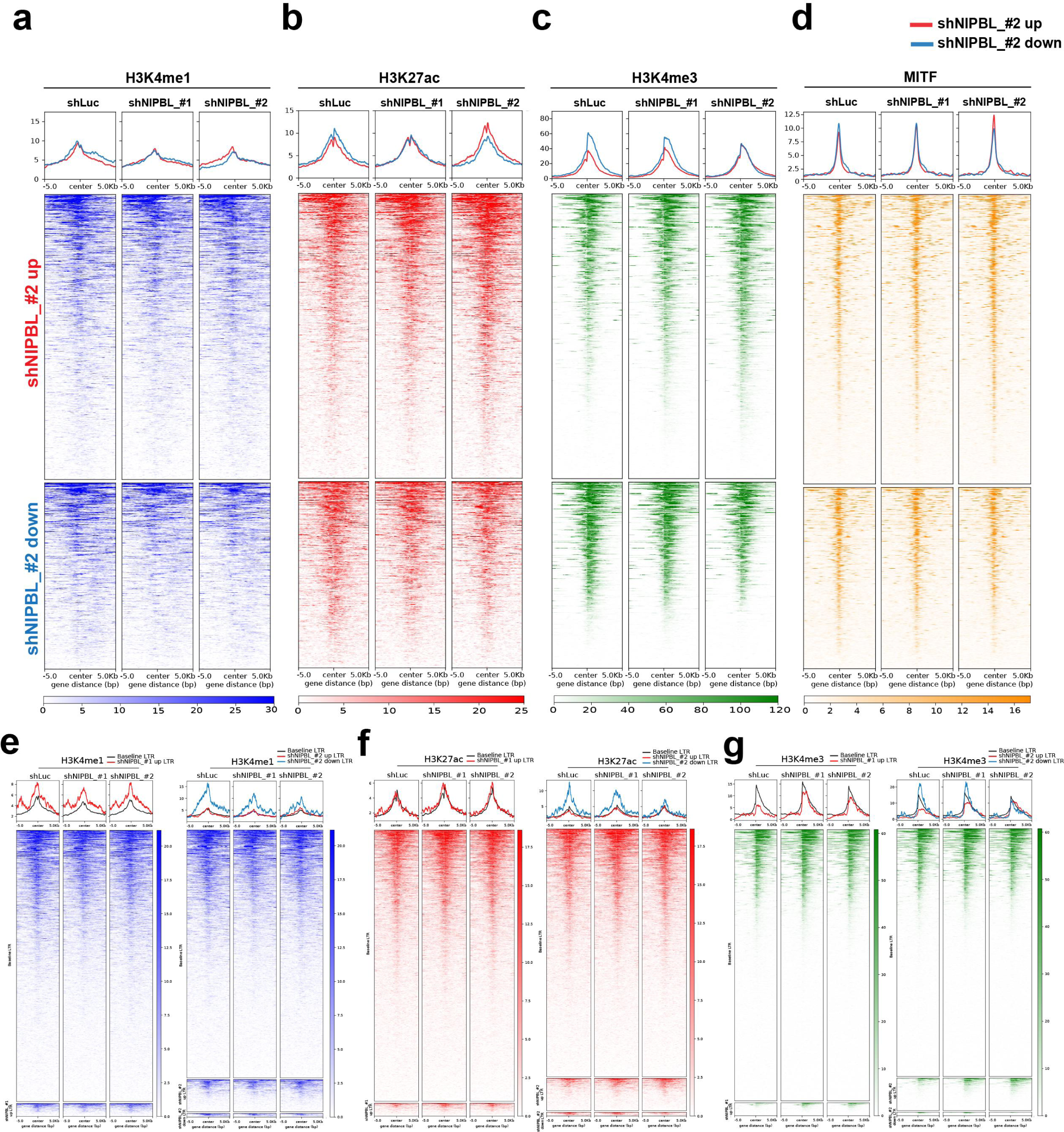
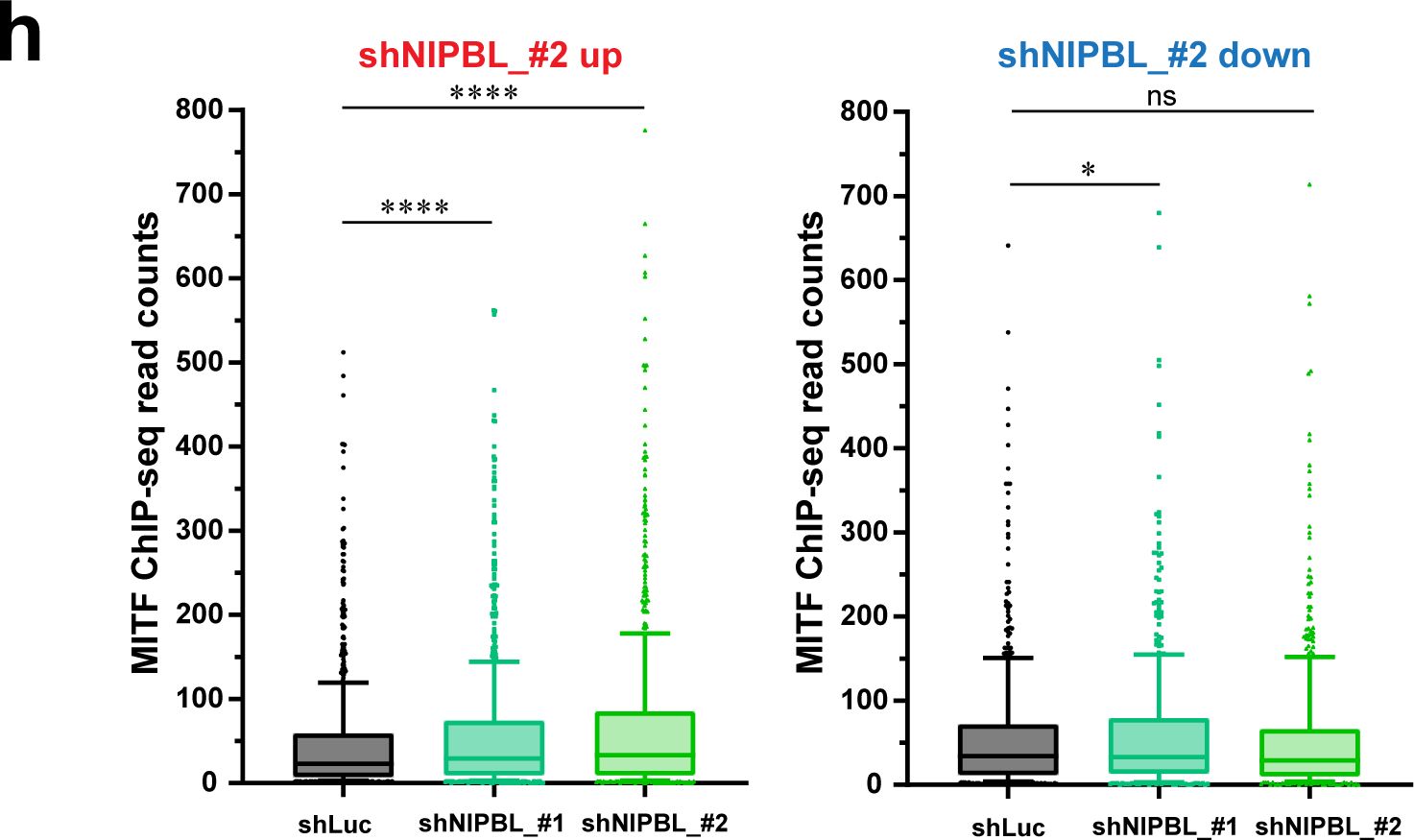
Genome-wide distribution of indicated chromatin marks under *NIPBL* perturbation conditions. a-c, Density plots of normalized genome-wide profile of H3K4me1. (**a**), H3K27ac (**b**), H3K4me3 (**c**) by CUT&RUN and MITF (**d**) by ChIP-seq centered on differentially regulated CTSSs with shNIPBL_#2 perturbation. Data from one representative biological replicate (n=2) were presented. **e-g**, Density plots of genome-wide distribution of H3K4me1 (**e**), H3K27ac (**f**), H3K4me3 (**g**) by CUT&RUN, centered on baseline, differentially upregulated and downregulated LTR-derived CTSSs by shNIPBL. Data from one representative biological replicate (n=2) are shown. There were few LTR-derived CTSSs downregulated by shNIPBL_#1 (n=4) and therefore they were not included in the density plots. **h,** Box and whiskers plots showing 10-90 percentile of MITF binding by ChIP-seq read counts at differentially upregulated (left panel) and downregulated (right panel) CTSSs by shNIPBL. Each dot represented quantification of MITF binding at one differentially regulated CTSS locus. ns, not significant; \**P*<0.05, \*\*\*\**P*<0.0001, matched one-way ANOVA Dunnett’s multiple comparisons test.

**Extended Data Figure 9.**
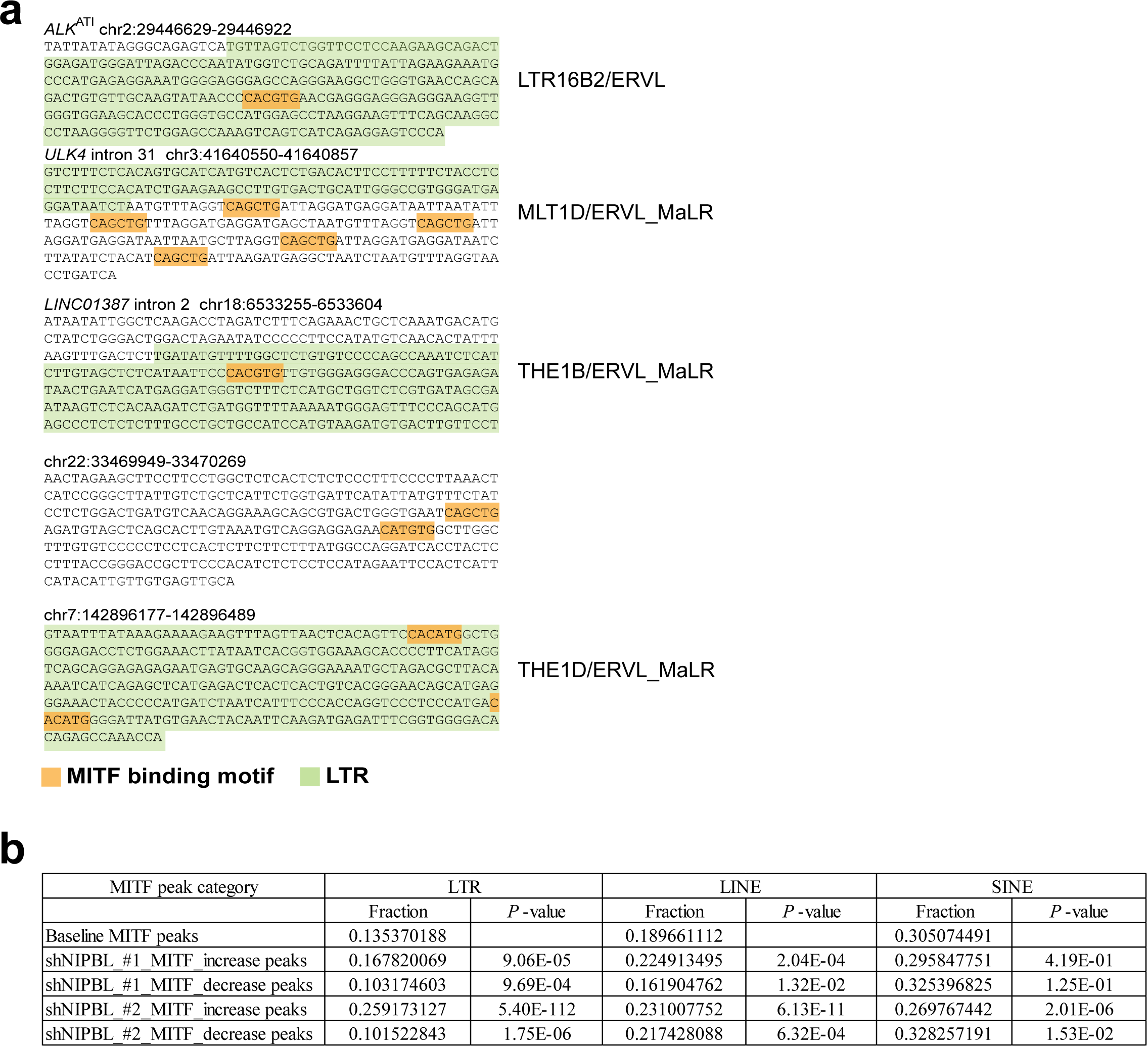
MITF binding at activated alternative promoters in intron and distal intergenic regions and the fraction that overlapped with transposable elements by *NIPBL* perturbation in melanoma cells. **a**, MITF binding motif (highlighted in orange) at representative activated alternative promoters often embedded within LTR sequence (highlighted in green). **b,** Fraction of baseline or shNIPBL-differentially increased/decreased MITF ChIP-seq peaks that were localized within LTR, LINE and SINE repetitive elements in the genome. E, exponents of 10. *P-*value by proportion test.

**Extended Data Figure 10.**
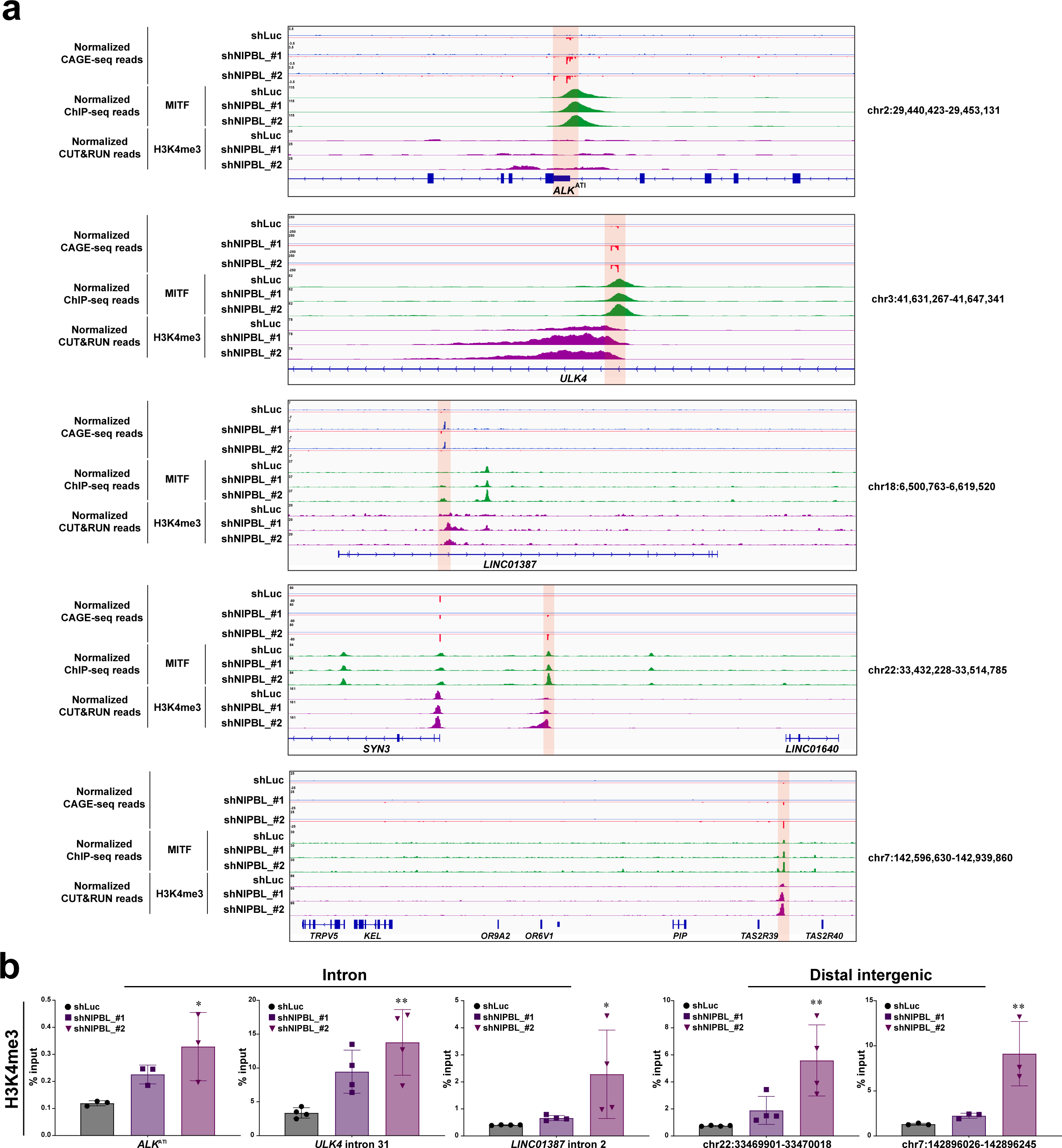
Recruitment of MITF to activated alternative promoters is associated with increased enrichment of H3K4me3 with *NIPBL* loss. **a**, Normalized CAGE-seq, MITF ChIP-seq, and H3K4me3 CUT&RUN profiles with *NIPBL* perturbation at representative activated alternative promoters (shaded in pink). **b,** H3K4me3 enrichment by ChIP-qPCR with *NIPBL* perturbations at selective activated alternative promoters. Data indicated the mean ± S.D. (n=3-4 biological replicates). \**P*<0.05, \*\**P*<0.01, ordinary one-way ANOVA Dunnett’s multiple comparisons test.

**Extended Data Figure 11.**
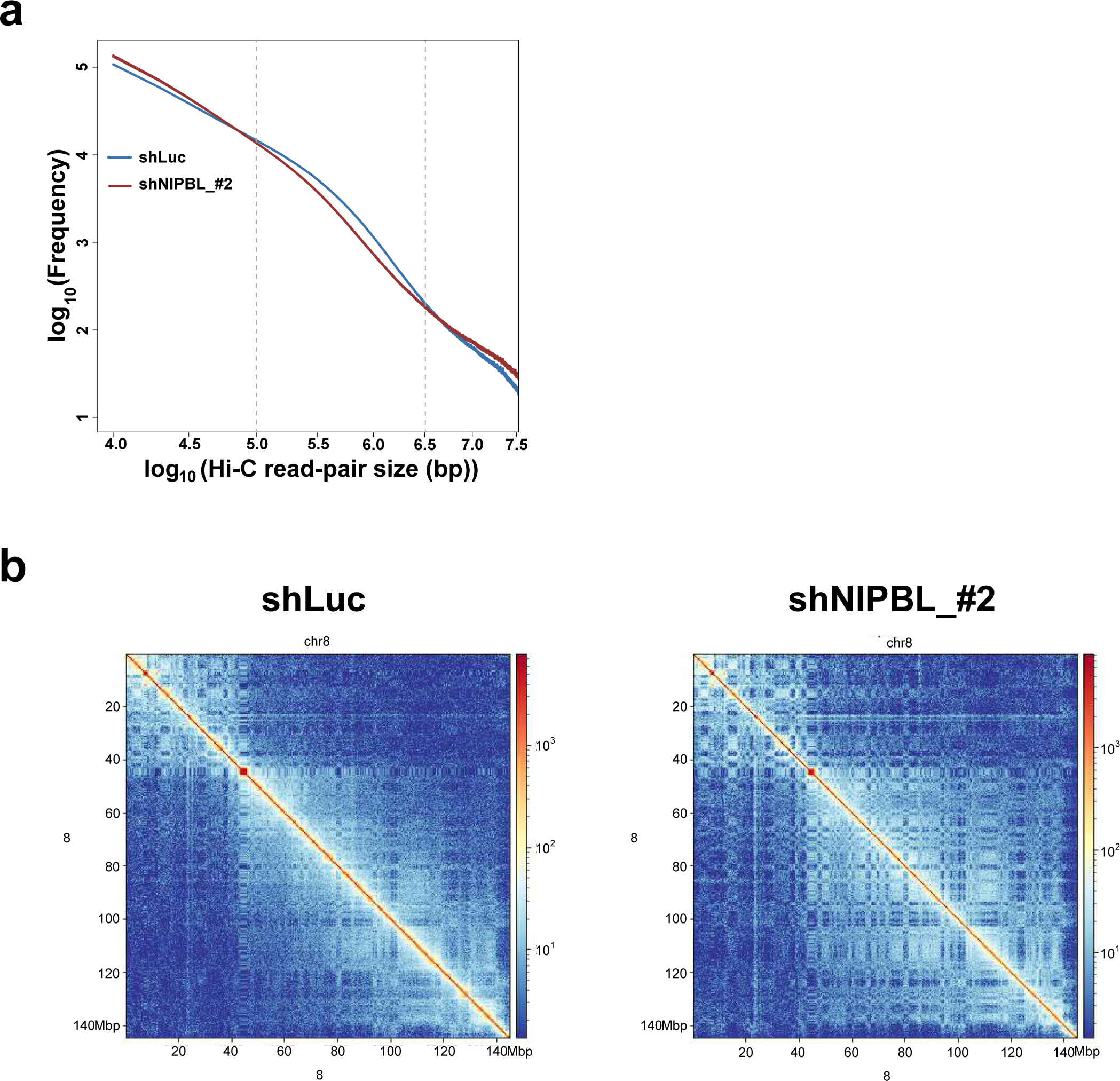
*NIPBL* downregulation leads to alterations in long-range interaction frequency and chromatin architecture. (**a**) Hi-C contact probability plot showing the average contact frequency of valid paired-end reads as a function of genomic distance with *NIPBL* perturbation. Genomic distance between valid cis read pairs was calculated and the distribution of log10 read pair genomic distance was plotted. (**b**) Hi-C contact map (at 100 kb resolution) of a representative region (chr8) with *NIPBL* downregulation (shNIPBL_#2, right panel) vs. control (shLuc, left panel), demonstrating finer segregation in compartments after *NIPBL* downregulation.

**Extended Data Figure 12.**
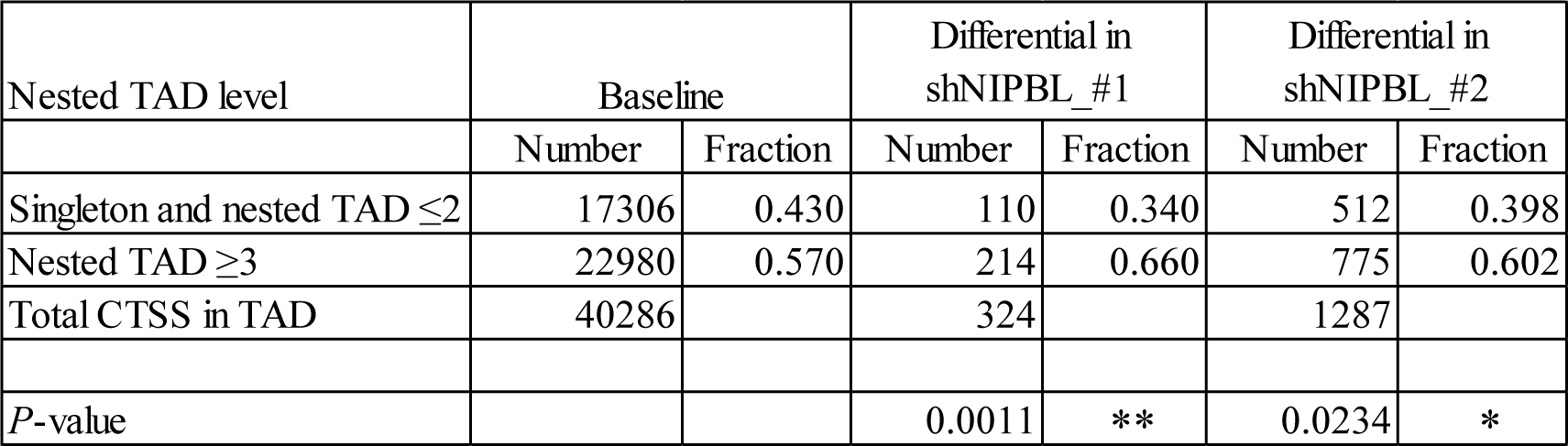
Number and fraction of CTSSs in hierarchical and non-hierarchical TADs defined under control condition (≥ level 3 vs. ≤ level 2 and singleton) \**P*<0.05, \*\**P*<0.01, two-tailed chi square test.

**Extended Data Figure 13.**
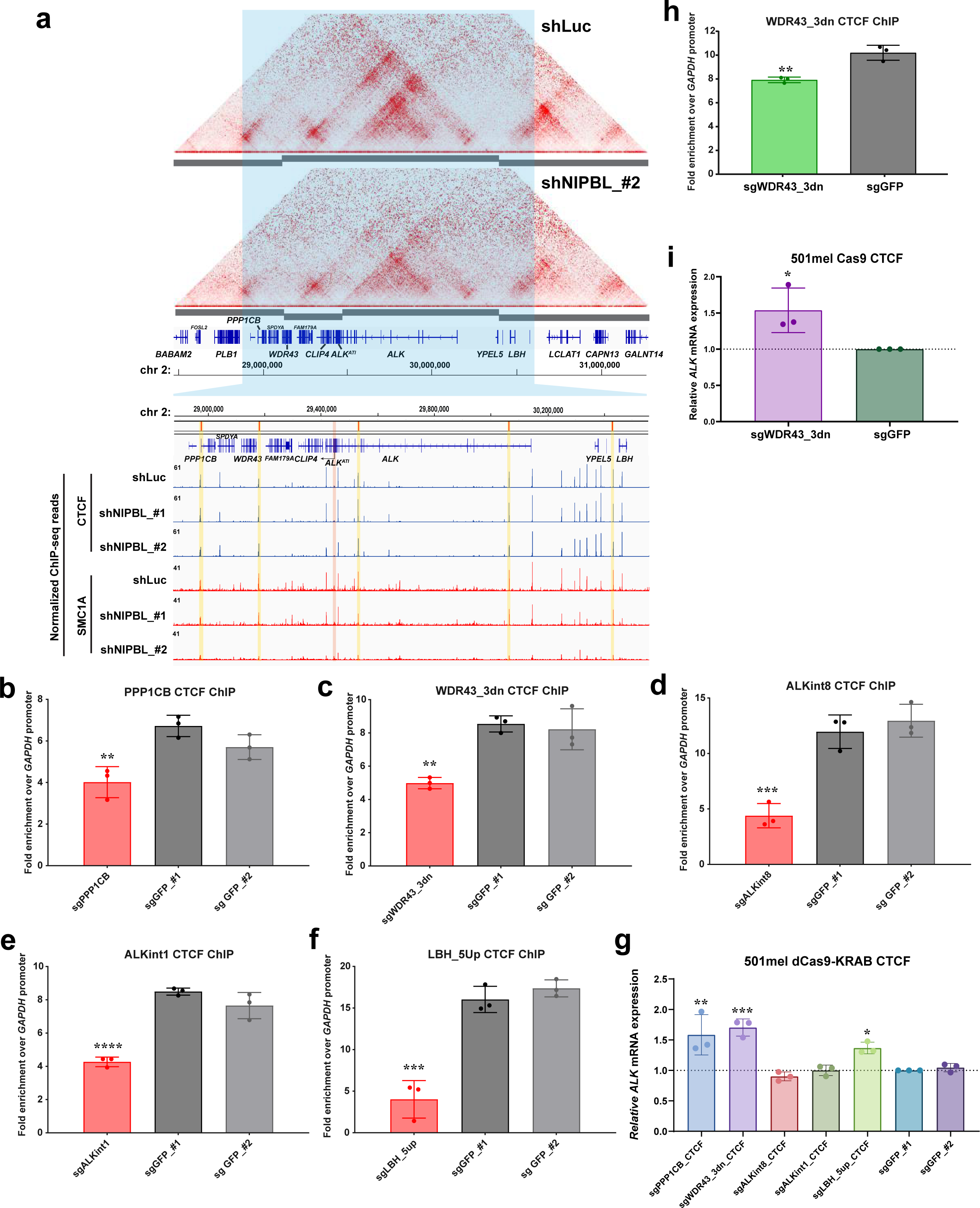
Perturbation of CTCF insulators in proximity to *ALK*^ATI^ alternative promoter. **a,** Hi-C contact map (10 kb resolution) of TADs surrounding *ALK*^ATI^ in control (top, shLuc) and *NIPBL* knockdown (bottom, shNIPBL_#2). Genomic region of interest around the *ALK*^ATI^ was shaded in blue. Selected CTCF insulator sites (shaded in yellow) that overlapped with SMC1A binding around the *ALK*^ATI^ alternative promoter (shaded in pink) were targeted. **b-f**, CTCF ChIP-qPCR with specific sgRNA targeting selective CTCF binding by dCas9-KRAB compared to sgGFP1/sgGFP2 controls. **g**, Relative *ALK*^ATI^ mRNA levels with dCas9-KRAB and respective sgRNAs. Error bars are mean ± S.D. (n=3 biological replicates). \**P*<0.05, \*\**P*<0.01, \*\*\**P*<0.001, **** *P*<0.0001, ordinary one-way ANOVA Dunnett’s multiple comparisons test (**b-g**). **h-i,** Quantification of CTCF binding by ChIP-qPCR at WDR43_3dn (**h**) by Cas9 deletion of CTCF and relative *ALK*^ATI^ mRNA levels (**i**). Error bars are mean ± S.D. (n=3 biological replicates). \**P*<0.05, \*\**P*<0.01, two-tailed unpaired t-test (**h, i**).

**Extended Data Figure 14.**
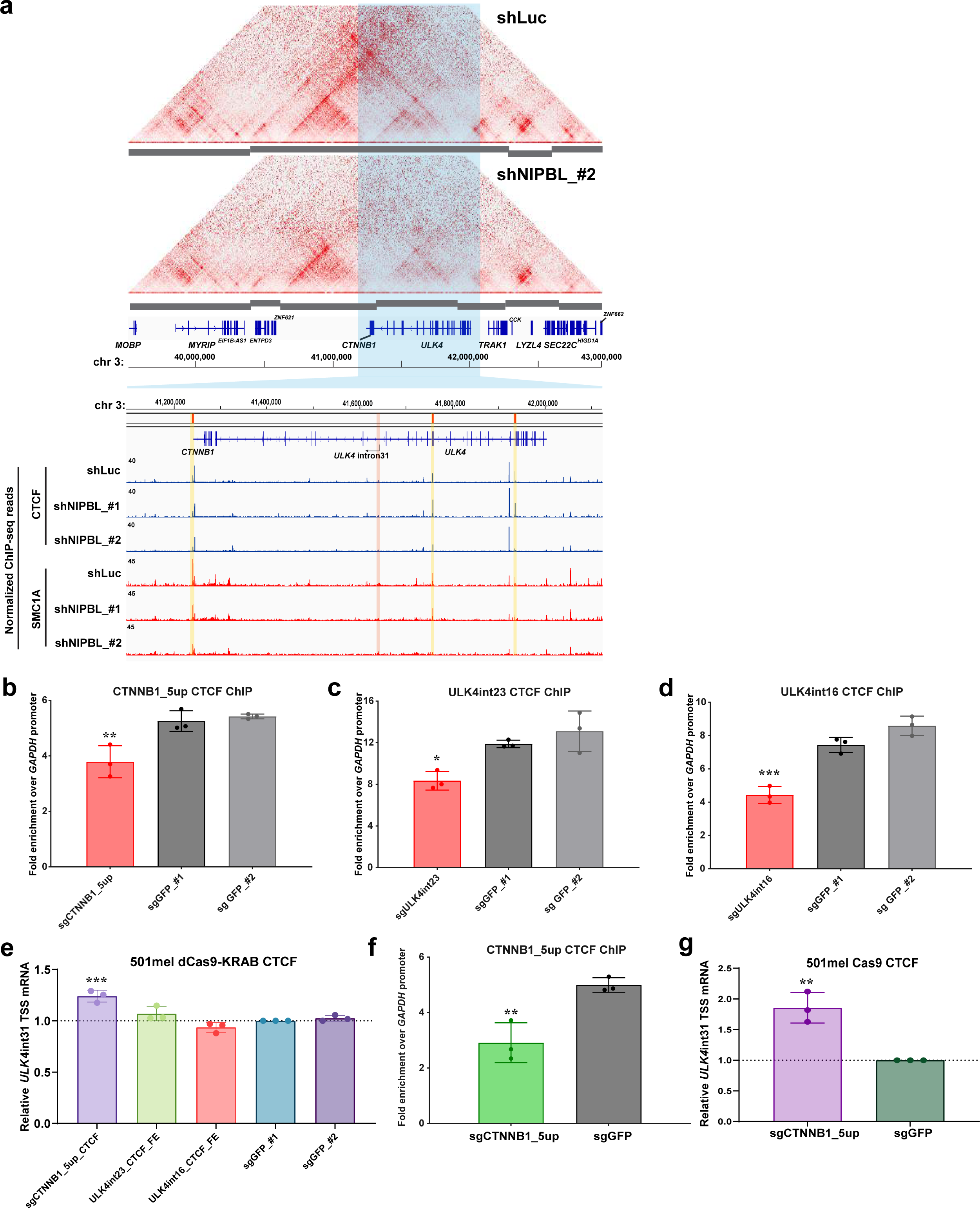
Perturbation of CTCF insulators in proximity to *ULK4* intron 31 alternative promoter. **a,** Hi-C contact map (10 kb resolution) of TADs surrounding *ULK4* intron 31 in control (top, shLuc) and *NIPBL* knockdown (bottom, shNIPBL_#2). Genomic region of interest around the *ULK4* alternative promoter was shaded in blue. Selected CTCF insulator sites (shaded in yellow) that overlapped with SMC1A binding around the *ULK4* intron 31 alternative promoter (shaded in pink) were targeted. **b-d**, CTCF ChIP-qPCR with specific sgRNA targeting selective CTCF binding by dCas9-KRAB compared to sgGFP1/sgGFP2 controls. **e**, Relative *ULK4* intron 31 TSS mRNA levels with dCas9-KRAB and respective sgRNAs. Error bars are mean ± S.D. (n=3 biological replicates). \**P*<0.05, \*\**P*<0.01, \*\*\**P*<0.001, ordinary one-way ANOVA Dunnett’s multiple comparisons test (**b-e**). **f-g,** Quantification of CTCF binding by ChIP-qPCR at CTNNB1_5up (**f**) by Cas9 deletion of CTCF and relative *ULK4* intron 31 TSS mRNA levels (**g**). Error bars are mean ± S.D. (n=3 biological replicates). \*\**P*<0.01, two-tailed unpaired t-test (**f, g**).

**Extended Data Figure 15.**
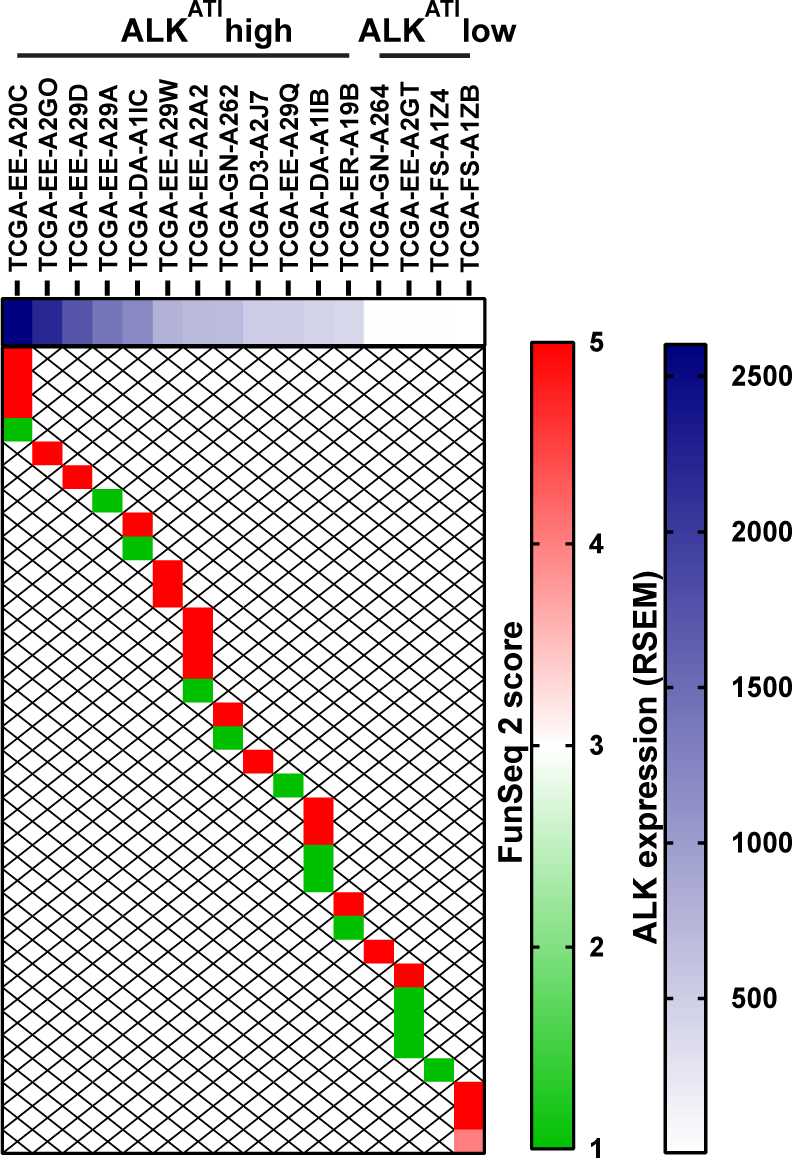
*NIPBL* functional perturbation by FunSeq2 of TCGA melanoma cases with high and low *ALK*^ATI^ expression (RSEM) Each cohort of high and low *ALK*^ATI^ expression has 50 cases. *NIPBL* mutations are present in 12 of 50 cases in high *ALK*^ATI^ expression cohort and in 4 of 50 cases of low *ALK*^ATI^ expression cohort. FunSeq2 score is considered 0 for cases without any *NIPBL* mutations.

**Extended Data Figure 16.**
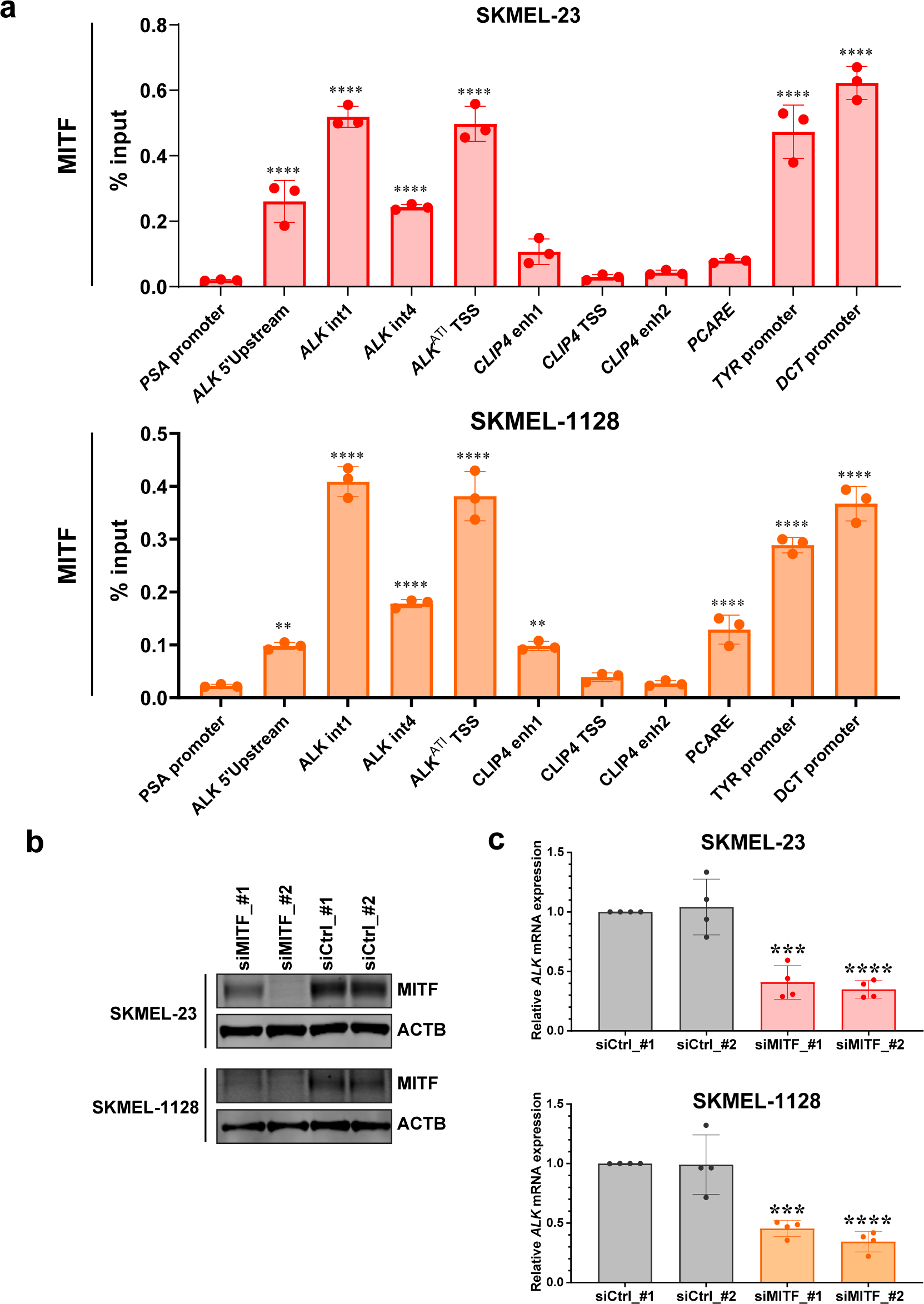
MITF binding is required for *ALK*^ATI^ transcriptional activation in patient-derived melanoma cell lines. **a**, Quantification of MITF binding by ChIP-qPCR at selective H3K27ac-enriched regions that interacted with *ALK*^ATI^ alternative promoter identified by H3K27ac HiChIP in *ALK*^ATI^-expressing melanoma cell lines. Negative control: *PSA* promoter; positive controls: *TYR* and *DCT* promoters. **b,** Immunoblots of MITF and control actin (ACTB) with siRNA-mediated knockdown of *MITF* in melanoma cells. **c,** Relative *ALK*^ATI^ mRNA levels with siRNA-mediated *MITF* knockdown in melanoma cells as in **b**. Error bars are mean ± S.D. (n=3-4 biological replicates). \*\**P*<0.01, \*\*\**P*<0.001, **** *P*<0.0001, ordinary one-way ANOVA Dunnett’s multiple comparisons test.

## Supplementary Table Legends

**Supplementary Table 1. Differential gene expression analysis for poly-A RNA-seq by DESeq2.**

**Supplementary Table 2. Differential CTSS analysis for CAGE-seq by DESeq2.**

**Supplementary Table 3. Genomic annotation of CTSSs and differential CTSS analysis by CAGE-seq with shRNA-mediated *NIPBL* downregulation in various cancer cell lines.**

**Supplementary Table 4. Homer motif analysis at differentially regulated CTSSs at non-promoter regions.**

**Supplementary Table 5. Primer list.**

## Acknowledgements

We want to thank Juan Li, Neeman Mohibullah, Cassidy Cobbs and Agnes Viale from Memorial Sloan Kettering Cancer Center Integrated Genomics Operation (IGO) for their excellent technical support. Cailian Liu for collecting and extracting RNA from patient-derived melanoma cell lines for screening *ALK*^ATI^-expressing cell lines.

This work was supported by grants from the NIH/NCI (R01 CA228216, DP2 CA174499, R01 CA280657, U01 CA252048 and P50 CA217694), Department of Defense (W81XWH-15-1-0124 and W81XWH-22-1-0326), Francis Collins Scholar Neurofibromatosis Therapeutic Acceleration Program, Geoffrey Beene Cancer Research Fund and Cycle for Survival Linn Family Discovery Fund to P. Chi; NIH/NCI grants (R01 CA208100-04, U54 CA224079-03, P50 CA092629-20 and R01 CA265026) to Y. Chen. The IGO core was funded by the NCI Cancer Center Support Grant (P30 CA08748), Cycle for Survival, and the Marie-Josée and Henry R. Kravis Center for Molecular Oncology.

## Authors’ Disclosures

Y. Chen reports other support from Oric Pharmaceuticals and grants from Foghorn outside the submitted work. P. Chi reports personal fees from Deciphera, grants and nonfinancial support from Deciphera, Pfizer/Array, and Ningbo NewBay, and personal fees from Zai Lab and Novartis outside the submitted work. No disclosures were reported by the other authors.

## Authors contributions

E.W.P.W.: conceptualization, resources, data curation, formal analysis, validation, visualization, investigation, project administration, writing-original draft, review & editing. M.S.: conceptualization, software, data curation, formal analysis, visualization, investigation, project administration, writing-original draft review & editing. R.Y.: data curation, formal analysis, visualization, writing-review & editing. U.L.: data curation, formal analysis, visualization, writing-review & editing. Y.A.Z.: formal analysis, visualization, writing-review & editing. R.M.: visualization, writing-review & editing. F.T.: data curation, validation, writing-review & editing. N.A.: data curation, formal analysis, visualization, software, writing-review & editing. A.P.: formal analysis, visualization, writing-review & editing. C.J.L.: resources, writing-review & editing. T.T.: data curation, writing-review & editing. A.M.F.: data curation, writing-review & editing. T.W.: resources, writing-review & editing. A.R.: data curation, writing-review & editing. S.C.: data curation, writing-review & editing. C.L.: data curation, writing-review & editing. M.L.: data curation, writing-review & editing. T.M.: resources, writing-review & editing. P.J.H.: supervision, writing-review & editing. R.K.: supervision, writing-review & editing. E.K.: supervision, writing-review & editing. E.A.: supervision, writing-review & editing. D.Z.: data curation, formal analysis, visualization, supervision, writing-review & editing. Y.C.: conceptualization, supervision, funding acquisition, writing-review & editing. C.S.L.: conceptualization, supervision, writing-review & editing. P.C.: conceptualization, supervision, funding acquisition, writing-original draft, review & editing.

## References

1 Fueyo, R., Judd, J., Feschotte, C. & Wysocka, J. Roles of transposable elements in the regulation of mammalian transcription. Nat Rev Mol Cell Biol 23, 481–497 (2022). 10.1038/s41580-022-00457-y

2 Hoyt, S. J. et al. From telomere to telomere: The transcriptional and epigenetic state of human repeat elements. Science 376, eabk3112 (2022). 10.1126/science.abk3112

3 Bourque, G. et al. Ten things you should know about transposable elements. Genome Biol 19, 199 (2018). 10.1186/s13059-018-1577-z

4 Senft, A. D. & Macfarlan, T. S. Transposable elements shape the evolution of mammalian development. Nature reviews. Genetics 22, 691–711 (2021). 10.1038/s41576-021-00385-1

5 Erwin, J. A., Marchetto, M. C. & Gage, F. H. Mobile DNA elements in the generation of diversity and complexity in the brain. Nat Rev Neurosci 15, 497–506 (2014). 10.1038/nrn3730

6 Faulkner, G. J. & Garcia-Perez, J. L. L1 Mosaicism in Mammals: Extent, Effects, and Evolution. Trends Genet 33, 802–816 (2017). 10.1016/j.tig.2017.07.004

7 Chuong, E. B., Elde, N. C. & Feschotte, C. Regulatory activities of transposable elements: from conflicts to benefits. Nature reviews. Genetics 18, 71–86 (2017). 10.1038/nrg.2016.139

8 Deniz, O., Frost, J. M. & Branco, M. R. Regulation of transposable elements by DNA modifications. Nature reviews. Genetics 20, 417–431 (2019). 10.1038/s41576-019-0106-6

9 Deleris, A., Berger, F. & Duharcourt, S. Role of Polycomb in the control of transposable elements. Trends Genet 37, 882–889 (2021). 10.1016/j.tig.2021.06.003

10 Shalgi, R., Pilpel, Y. & Oren, M. Repression of transposable-elements -a microRNA anti-cancer defense mechanism? Trends Genet 26, 253–259 (2010). 10.1016/j.tig.2010.03.006

11 Huang, T. C. et al. Sex-specific chromatin remodelling safeguards transcription in germ cells. Nature 600, 737–742 (2021). 10.1038/s41586-021-04208-5

12 Walter, M., Teissandier, A., Perez-Palacios, R. & Bourc’his, D. An epigenetic switch ensures transposon repression upon dynamic loss of DNA methylation in embryonic stem cells. Elife 5 (2016). 10.7554/eLife.11418

13 Burns, K. H. Transposable elements in cancer. Nature reviews. Cancer 17, 415–424 (2017). 10.1038/nrc.2017.35

14 Jang, H. S. et al. Transposable elements drive widespread expression of oncogenes in human cancers. Nat Genet 51, 611–617 (2019). 10.1038/s41588-019-0373-3

15 Thompson, P. J., Macfarlan, T. S. & Lorincz, M. C. Long Terminal Repeats: From Parasitic Elements to Building Blocks of the Transcriptional Regulatory Repertoire. Molecular cell 62, 766–776 (2016). 10.1016/j.molcel.2016.03.029

16 Haws, S. A., Simandi, Z., Barnett, R. J. & Phillips-Cremins, J. E. 3D genome, on repeat: Higher-order folding principles of the heterochromatinized repetitive genome. Cell 185, 2690–2707 (2022). 10.1016/j.cell.2022.06.052

17 Rowley, M. J. & Corces, V. G. Organizational principles of 3D genome architecture. Nature Reviews Genetics 19, 789–800 (2018). 10.1038/s41576-018-0060-8

18 Kempfer, R. & Pombo, A. Methods for mapping 3D chromosome architecture. Nature reviews. Genetics 21, 207–226 (2020). 10.1038/s41576-019-0195-2

19 An, L. et al. OnTAD: hierarchical domain structure reveals the divergence of activity among TADs and boundaries. Genome Biol 20, 282 (2019). 10.1186/s13059-019-1893-y

20 Choudhary, M. N. K., Quaid, K., Xing, X., Schmidt, H. & Wang, T. Widespread contribution of transposable elements to the rewiring of mammalian 3D genomes. Nat Commun 14, 634 (2023). 10.1038/s41467-023-36364-9

21 Diehl, A. G., Ouyang, N. & Boyle, A. P. Transposable elements contribute to cell and species-specific chromatin looping and gene regulation in mammalian genomes. Nat Commun 11, 1796 (2020). 10.1038/s41467-020-15520-5

22 Schwarzer, W. et al. Two independent modes of chromatin organization revealed by cohesin removal. Nature 551, 51–56 (2017). 10.1038/nature24281

23 Kline, A. D. et al. Diagnosis and management of Cornelia de Lange syndrome: first international consensus statement. Nature reviews. Genetics 19, 649–666 (2018). 10.1038/s41576-018-0031-0

24 Bailey, M. H. et al. Comprehensive Characterization of Cancer Driver Genes and Mutations. Cell 173, 371–385 e318 (2018). 10.1016/j.cell.2018.02.060

25 Adiconis, X. et al. Comprehensive comparative analysis of 5’-end RNA-sequencing methods. Nature methods 15, 505–511 (2018). 10.1038/s41592-018-0014-2

26 Wiesner, T. et al. Alternative transcription initiation leads to expression of a novel ALK isoform in cancer. Nature 526, 453-+ (2015). 10.1038/nature15258

27 Gilbert, L. A. et al. CRISPR-mediated modular RNA-guided regulation of transcription in eukaryotes. Cell 154, 442–451 (2013). 10.1016/j.cell.2013.06.044

28 Consortium, F. et al. A promoter-level mammalian expression atlas. Nature 507, 462–470 (2014). 10.1038/nature13182

29 Carninci, P. et al. Genome-wide analysis of mammalian promoter architecture and evolution. Nat Genet 38, 626–635 (2006). 10.1038/ng1789

30 Bergman, Y. & Cedar, H. DNA methylation dynamics in health and disease. Nat Struct Mol Biol 20, 274–281 (2013). 10.1038/nsmb.2518

31 Anastasiadi, D., Esteve-Codina, A. & Piferrer, F. Consistent inverse correlation between DNA methylation of the first intron and gene expression across tissues and species. Epigenetics Chromatin 11, 37 (2018). 10.1186/s13072-018-0205-1

32 Skene, P. J. & Henikoff, S. An efficient targeted nuclease strategy for high-resolution mapping of DNA binding sites. Elife 6 (2017). 10.7554/eLife.21856

33 Ernst, J. & Kellis, M. Chromatin-state discovery and genome annotation with ChromHMM. Nature protocols 12, 2478–2492 (2017). 10.1038/nprot.2017.124

34 Oh, S. et al. Enhancer release and retargeting activates disease-susceptibility genes. Nature 595, 735–740 (2021). 10.1038/s41586-021-03577-1

35 Goding, C. R. & Arnheiter, H. MITF-the first 25 years. Genes Dev 33, 983–1007 (2019). 10.1101/gad.324657.119

36 Rao, S. S. P. et al. Cohesin Loss Eliminates All Loop Domains. Cell 171, 305–320 e324 (2017). 10.1016/j.cell.2017.09.026

37 Dixon, J. R., Gorkin, D. U. & Ren, B. Chromatin Domains: The Unit of Chromosome Organization. Molecular cell 62, 668–680 (2016). 10.1016/j.molcel.2016.05.018

38 Hsieh, T. S. et al. Resolving the 3D Landscape of Transcription-Linked Mammalian Chromatin Folding. Molecular cell 78, 539–553 e538 (2020). 10.1016/j.molcel.2020.03.002

39 Beagan, J. A. & Phillips-Cremins, J. E. On the existence and functionality of topologically associating domains. Nat Genet 52, 8–16 (2020). 10.1038/s41588-019-0561-1

40 Crane, E. et al. Condensin-driven remodelling of X chromosome topology during dosage compensation. Nature 523, 240–244 (2015). 10.1038/nature14450

41 de Wit, E. & Nora, E. P. New insights into genome folding by loop extrusion from inducible degron technologies. Nature reviews. Genetics (2022). 10.1038/s41576-022-00530-4

42 Dixon, J. R. et al. Topological domains in mammalian genomes identified by analysis of chromatin interactions. Nature 485, 376–380 (2012). 10.1038/nature11082

43 Krietenstein, N. et al. Ultrastructural Details of Mammalian Chromosome Architecture. Molecular cell 78, 554–565 e557 (2020). 10.1016/j.molcel.2020.03.003

44 Tarjan, D. R., Flavahan, W. A. & Bernstein, B. E. Epigenome editing strategies for the functional annotation of CTCF insulators. Nat Commun 10, 4258 (2019). 10.1038/s41467-019-12166-w

45 Krijger, P. H. L., Geeven, G., Bianchi, V., Hilvering, C. R. E. & de Laat, W. 4C-seq from beginning to end: A detailed protocol for sample preparation and data analysis. Methods 170, 17–32 (2020). 10.1016/j.ymeth.2019.07.014

46 Fu, Y. et al. FunSeq2: a framework for prioritizing noncoding regulatory variants in cancer. Genome Biol 15, 480 (2014). 10.1186/s13059-014-0480-5

47 Demircioglu, D. et al. A Pan-cancer Transcriptome Analysis Reveals Pervasive Regulation through Alternative Promoters. Cell 178, 1465–1477 e1417 (2019). 10.1016/j.cell.2019.08.018

48 Hermant, C. & Torres-Padilla, M. E. TFs for TEs: the transcription factor repertoire of mammalian transposable elements. Genes Dev 35, 22–39 (2021). 10.1101/gad.344473.120

49 Ito, J. et al. Systematic identification and characterization of regulatory elements derived from human endogenous retroviruses. Plos Genet 13, e1006883 (2017). 10.1371/journal.pgen.1006883

50 Sundaram, V. et al. Widespread contribution of transposable elements to the innovation of gene regulatory networks. Genome research 24, 1963–1976 (2014). 10.1101/gr.168872.113

51 Tome, J. M., Tippens, N. D. & Lis, J. T. Single-molecule nascent RNA sequencing identifies regulatory domain architecture at promoters and enhancers. Nat Genet 50, 1533–1541 (2018). 10.1038/s41588-018-0234-5

52 Core, L. J., Martins, A. L., Danko, C. G., Waters, C. T., Siepel, A. & Lis, J. T. Analysis of nascent RNA identifies a unified architecture of initiation regions at mammalian promoters and enhancers. Nat Genet 46, 1311–1320 (2014). 10.1038/ng.3142

53 Newkirk, D. A. et al. The effect of Nipped-B-like (Nipbl) haploinsufficiency on genome-wide cohesin binding and target gene expression: modeling Cornelia de Lange syndrome. Clin Epigenetics 9, 89 (2017). 10.1186/s13148-017-0391-x

54 Goel, V. Y., Huseyin, M. K. & Hansen, A. S. Region Capture Micro-C reveals coalescence of enhancers and promoters into nested microcompartments. bioRxiv, 2022.2007. 2012.499637 (2022).

55 Lupianez, D. G. et al. Disruptions of topological chromatin domains cause pathogenic rewiring of gene-enhancer interactions. Cell 161, 1012–1025 (2015). 10.1016/j.cell.2015.04.004

56 Rinzema, N. J. et al. Building regulatory landscapes reveals that an enhancer can recruit cohesin to create contact domains, engage CTCF sites and activate distant genes. Nat Struct Mol Biol 29, 563–574 (2022). 10.1038/s41594-022-00787-7

57 Xu, C., Park, J. K. & Zhang, J. Evidence that alternative transcriptional initiation is largely nonadaptive. PLoS Biol 17, e3000197 (2019). 10.1371/journal.pbio.3000197

58 Davuluri, R. V., Suzuki, Y., Sugano, S., Plass, C. & Huang, T. H. The functional consequences of alternative promoter use in mammalian genomes. Trends Genet 24, 167–177 (2008). 10.1016/j.tig.2008.01.008

59 Huang, K. K. et al. Long-read transcriptome sequencing reveals abundant promoter diversity in distinct molecular subtypes of gastric cancer. Genome Biol 22, 44 (2021). 10.1186/s13059-021-02261-x

60 Schumacher, T. N. & Schreiber, R. D. Neoantigens in cancer immunotherapy. Science 348, 69–74 (2015). 10.1126/science.aaa4971

61 Pearlman, A. H. et al. Targeting public neoantigens for cancer immunotherapy. Nat Cancer 2, 487–497 (2021). 10.1038/s43018-021-00210-y

62 Wong, E. W., Lee, W. M. & Cheng, C. Y. Secreted Frizzled-related protein 1 (sFRP1) regulates spermatid adhesion in the testis via dephosphorylation of focal adhesion kinase and the nectin-3 adhesion protein complex. FASEB J 27, 464–477 (2013). 10.1096/fj.12-212514

63 Concordet, J. P. & Haeussler, M. CRISPOR: intuitive guide selection for CRISPR/Cas9 genome editing experiments and screens. Nucleic Acids Res 46, W242–W245 (2018). 10.1093/nar/gky354

64 Chen, B. et al. Dynamic imaging of genomic loci in living human cells by an optimized CRISPR/Cas system. Cell 155, 1479–1491 (2013). 10.1016/j.cell.2013.12.001

65 Takahashi, H., Lassmann, T., Murata, M. & Carninci, P. 5’ end-centered expression profiling using cap-analysis gene expression and next-generation sequencing. Nature protocols 7, 542–561 (2012). 10.1038/nprot.2012.005

66 Chi, P. et al. ETV1 is a lineage survival factor that cooperates with KIT in gastrointestinal stromal tumours. Nature 467, 849–853 (2010). 10.1038/nature09409

67 Mumbach, M. R. et al. HiChIP: efficient and sensitive analysis of protein-directed genome architecture. Nature methods 13, 919–922 (2016). 10.1038/nmeth.3999

68 Ewels, P. A. et al. The nf-core framework for community-curated bioinformatics pipelines. Nat Biotechnol 38, 276–278 (2020). 10.1038/s41587-020-0439-x

69 Frith, M. C., Valen, E., Krogh, A., Hayashizaki, Y., Carninci, P. & Sandelin, A. A code for transcription initiation in mammalian genomes. Genome research 18, 1–12 (2008). 10.1101/gr.6831208

70 Xie, Y. et al. SETD2 loss perturbs the kidney cancer epigenetic landscape to promote metastasis and engenders actionable dependencies on histone chaperone complexes. Nat Cancer 3, 188–202 (2022). 10.1038/s43018-021-00316-3

71 Krueger, F. & Andrews, S. R. Bismark: a flexible aligner and methylation caller for Bisulfite-Seq applications. Bioinformatics 27, 1571–1572 (2011). 10.1093/bioinformatics/btr167

72 Hahne, F. & Ivanek, R. Visualizing Genomic Data Using Gviz and Bioconductor. Methods Mol Biol 1418, 335–351 (2016). 10.1007/978-1-4939-3578-9_16

73 Sahin, M., Wong, W., Zhan, Y., Van Deynze, K., Koche, R. & Leslie, C. S. HiC-DC+ enables systematic 3D interaction calls and differential analysis for Hi-C and HiChIP. Nat Commun 12, 3366 (2021). 10.1038/s41467-021-23749-x

