## Supplementary table 4 for "TAD hierarchy restricts poised LTR activation and loss of TAD hierarchy promotes LTR co-option in cancer"

### Supplementary Table 4. Homer motif analysis at differentially regulated CTSSs at non-promoter regions.

#### Homer *de novo* Motif Results (shNIPBL_#1_CTSS_non-promoter)

* - possible false positive

| Rank | Motif | P-value | log P-pvalue | % of Targets | % of Background | STD(Bg STD) | Best Match/Details | Motif File |
|---|---|---|---|---|---|---|---|---|
| 1 | 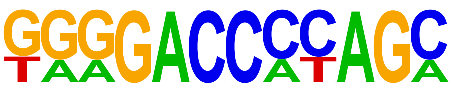 | 1e-21 | -5.055e+01 | 7.96% | 0.15% | 39.1bp (41.9bp) | LRF(Zf)/Erythroblasts-ZBTB7A-ChIP-Seq(GSE74977)/Homer(0.706) More Information \| Similar Motifs Found | motif file (matrix) |
| 2 | 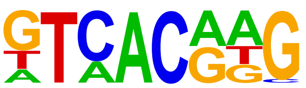 | 1e-19 | -4.396e+01 | 44.28% | 16.80% | 55.1bp (80.8bp) | MITF(bHLH)/MastCells-MITF-ChIP-Seq(GSE48085)/Homer(0.881) More Information \| Similar Motifs Found | motif file (matrix) |
| 3 | 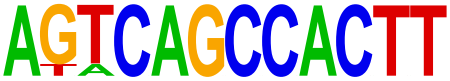 | 1e-18 | -4.328e+01 | 3.48% | 0.01% | 26.3bp (23.3bp) | PH0115.1_Nkx2-6/Jaspar(0.700) More Information \| Similar Motifs Found | motif file (matrix) |
| 4 | 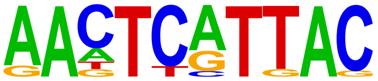 | 1e-17 | -4.045e+01 | 9.45% | 0.52% | 39.3bp (85.4bp) | POU6F1(var.2)/MA1549.1/Jaspar(0.780) More Information \| Similar Motifs Found | motif file (matrix) |
| 5 | 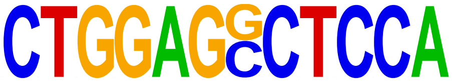 | 1e-16 | -3.843e+01 | 3.48% | 0.01% | 35.4bp (41.3bp) | ZBTB6/MA1581.1/Jaspar(0.658) More Information \| Similar Motifs Found | motif file (matrix) |
| 6 | 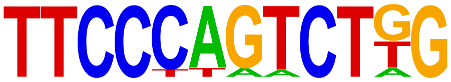 | 1e-16 | -3.726e+01 | 5.97% | 0.12% | 51.6bp (72.4bp) | RBPJ/MA1116.1/Jaspar(0.720) More Information \| Similar Motifs Found | motif file (matrix) |
| 7 | 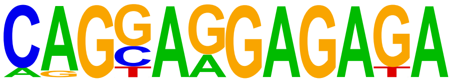 | 1e-15 | -3.638e+01 | 9.45% | 0.65% | 34.2bp (55.9bp) | ZKSCAN5/MA1652.1/Jaspar(0.719) More Information \| Similar Motifs Found | motif file (matrix) |
| 8 | 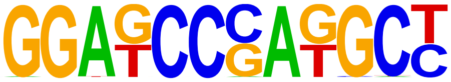 | 1e-15 | -3.539e+01 | 5.97% | 0.15% | 59.4bp (36.7bp) | POL013.1_MED-1/Jaspar(0.620) More Information \| Similar Motifs Found | motif file (matrix) |
| 9 | 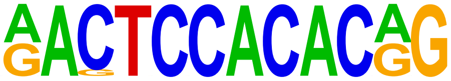 | 1e-14 | -3.359e+01 | 3.48% | 0.02% | 41.0bp (36.2bp) | FOXH1/MA0479.1/Jaspar(0.698) More Information \| Similar Motifs Found | motif file (matrix) |
| 10 | 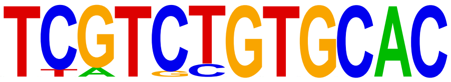 | 1e-13 | -3.036e+01 | 4.98% | 0.12% | 31.1bp (47.7bp) | PB0026.1_Gm397_1/Jaspar(0.770) More Information \| Similar Motifs Found | motif file (matrix) |
| 11 | 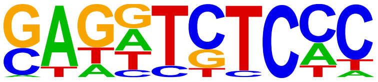 | 1e-13 | -3.030e+01 | 35.82% | 14.59% | 57.0bp (85.9bp) | ZNF274/MA1592.1/Jaspar(0.683) More Information \| Similar Motifs Found | motif file (matrix) |
| 12 | 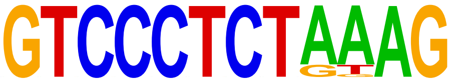 | 1e-12 | -2.977e+01 | 2.99% | 0.01% | 33.5bp (11.7bp) | BCL6/MA0463.2/Jaspar(0.642) More Information \| Similar Motifs Found | motif file (matrix) |
| 13 | 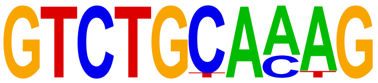 | 1e-12 | -2.966e+01 | 5.97% | 0.24% | 49.8bp (72.3bp) | Smad4(MAD)/ESC-SMAD4-ChIP-Seq(GSE29422)/Homer(0.718) More Information \| Similar Motifs Found | motif file (matrix) |
| 14 | 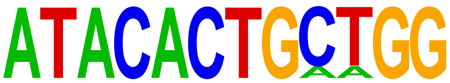 | 1e-12 | -2.959e+01 | 2.49% | 0.00% | 34.2bp (0.0bp) | Zic3(Zf)/mES-Zic3-ChIP-Seq(GSE37889)/Homer(0.606) More Information \| Similar Motifs Found | motif file (matrix) |
| 15 | 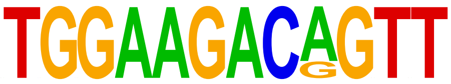 | 1e-12 | -2.959e+01 | 2.49% | 0.00% | 9.2bp (16.6bp) | ZNF528(Zf)/HEK293-ZNF528.GFP-ChIP-Seq(GSE58341)/Homer(0.684) More Information \| Similar Motifs Found | motif file (matrix) |
| 16 | 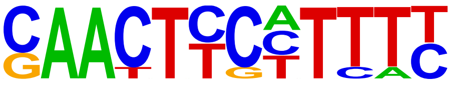 | 1e-12 | -2.880e+01 | 6.47% | 0.33% | 66.7bp (60.9bp) | ELF3(ETS)/PDAC-ELF3-ChIP-Seq(GSE64557)/Homer(0.652) More Information \| Similar Motifs Found | motif file (matrix) |
| 17 | 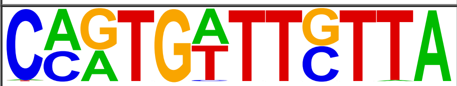 | 1e-12 | -2.773e+01 | 5.47% | 0.21% | 52.7bp (75.6bp) | PB0172.1_Sox1_2/Jaspar(0.708) More Information \| Similar Motifs | motif file |

| | | | | | | | | |
|---|---|---|---|---|---|---|---|---|
| | | | | | | | Found | (matrix) |
| 18 * | 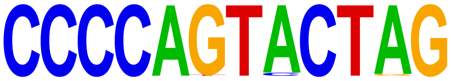 | 1e-11 | -2.721e+01 | 3.48% | 0.04% | 30.2bp (51.8bp) | PB0152.1_Nkx3-1_2/Jaspar(0.591) More Information \| Similar Motifs Found | motif file (matrix) |
| 19 * | 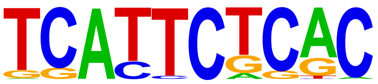 | 1e-11 | -2.721e+01 | 9.95% | 1.24% | 37.1bp (56.9bp) | Eomes(T-box)/H9-Eomes-ChIP-Seq(GSE26097)/Homer(0.645) More Information \| Similar Motifs Found | motif file (matrix) |
| 20 * | 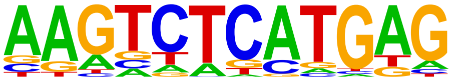 | 1e-11 | -2.715e+01 | 26.87% | 9.53% | 48.4bp (81.5bp) | USF2/MA0526.3/Jaspar(0.660) More Information \| Similar Motifs Found | motif file (matrix) |
| 21 * | 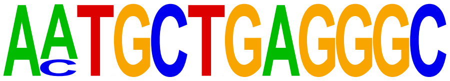 | 1e-11 | -2.613e+01 | 2.49% | 0.01% | 76.6bp (26.3bp) | Mafb/MA0117.2/Jaspar(0.684) More Information \| Similar Motifs Found | motif file (matrix) |
| 22 * | 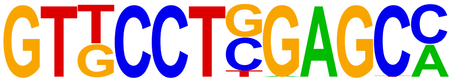 | 1e-11 | -2.581e+01 | 4.98% | 0.18% | 73.9bp (101.0bp) | ZBTB6/MA1581.1/Jaspar(0.774) More Information \| Similar Motifs Found | motif file (matrix) |
| 23 * | 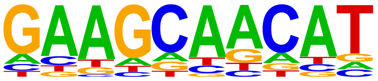 | 1e-10 | -2.464e+01 | 8.96% | 1.11% | 54.2bp (85.3bp) | POL008.1_DCE_S_I/Jaspar(0.690) More Information \| Similar Motifs Found | motif file (matrix) |
| 24 * | 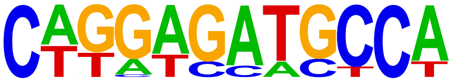 | 1e-10 | -2.322e+01 | 2.99% | 0.03% | 62.7bp (79.3bp) | Hic1/MA0739.1/Jaspar(0.634) More Information \| Similar Motifs Found | motif file (matrix) |
| 25 * | 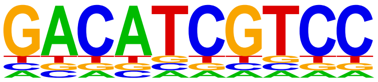 | 1e-9 | -2.301e+01 | 1.99% | 0.01% | 36.8bp (18.2bp) | ZNF669(Zf)/HEK293-ZNF669.GFP-ChIP-Seq(GSE58341)/Homer(0.658) More Information \| Similar Motifs Found | motif file (matrix) |
| 26 * | 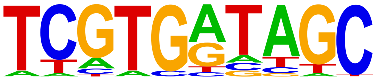 | 1e-9 | -2.285e+01 | 18.41% | 5.52% | 54.0bp (76.5bp) | Npas4(bHLH)/Neuron-Npas4-ChIP-Seq(GSE127793)/Homer(0.723) More Information \| Similar Motifs Found | motif file (matrix) |
| 27 * | 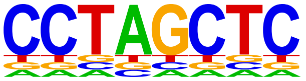 | 1e-9 | -2.227e+01 | 5.97% | 0.45% | 35.9bp (68.2bp) | POL010.1_DCE_S_III/Jaspar(0.614) More Information \| Similar Motifs Found | motif file (matrix) |
| 28 * | 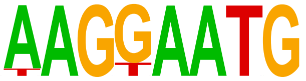 | 1e-8 | -2.040e+01 | 11.94% | 2.68% | 49.4bp (80.3bp) | TEAD(TEA)/Fibroblast-PU.1-ChIP-Seq(Unpublished)/Homer(0.831) More Information \| Similar Motifs Found | motif file (matrix) |
| 29 * | 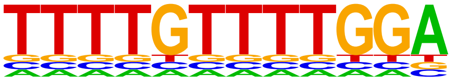 | 1e-8 | -2.024e+01 | 1.99% | 0.01% | 44.1bp (16.5bp) | PB0123.1_Foxl1_2/Jaspar(0.771) More Information \| Similar Motifs Found | motif file (matrix) |
| 30 * | 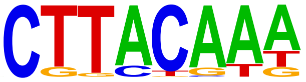 | 1e-8 | -1.954e+01 | 18.41% | 6.24% | 46.7bp (80.4bp) | Sox21(HMG)/ESC-SOX21-ChIP-Seq(GSE110505)/Homer(0.630) More Information \| Similar Motifs Found | motif file (matrix) |
| 31 * | 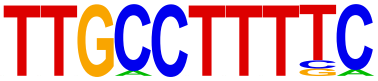 | 1e-8 | -1.924e+01 | 5.47% | 0.47% | 28.4bp (69.2bp) | Nr2e1/MA0676.1/Jaspar(0.774) More Information \| Similar Motifs Found | motif file (matrix) |
| 32 * | 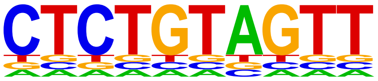 | 1e-7 | -1.813e+01 | 2.49% | 0.04% | 26.6bp (26.2bp) | RUNX-AML(Runt)/CD4+-PolII-ChIP-Seq(Barski_et_al.)/Homer(0.686) More Information \| Similar Motifs Found | motif file (matrix) |
| 33 * | 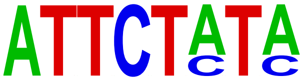 | 1e-7 | -1.749e+01 | 11.94% | 3.12% | 43.9bp (75.0bp) | PB0068.1_Sox1_1/Jaspar(0.679) More Information \| Similar Motifs Found | motif file (matrix) |
| 34 * | 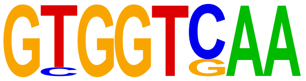 | 1e-7 | -1.742e+01 | 9.45% | 1.98% | 59.5bp (80.4bp) | PB0030.1_Hnf4a_1/Jaspar(0.797) More Information \| Similar Motifs Found | motif file (matrix) |
| 35 * | 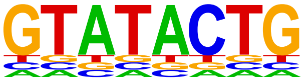 | 1e-6 | -1.580e+01 | 3.48% | 0.19% | 32.9bp (67.0bp) | ZBTB32/MA1580.1/Jaspar(0.628) More Information \| Similar Motifs Found | motif file (matrix) |
| 36 * | 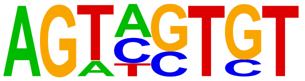 | 1e-5 | -1.368e+01 | 8.46% | 2.05% | 44.3bp (85.6bp) | PB0141.1_Isgf3g_2/Jaspar(0.686) More Information \| Similar Motifs Found | motif file (matrix) |

### Supplementary Table 4. Homer motif analysis at differentially regulated CTSSs at non-promoter regions.

#### Homer *de novo* Motif Results (shNIPBL_#2_CTSS_non-promoter)

* - possible false positive

| Rank | Motif | P-value | log P-pvalue | % of Targets | % of Background | STD(Bg STD) | Best Match/Details | Motif File |
|---|---|---|---|---|---|---|---|---|
| 1 | 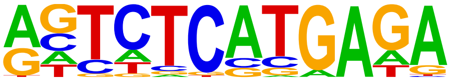 | 1e-41 | -9.660e+01 | 23.82% | 7.53% | 53.8bp (74.2bp) | GFX(?)/Promoter/Homer(0.748) More Information \| Similar Motifs Found | motif file (matrix) |
| 2 | 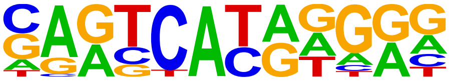 | 1e-41 | -9.659e+01 | 30.45% | 11.63% | 63.9bp (107.9bp) | MITF(bHLH)/MastCells-MITF-ChIP-Seq(GSE48085)/Homer(0.748) More Information \| Similar Motifs Found | motif file (matrix) |
| 3 | 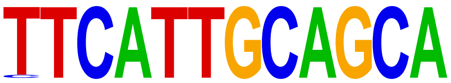 | 1e-38 | -8.965e+01 | 2.30% | 0.01% | 14.9bp (0.0bp) | Hoxa10(Homeobox)/ChickenMSG-Hoxa10.Flag-ChIP-Seq(GSE86088)/Homer(0.706) More Information \| Similar Motifs Found | motif file (matrix) |
| 4 | 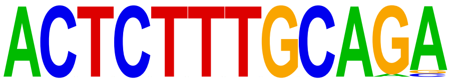 | 1e-38 | -8.965e+01 | 2.30% | 0.01% | 42.5bp (4.3bp) | NFIC/MA0161.2/Jaspar(0.630) More Information \| Similar Motifs Found | motif file (matrix) |
| 5 | 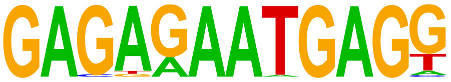 | 1e-36 | -8.421e+01 | 7.31% | 0.68% | 23.7bp (52.7bp) | PH0152.1_Pou6f1_2/Jaspar(0.675) More Information \| Similar Motifs Found | motif file (matrix) |
| 6 | 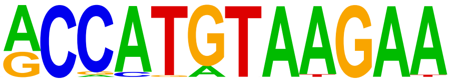 | 1e-36 | -8.323e+01 | 7.17% | 0.65% | 34.1bp (50.0bp) | PH0085.1_Irx4/Jaspar(0.599) More Information \| Similar Motifs Found | motif file (matrix) |
| 7 | 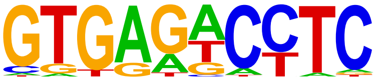 | 1e-35 | -8.204e+01 | 21.65% | 7.15% | 61.2bp (90.0bp) | ZNF274/MA1592.1/Jaspar(0.739) More Information \| Similar Motifs Found | motif file (matrix) |
| 8 | 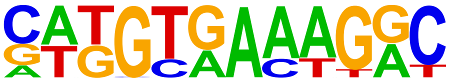 | 1e-34 | -7.896e+01 | 5.28% | 0.30% | 43.8bp (67.0bp) | Tbox:Smad(T-box,MAD)/ESCd5-Smad2_3-ChIP-Seq(GSE29422)/Homer(0.683) More Information \| Similar Motifs Found | motif file (matrix) |
| 9 | 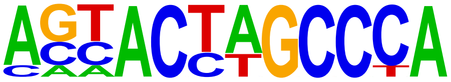 | 1e-33 | -7.704e+01 | 2.17% | 0.01% | 32.5bp (13.1bp) | SD0002.1_at_AC_acceptor/Jaspar(0.651) More Information \| Similar Motifs Found | motif file (matrix) |
| 10 | 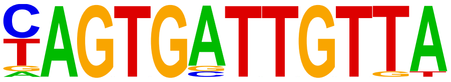 | 1e-33 | -7.615e+01 | 2.44% | 0.02% | 40.7bp (58.7bp) | Sox5/MA0087.1/Jaspar(0.732) More Information \| Similar Motifs Found | motif file (matrix) |
| 11 | 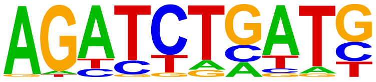 | 1e-32 | -7.445e+01 | 25.58% | 10.12% | 48.5bp (79.5bp) | GATA(Zf),IR3/iTreg-Gata3-ChIP-Seq(GSE20898)/Homer(0.691) More Information \| Similar Motifs Found | motif file (matrix) |
| 12 | 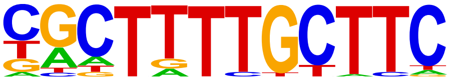 | 1e-32 | -7.425e+01 | 10.55% | 1.94% | 39.2bp (72.6bp) | IRF2(IRF)/Erythroblas-IRF2-ChIP-Seq(GSE36985)/Homer(0.677) More Information \| Similar Motifs Found | motif file (matrix) |
| 13 | 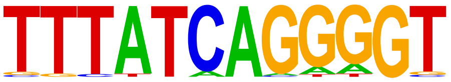 | 1e-31 | -7.325e+01 | 6.90% | 0.73% | 32.0bp (40.4bp) | HOXB13(Homeobox)/ProstateTumor-HOXB13-ChIP-Seq(GSE56288)/Homer(0.685) More Information \| Similar Motifs Found | motif file (matrix) |
| 14 | 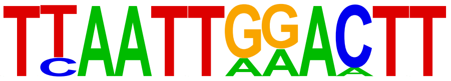 | 1e-28 | -6.596e+01 | 4.87% | 0.34% | 74.6bp (46.0bp) | ALX3/MA0634.1/Jaspar(0.733) More Information \| Similar Motifs Found | motif file (matrix) |
| 15 | 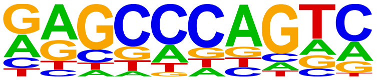 | 1e-24 | -5.687e+01 | 4.33% | 0.32% | 51.5bp (62.1bp) | ZNF416(Zf)/HEK293-ZNF416.GFP-ChIP-Seq(GSE58341)/Homer(0.784) More Information \| Similar Motifs Found | motif file (matrix) |
| 16 | 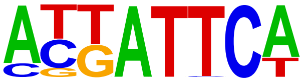 | 1e-24 | -5.572e+01 | 21.24% | 8.81% | 60.6bp (94.4bp) | Brn2(POU,Homeobox)/NPC-Brn2-ChIP-Seq(GSE35496)/Homer(0.771) More Information \| Similar Motifs Found | motif file (matrix) |
| 17 | 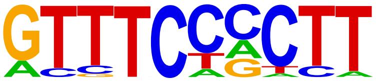 | 1e-23 | -5.500e+01 | 12.86% | 3.80% | 44.7bp (76.5bp) | ISRE(IRF)/ThioMac-LPS-Expression(GSE23622)/Homer(0.707) More Information \| Similar Motifs Found | motif file (matrix) |
| 18 | 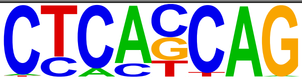 | 1e-23 | -5.450e+01 | 27.06% | 13.00% | 57.3bp (86.2bp) | Unknown-ESC-element(?)/mES-Nanog-ChIP-Seq(GSE11724)/Homer(0.768) | motif file |

| | | | | | | | | |
|---|---|---|---|---|---|---|---|---|
| | | | | | | | More Information \| Similar Motifs Found | (matrix) |
| 19 | 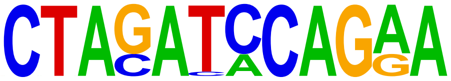 | 1e-23 | -5.342e+01 | 2.03% | 0.03% | 19.7bp (56.3bp) | ZBTB26/MA1579.1/Jaspar(0.678) More Information \| Similar Motifs Found | motif file (matrix) |
| 20 | 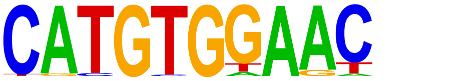 | 1e-22 | -5.191e+01 | 9.20% | 2.13% | 67.0bp (71.8bp) | MXI1/MA1108.2/Jaspar(0.807) More Information \| Similar Motifs Found | motif file (matrix) |
| 21 | 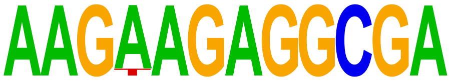 | 1e-21 | -4.871e+01 | 1.22% | 0.00% | 42.0bp (14.6bp) | SPI1/MA0080.5/Jaspar(0.580) More Information \| Similar Motifs Found | motif file (matrix) |
| 22 | 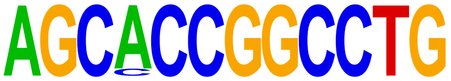 | 1e-21 | -4.871e+01 | 1.22% | 0.00% | 32.0bp (5.9bp) | Zfx/MA0146.2/Jaspar(0.691) More Information \| Similar Motifs Found | motif file (matrix) |
| 23 | 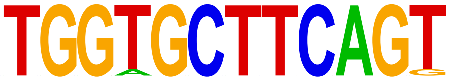 | 1e-20 | -4.810e+01 | 1.35% | 0.01% | 51.9bp (20.2bp) | OSR2/MA1646.1/Jaspar(0.658) More Information \| Similar Motifs Found | motif file (matrix) |
| 24 | 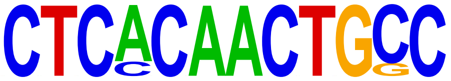 | 1e-20 | -4.810e+01 | 1.35% | 0.01% | 15.9bp (3.5bp) | MYB/MA0100.3/Jaspar(0.744) More Information \| Similar Motifs Found | motif file (matrix) |
| 25 | 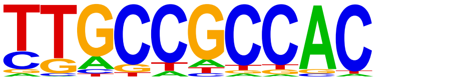 | 1e-19 | -4.497e+01 | 6.77% | 1.33% | 28.7bp (49.7bp) | YY2/MA0748.2/Jaspar(0.706) More Information \| Similar Motifs Found | motif file (matrix) |
| 26 | 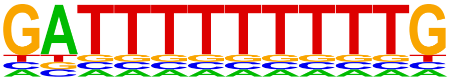 | 1e-19 | -4.406e+01 | 1.35% | 0.01% | 22.3bp (34.3bp) | ZNF384/MA1125.1/Jaspar(0.790) More Information \| Similar Motifs Found | motif file (matrix) |
| 27 | 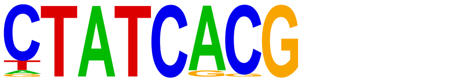 | 1e-19 | -4.398e+01 | 9.61% | 2.68% | 44.5bp (67.5bp) | Npas4(bHLH)/Neuron-Npas4-ChIP-Seq(GSE127793)/Homer(0.712) More Information \| Similar Motifs Found | motif file (matrix) |
| 28 | 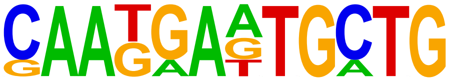 | 1e-18 | -4.249e+01 | 1.22% | 0.01% | 24.6bp (11.2bp) | MAFG/MA0659.2/Jaspar(0.678) More Information \| Similar Motifs Found | motif file (matrix) |
| 29 | 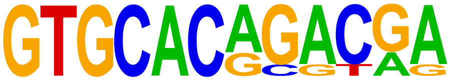 | 1e-17 | -4.131e+01 | 1.76% | 0.03% | 25.4bp (23.8bp) | PB0026.1_Gm397_1/Jaspar(0.789) More Information \| Similar Motifs Found | motif file (matrix) |
| 30 | 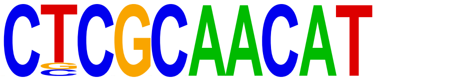 | 1e-17 | -4.120e+01 | 1.35% | 0.01% | 51.0bp (34.3bp) | CEBP:AP1(bZIP)/ThioMac-CEBPb-ChIP-Seq(GSE21512)/Homer(0.708) More Information \| Similar Motifs Found | motif file (matrix) |
| 31 | 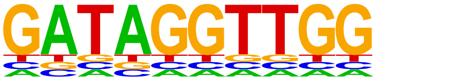 | 1e-16 | -3.699e+01 | 1.08% | 0.01% | 46.6bp (0.0bp) | PB0029.1_Hic1_1/Jaspar(0.664) More Information \| Similar Motifs Found | motif file (matrix) |
| 32 | 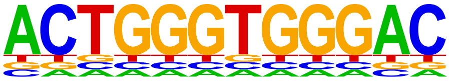 | 1e-16 | -3.699e+01 | 1.08% | 0.01% | 34.1bp (6.9bp) | PB0167.1_Sox13_2/Jaspar(0.728) More Information \| Similar Motifs Found | motif file (matrix) |
| 33 | 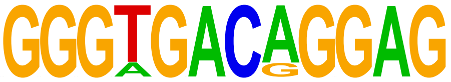 | 1e-16 | -3.699e+01 | 1.08% | 0.01% | 38.9bp (37.2bp) | PBX3/MA1114.1/Jaspar(0.702) More Information \| Similar Motifs Found | motif file (matrix) |
| 34 | 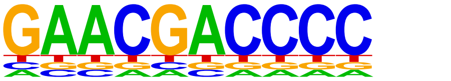 | 1e-15 | -3.645e+01 | 0.95% | 0.00% | 34.5bp (18.6bp) | LRF(Zf)/Erythroblasts-ZBTB7A-ChIP-Seq(GSE74977)/Homer(0.745) More Information \| Similar Motifs Found | motif file (matrix) |
| 35 | 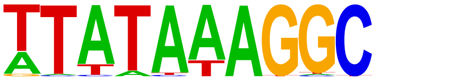 | 1e-15 | -3.480e+01 | 2.44% | 0.16% | 47.3bp (64.3bp) | TATA-Box(TBP)/Promoter/Homer(0.727) More Information \| Similar Motifs Found | motif file (matrix) |
| 36 | 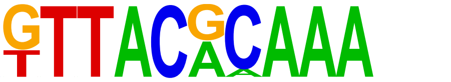 | 1e-14 | -3.402e+01 | 2.44% | 0.17% | 43.9bp (70.9bp) | CEBPB/MA0466.2/Jaspar(0.755) More Information \| Similar Motifs Found | motif file (matrix) |
| 37 * | 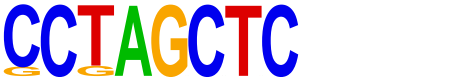 | 1e-8 | -2.071e+01 | 4.19% | 1.13% | 45.3bp (66.2bp) | POL010.1_DCE_S_III/Jaspar(0.639) More Information \| Similar Motifs Found | motif file (matrix) |
| 38 * | 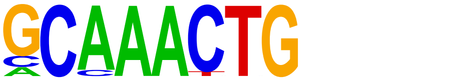 | 1e-8 | -1.857e+01 | 8.80% | 4.06% | 48.5bp (83.6bp) | PB0150.1_Mybl1_2/Jaspar(0.740) More Information \| Similar Motifs Found | motif file (matrix) |
| 39 * | 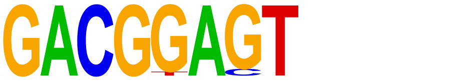 | 1e-7 | -1.622e+01 | 2.84% | 0.69% | 26.6bp (65.6bp) | POL013.1_MED-1/Jaspar(0.622) More Information \| Similar Motifs Found | motif file (matrix) |
